# Exon 9 *LEPR* Gene SNP Polymorphism of Hybrid Chickens F_2_ *Kambro* Crossbreeds of ♀ F_1_ *Kambro* with ♂ F_1_ *Kambro*

**DOI:** 10.1101/2021.02.13.431072

**Authors:** I Wayan Swarautama Mahardhika, Budi Setiadi Daryono

**Author notes:** Departement of Biology, Faculty of Science and Technology, Universitas Airlangga.

## Abstract

The implementation of the T-ARMS PCR method in the detection of single nucleotide polymorphisms (SNPs) in the *LEPR* gene in chicken DNA samples has never been conducted. This research aims to design a specific protocol for exon 9 *LEPR* gene SNPs detection and detect *LEPR* gene expression or *LEPR* SNPs in *Pelung* chicken samples, F_1_ *Pelung*, Layer, Broiler Cobb 500, F_1_ *Kambro* chicken and F_2_ *Kambro* chicken using the T-ARMS PCR method. Determination of *LEPR* gene correlation degree on Body Weight (BT) and Egg Productivity (PT) in F_1_ Kambro population and F_2_ *Kambro*. Qualitative phenotype parameters showed six groups of segregated phenotypes compared to F_1_ *Kambro* chicken. Growth of F_2_ *Kambro* chicken weight reached 753.36 ± 155.31 grams in 8 weeks was not significant for F_1_ *Kambro* chicken due to inbreeding depression (Fx = 25%, IR = 4.925%) and transversion of A *LEPR* allele mutations. Specific protocol detection of exon 9 *LEPR* gene SNPs using the T-ARMS PCR method can detect C127A *LEPR* mutations with IP: OP ratio 10:1 pmol / µM, chicken DNA template concentration of 100 ng / µL with annealing temperature of 55.7° C / 30s. The transversion mutation of C127A of *LEPR* exon 9 SNP were detected in DNA samples of F_1_ *Kambro* hens (80%), F_2_ *Kambro* roosters (20%), Broiler Cobb 500 hens (75%). The mutations were not detected in Layer, *Pelung Blirik Hitam* chicken and F_1_ *Pelung* populations.

## 1. INTRODUCTIONS

Advances in molecular biotechnology enable fast and reliable methods for the accurate diagnosis of mutations responsible for different genetic defects. The Amplification Refractory Mutation System (ARMS)-PCR and tetra-primer PCR detects known sequence polymorphisms (Alyethodi et al., 2016). The combination of aforesaid two technique generated tetra-primer ARMS-PCR or T-ARMS technique. ARMS-PCR has several advantages, namely the need for samples is small, rapid and effective, efficient and simultaneous, the level of sensitivity and accuracy are high and consistent (Peng et al., 2017). The implementation of this method using DNA samples from chicken to detect single nucleotide polymorphism in the *LEPR* gene has never been done. This research was aimed to set the protocol for detecting *LEPR* SNP polymorphism (C127A) at exon 9 in F_2_ Kambro hybrid chickens using Tetra Primer Amplification Refractory Mutation System (T-ARMS) PCR method.

## 2. METHODS

### 2.1. Sample Collection

Hybrid chickens (F_1_ *Pelung, Kambro* and F_2_ *Kambro*) and parental generation (*Pelung Blirik Hitam*, Broiler Cobb 500 and Layer) reared at the chicken rearing unit of Agrotechnology Innovation Center (PIAT) Kali Tirto Berbah Sleman Regency Yogyakarta, Indonesia were used for the study (Table 1.). Phenotypic characters of body weight, egg productivity and phenotype parameters were measured. Genomic DNA was isolated from whole blood by chelex based method (Ernanto et al., 2018) with minor modifications. The isolated DNA purity were quantified with Spark® Reader spectrophotometer (TECAN). The DNA were kept dissolved in TE buffer (pH 8.0) at -20° C until use.

**Table 1.**
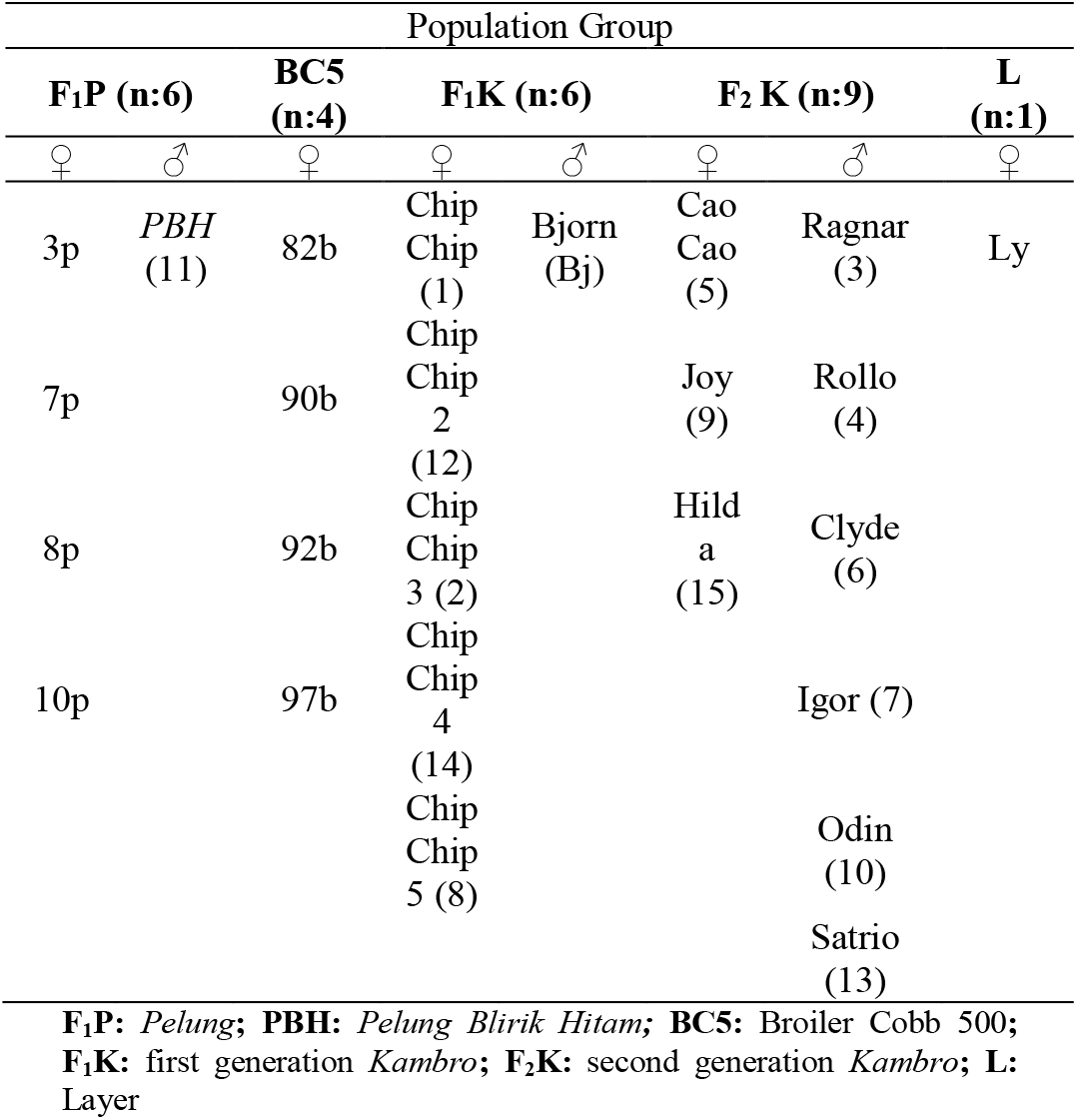
Chicken DNA sample tags.

### 2.2. Tetra ARMS Primer Designs

Primers were designed by the original software on the website: http://primer1.soton.ac.uk (Table 2.). Four primers were designed (FIA: forward inner primer A allele; RIC: reverse inner primer C allele; FOP: forward outer primer; ROP: reverse outer primer) enabling the specific amplification of normal and mutant alleles. The primer specifity was tested using the BLAST program of NCBI. Allele-specific amplicons with different product lengths were separated by standard agarose gel electrophoresis.

**Table 2.**
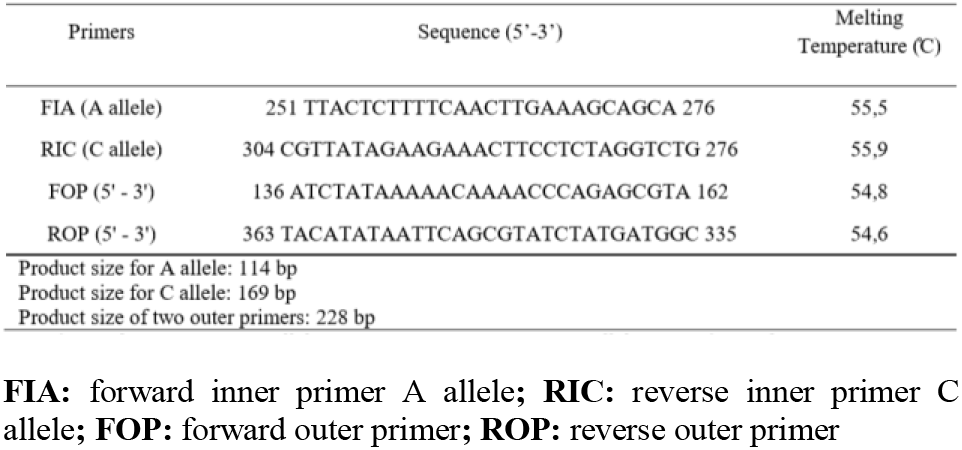
Tetra ARMS PCR primers.

### 2.3. In Vitro Amplification, Visualization and Analysis

The composition of each T-ARMS PCR reaction mix consisted of 50-100 ng of genomic DNA with concentration range from 1-2 (260/280 nm), 200 µM of each dNTP at 1X and 0.5 U per 25 µL reaction, KAPA Taq DNA polymerase with buffer containing MgCl_2_ (1.5 mM at 1X) and stabilizers. The final reaction volume of 25 µL was made with nuclease-free water. Each PCR reaction were amplified with BIORAD T100 ™ PCR Thermal Cycler. A series of experiments were performed to validate the performance of Tetra ARMS PCR to detect chicken *LEPR* gene by changing primers concentration ratio and PCR steps as can be seen in Table 3.

**Table 3.**
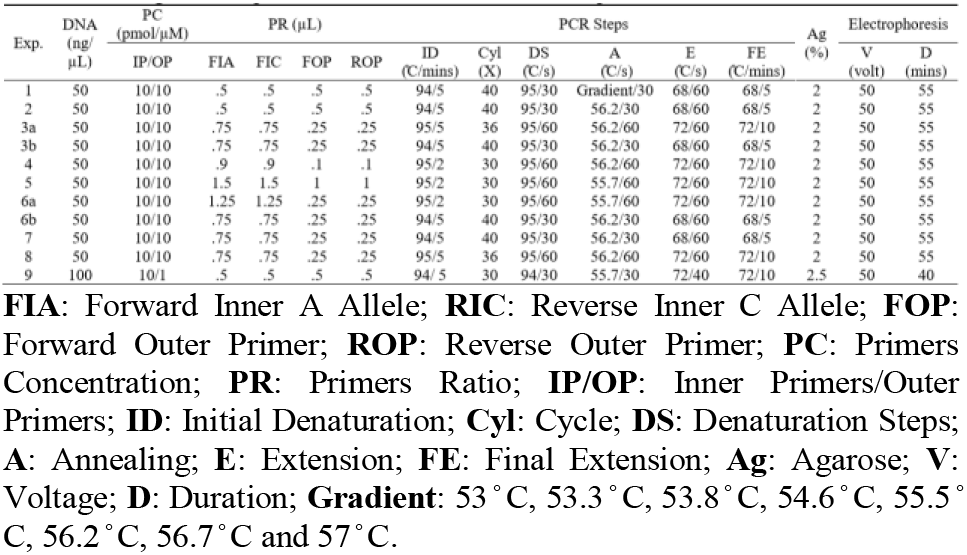
Configuration of primers and PCR steps

The generated amplicons were separated by agarose gel and visualized under UV light by AnalytikJena gel imaging system. The mutant allele and normal allele are differentiated by checking the amplicon sizes in reference with size markers. Images of electrophoresis gel were analyzed with ImageLab (V. 6.0.1) to identify normal and mutant bands based on base pair length with BenchTop 100 bp ladder. Agarose gel were added with 3 µL FloroSafe DNA Stain 1st Base for each running with 4 µL PCR sample and 4 µL 100 bp ladder.

## 3. RESULTS AND DISCUSSIONS

### 3.1. Body weight

Inbreeding cross of F_1_ *Kambro* was initiated in the development of new chicken breed by increasing the allele homozigozity.

Based on measurement of Body Weight (BW), *Kambro* (1244.14 ± 453.82 grams) performed significantly (p<0.01) better than F_1_ *Pelung* (602.88 ± 79.93 grams) in 8 weeks period with *ad libitum* diet of standard feed (Mahardhika and Daryono, 2019). Inbred of F_1_ *Kambro* (n = 20) were F_2_ *Kambro* (n = 22) reached the average BW of 753.36 ± 155.31 gram in 8 weeks period measured with digital scaling KrisChef® EK9350H as shown in **Fig.1**. Based on ANOVA body weight of F_2_ *Kambro* was insignificant compared to F_1_ *Kambro* (p>0.001). Inbreeding depression had affected the body weight of inbred generation (F_2_ *Kambro*). Calculation of inbreeding coefficient (Fx) of F_2_ *Kambro* was 25% and inbreeding rate (IR) was 4.925 %. Inbreeding coefficient indicated an increase in allele homozigozity of inbred generation. Habibah (2018) stated that tolerance level of inbreeding coefficient is 37.5%. Inbreeding coefficient of F_2_ *Kambro* was in the tolerance level although it had shown a decline in phenotypic performance. Wakchaure and Ganguly (2015) stated that inbreeding depression has the greatest effect on reproductive traits, such as fertility, followed by productive traits, growth and milk production, with little or no effect on carcass traits.

**Fig. 1.**
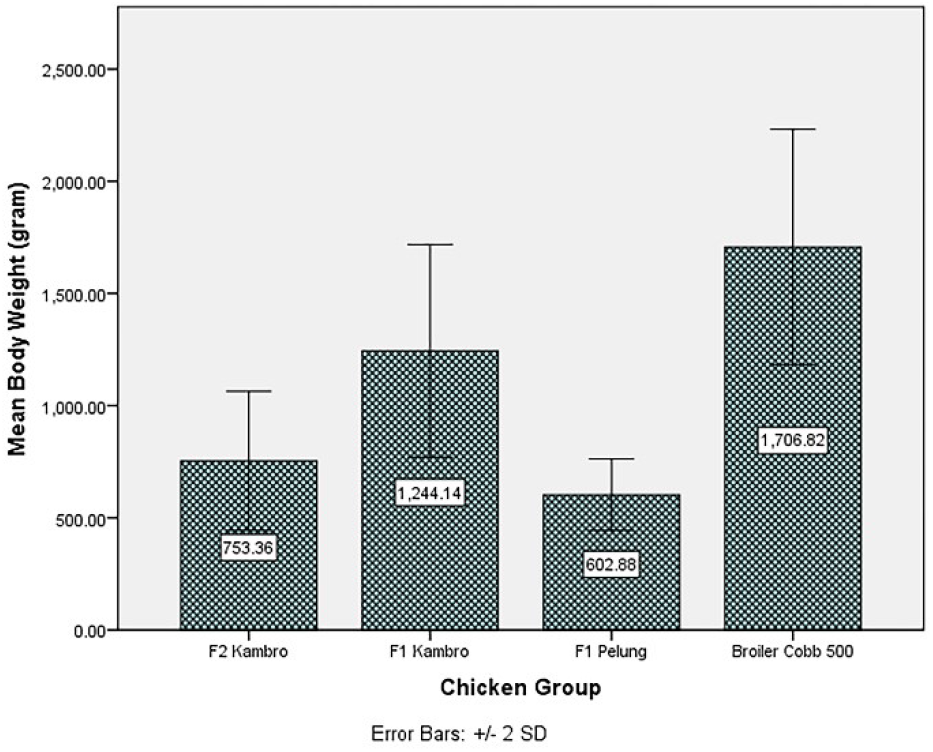
Body Weight of F_2_K, F_1_K, F_1_P dan BC5 at 8 weeks period. F_1_K: F_1_ *Kambro*; F_2_K: F_2_ *Kambro*; F_1_P: F_1_ *Pelung*; BC5: Broiler Cobb 500

### 3.2. Qualitative phenotypic groups

Based on visual observation, there are 6 groups of phenotypic character in the population F_2_ Kambro (n= 22) (**Fig.2.)**. F_2_ *Kambro* population was grouped based on boyd feather colour and shank colour which can be classified as group A (pure white), group B (black barred), group C (brown-white), group D (yellow white), group E (yellow-black), and group F (black-yellow). Group classification provided a validation of segregation in the allelic formation of parental generation.

**Fig. 2.**
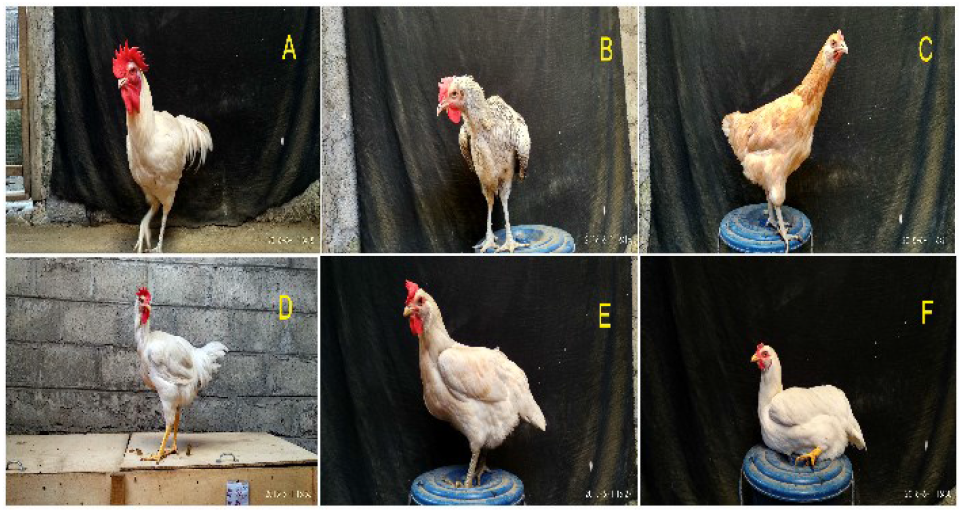
Phenotype groups of F_2_ *Kambro* population

### 3.3. Egg productivity

Egg productivity of female F_2_ *Kambro* (n = 4) reached 66 eggs during 6 months period with hen day production (HDP) of 16.5%. Egg productivity of female F_1_ *Kambro* (n = 4) reached 96 eggs during the same 6 months period with hen day production of 24%. Egg productivity of F_2_ *Kambro* was insignificant (p> 0.01) compared with F_1_ *Kambro*. Based on this result inbreeding depression had a significant impact on phenotypic performance of inbred generation, F_2_ *Kambro*.

### 3.4. T-ARMS PCR genotyping of *LEPR* (C127A)

Selective breeding assisted by both Mendelian and molecular genetics. Employment of genetics research accommodates the assessment process of parental generation in selective breeding and enables breeder to correctly selecting each hybrid for certain traits. In the effort to produce a native chicken breed with rapid growth and appealing phenotypic characteristics relative to Broiler Cobb 500 several genes were investigated, *LEPR* is amongst them. Leptin receptors (*LEPR*) have been located on neurons producing NeuroPeptide Y (NPY) and when activated by leptin binding, it is hypothesized to function in part by down-regulating the production of hypohypothalamic NPY (orexigenic effector) to inhibit ingestive behavior (Schwartz et al., 1997; Niv-Spector et al., 2005; Abbasi et al., 2011). Chicken (*Gallus gallus*) is an agriculturally important species and a model organism in developmental biology. Thus, the identification of chicken leptin and its possible role in metabolic regulation is of high interest (Seroussi et al., 2016). According to Seroussi et al. (2016) both leptin (*LEP*) and leptin receptor (*LEPR*) in mammalian species play a critical and nonredundant role in the control of food intake and energy expenditure, affecting body weight, fat accumulation, thermogenesis, insulin sensitivity, and lipid metabolism, and in addition, many other physiological processes such as puberty, reproductive cycle, immune response, bone growth and remodeling, and neural development. Recently the first genuine avian *LEP* were identified in the genomes of falcons (*Falco peregrinus* and *Falco cherrug*), Tibetan ground tit (*Pseudopodoces humilis*), zebra finch (*Taeniopygia guttata*), rock dove (*Columba livia*), bald eagle (*Haliaeetus leucocephalus*), downy woodpecker (*Picoides pubescens*), and budgerigar (*Melopsittacus undulatus*) (Seroussi et al., 2016). The identification approach in several studies regarding *LEP* gene association with number of chicken traits was carried out by identifying the presence of single nucleotide polymorphisms (SNPs) or concentrations of leptin receptors and *LEPR* mRNA expressions using the PCR-SSCP method (Wang et al., 2006). The association of *LEPR* SNPs and feed conversion ratio (FCR) efficiency in broiler chicken was determined by four *LEPR* genes SNPs associated significantly with feed intake which correlated positively with the weight growth rate and broiler chicken FCR (El Moujahid et al., 2014). Association of *LEPR* and ovarian dysfunction in broiler hens with variations in feed treatment in *ad libitum* found a positive correlation between ovarian follicle formation and the percentage level of leptin and leptin receptors in ovarian follicular tissue through regulation of the mechanism of steroidogenesis (Cassy et al., 2004). Genomic selection using gene marker can be implemented in order to minimize the effect of inbreeding during the selection period. Genomic selection is a promising alternative to conventional breeding for genetic improvement of layer chickens (Wolc et al., 2015). Tetra Primer Amplification Refractory Mutation System PCR (Tetra ARMS PCR or T-ARMS PCR) is a genotyping method based on the principle that PCR amplification is inefficient or completely refractory if there is a mismatch between the 3’ terminal nucleotide of a primer and its template sequence (Alyethodi et al., 2016; Peng et al., 2017). Tetra ARMS PCR incorporates 4 sets of primer, two outer primers (OF, OR) ensure the gene specificity and PCR efficiency, the inner outer combination (OF/IR, IF/OR) ensures the allele specificity which can be visualized by simple gel electrophoresis procedure (Alyethodi et al., 2016). Tetra ARMS PCR introduced as a simple, effective, and economical SNP genotyping method, but on contrary it requires difficult procedure for optimization and on some occasions fails to distinguish the target allele in SNP genotyping (Medrano and de Oliveira, 2014; Tanha et al., 2015). Designed Tetra ARMS PCR primers for single nucleotide polymorphism in exon 9 site of *LEPR* gene of chicken showed some minor disadvantages (Table 3.). Performance of Tetra ARMS PCR based on chicken DNA template requires further optimization to enhance its binding into mutated site. During the optimization experiment the results of experiment 9 was sufficient to be used in further detection of exon 9 SNP polymorphism of chicken *LEPR* gene. Experiment 9 protocol provided a high resolution DNA band with IP: OP ratio 10:1 pmol/µM, chicken DNA template concentration of 100 ng/µL with annealing temperature of 55.7°C /30s.

Based on genotyping result transversion mutation of *LEPR* (C127A) Exon 9 SNP can be detected using T-ARMS PCR method. Transversion mutation of C allele into A Allele *LEPR* gene were detected in **Fig.3.A** on line 3, 8, 9, 11, 13, 14, 15 and **Fig.3.B**. Line 2, 3 dan 4. Individual tag on each line were tag 2, tag 7, tag 8, tag 10, tag 12, tag 13, tag 14, tag 82b, tag 90b and tag 92b. Tag 2 (ChipChip3), tag 8 (ChipChip5), tag 12 (ChipChip2) and tag 14 (ChipChip4) of F_1_ Kambro population. Tag 7 (Igor), tag 10 (Odin) and tag 13 (Satrio) were F_2_ Kambro population. Tag 82b, tag 90b dan tag 92b were Broiler Cobb 500 population. Chicken with wild type allele C were detected in **Fig.3.A** on line 2, 4, 5, 6, 7, 10, 12, 16, 17 and **Fig.3.B** on line 1, 5, 6, 7, 8 and 9. Individual tag on each line were tag 1, tag 3, tag 4, tag 5, tag 6, tag 9, tag 11, tag 15, tag Bj, tag Ly, tag 97b, tag 3p, tag 7p, tag 8p and tag 10p. Tag 1 (ChipChip) and tag Bj (Bjorn) were F_1_ Kambro population. Tag 3 (Ragnar), tag 4 (Rollo), tag 5 (CaoCao), tag 6 (Clyde), tag 9 (Joy), tag 15 (Hilda) were F_2_ Kambro population. Tag 11 (*Pelung Blirik Hitam*), tag 3p, tag 7p, tag 8p and tag 10p were F_1_ *Pelung* population. Tag Ly was Layer population. Tag 97b was Broiler Cobb 500 population. Based on these results it can be concluded that there were a molecular correlation between body weight and egg productivity. In Guo et al. (2017) found that transversion mutations (Tv) can be disruptive to transcription factor binding (TF binding) compared to transition mutations. This disruptive effect is the inability of transcription of several amino acids that are associative to physiological functions. Based on genotyping results, Broiler Cobb 500 grandparent stock (GS) individuals have A allele transversion in exon 9 *LEPR gene*. This transversion mutation was inherited to F_1_ female *Kambro* parent stock (PS) whose inheritance can be detected in the F_2_ *Kambro* generation. From 5 DNA samples of female PS F_1_ *Kambro* chickens, 80% had A allele transversion in exon 9 *LEPR gene*. In F_2_ *Kambro* chickens from 15 DNA samples it was detected that 20% had A allele transversion with male sex. In the female GS generation of Broiler Cobb 500 from 4 DNA samples, 75% were detected having an A allele transversion in exon 9 *LEPR gene*. In the F_1_ *Pelung* group (n = 5) DNA samples, Layer (n = 1) DNA sample and the generation of F_2_ *Kambro* chickens (n = 12) DNA samples there were no A allele transversions in exon 9 *LEPR gene*. Transversion mutations in *LEPR* are vital because leptin is associated with chicken body weight gain and egg productivity. Abbasi et al. (2011) revealed that the *LEPR* polymorphism in Mazandaran chickens with the Restriction Fragment Length Polymorphism (RFLP) method indicated the dominance of A allele frequency. Molecular selection method with T-ARMS PCR can be used in selecting parental F_3_ *Kambro* generation. The effect of transversion and inbreeding depression affect the achievement of body weight and egg productivity in F_2_ *Kambro* chickens.

**Fig. 3.**
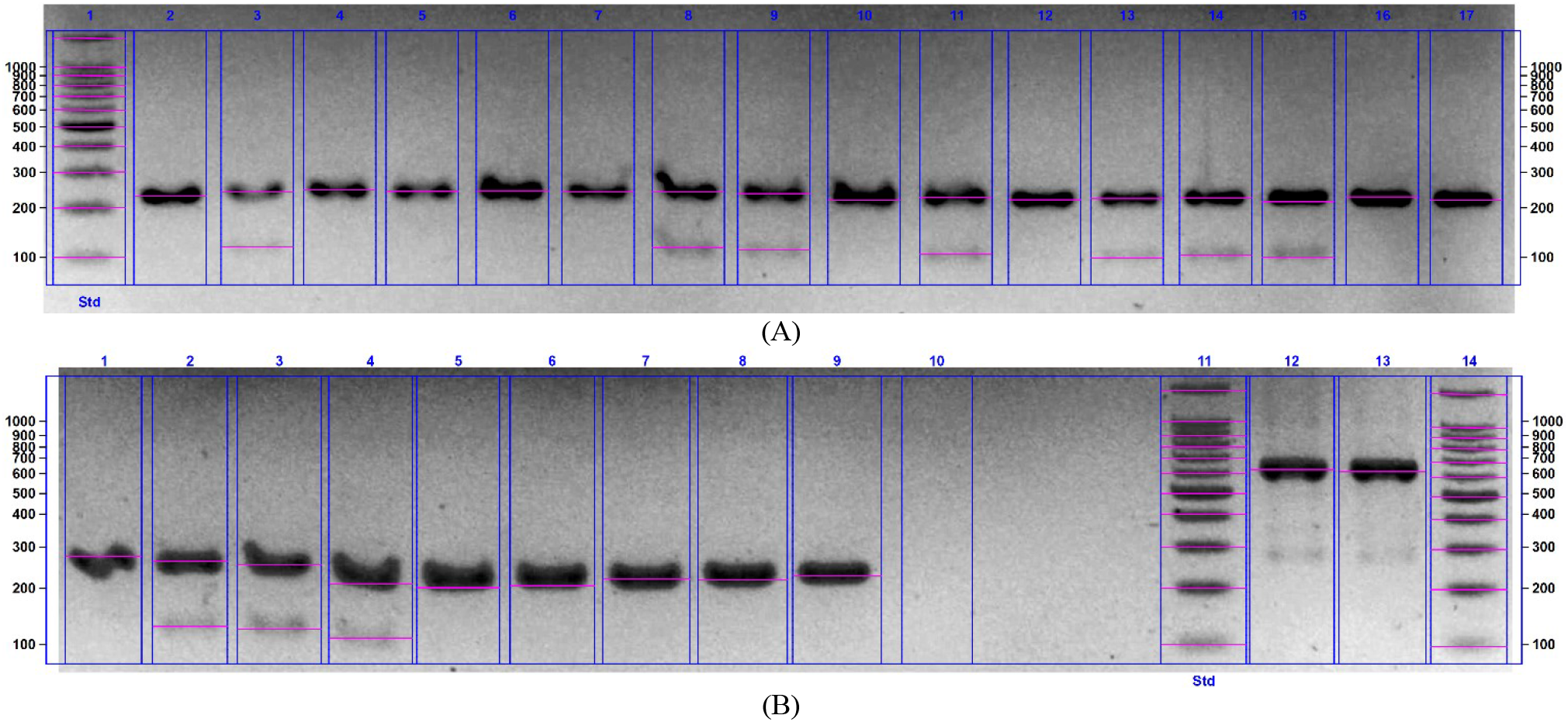
T-ARMS PCR result of Exon 9 SNP Polymorphism (C127A). (A) **Line 1**: marker 100 bp **Line 2**: OF/OR (C allele/wild type) (A) **Line 3, 8, 9, 11, 13, 14, 15**: A allele (A) **Line 2, 4, 5, 6, 7, 10, 12, 16, 17**: OF/OR (C allele/wild type) (B) **Line 2, 3, 4**: A allele (B) **Line 1, 5, 6, 7, 8, 9**: OF/OR (C allele/normal)

## 4. CONCLUSIONS

Qualitative phenotype parameters showed six groups of segregated phenotypes compared to F_1_ *Kambro* chicken. Growth of F_2_ *Kambro* chicken weight reached 753.36 ± 155.31 grams in 8 weeks was not significant for F_1_ *Kambro* chicken due to inbreeding depression (Fx = 25%, IR = 4.925%) and transversion of A *LEPR* allele mutations. Specific protocol detection of exon 9 *LEPR* gene SNPs using the T-ARMS PCR method can detect C127A *LEPR* mutations with IP: OP ratio 10:1 pmol / µM, chicken DNA template concentration of 100 ng / µL with annealing temperature of 55.7 °C/30s. The transversion mutation of C127A of *LEPR* exon 9 SNP were detected in DNA samples of F_1_ *Kambro* hens (80%), F_2_ *Kambro* roosters (20%), Broiler Cobb 500 hens (75%). The mutations were not detected in Layer, *Pelung Blirik Hitam* chicken and F_1_ *Pelung* populations.

