## Supplementary material for "Exon 9 *LEPR* Gene SNP Polymorphism of Hybrid Chickens F_2_ *Kambro* Crossbreeds of ♀ F_1_ *Kambro* with ♂ F_1_ *Kambro*": Kambro_manuscript.pdf

Indonesia Indigenous Chicken Genetic Profile

### Genetic Profile of *Kampong* Broiler-Type Chicken, Progenies of Indonesia Indigenous Chicken (*Gallus gallus*, Linn. 1758)

**Abstract**

*Kambro* (*Kampong* Broiler-Type) are the progenies Indonesia indigenous chicken (*Gallus gallus*, Linn 1758) with the sole purpose to be a fast-growing meat-type breed through selective breeding of *Pelung* beneficial traits. *Pelung* known in Indonesia as *ayam Kampong* is more adaptive, resistance to disease and preferable by the locals, although it has slow-growing performance. Leptin receptor gene (LEPR) single nucleotide polymorphism is known to have associated with several commercial and fitness traits in chicken such as body weight performance and productivity in general. In this work, we investigated the LEPR polymorphism association with body weight (BWT) and inbreeding depression in hybrid chicken *Kambro*. Whole blood samples of *Pelung Blirik Hitam*, $F_1$ *Pelung*, Broiler Cobb 500, $F_1$ *Kambro* and $F_2$ *Kambro* were collected. Two primer sets specifically designed to detect chicken exon 9 LEPR gene SNP (NCBI Ref. Seq. **AY048693.1**) using web-based software. We hypothesized that T-ARMS-PCR could detect LEPR polymorphism in the hybrid chicken *Kambro* with several optimization adjustments. The presence of a single nucleotide variant of **AY048693.1**: g. 127C>A detected in LEPR locus, with allele A frequency of 0.416. *Kambro* chicken and Broiler Cobb 500 both found to have a single nucleotide variant of g. 127C>A. Intercrossing of $F_1$ *Kambro* resulted in inbreeding depression and LEPR polymorphism in the hybrid chicken *Kambro* chicken was found to affect body weight. Single nucleotide polymorphism and its association reported in this preliminary study could be used as a selection marker and a tool in breeding programs for broader types of indigenous meat-type chicken.

**Keywords**

selective breeding; Indonesian indigenous chicken; *Pelung*; body weight; meat-type chicken; intercross; chicken germplasm

#### Introduction

A trend of exploiting indigenous chicken breed into local poultry sector has been increasing especially in developing countries for example a study on Korean native chicken (Manjula et al, 2018), Nigerian native chicken (Nwenya et al, 2017) and Mazandaran indigenous chicken (Niknafs et al, 2013). Nwenya et al (2017) stated that great genetic resources embedded in the indigenous poultry await full exploitation that will provide basis for genetic improvement and diversification to produce breeds that are adapted to local conditions for the benefits of farmers especially in developing countries. Manjula et al (2018) stated that there are numerous indigenous chicken breeds and their economic traits and genetic potential remain largely unknown. Recently, much attention has been focused on indigenous chickens as meat or layer strains because of increasing consumer demand and environmentally viable characteristics of local ecotypes (Manjula et al, 2018). In classification, Indonesia's indigenous chicken are identified by meat-type, laying-type and ornamental-type. Henuk & Bakti (2018) classified Indonesia's indigenous chicken into 34 distinct breeds, *Ayunai, Balenggek, Banten, Bangkok, Burgo, Bekisar, Cangehgar, Cemani, Ciparage, Gaok, Jepun, Kampung, Kasintu, Kedu (hitam* and *putih), Pelung, Lamba, Maleo, Melayu, Merawang, Nagrak, Nunukan, Nusa Penida, Olagan, Rintit atau Walik, Sedayu, Sentul, Siem, Sumatera, Tolaki, Tukung, Wareng, Sabu,* and *Semau*. Indonesia's indigenous chicken breed known as *Ayam Kampung* has high nutritional and has high demand in the local market. *Kampung* chicken is mostly bred by villgers in Indonesia because it is easily maintained, has high nutritional meats, strong posture, and *Ayam Kampung*'s egg are also in high demand due to its nutritional content.

The fulfillment of 67% poultry product consumption is dominated by broilers and 23% of Indonesia indigenous chickens, with a total contribution of 60.73% of poultry farms to national animal food production (Suprijatna, 2010). The indigenous Indonesian broiler and layer livestock industry are experiencing rapid development with increasing interest and community involvement in the native chicken or free-range chicken poultry industry (Iskandar, 2017). Increased productivity and quality competitiveness of native broilers can be achieved by selective breeding of Indonesian native chicken breeds. Selective breeding aims to produce chicken breeds with certain phenotypic qualities according to human needs (Das et al, 2008; Cheng, 2010; Oldenbroek & van der waaij, 2014; Mariandayani et al, 2017; Sudrajat & Isyanto, 2018).

In this study we exploited the genetic resource of *Pelung* chicken from Cianjur, West Java, Indonesia. *Pelung* chicken has a higher body weight growth, unique meat flavor and superior posture compare with other indigenous breeds (Mahardhika and Daryono, 2019). Selective breeding program conducted between *Pelung Blirik Hitam* and Broiler Cobb 500 produces hybrid chicken *Kampong* Broiler-Type (*Kambro*). Body weight (BWT) of *Kambro* (1244.14 ± 453.82 grams) significantly (p <.001) found to be surpassing $F_1$ *Pelung* (602. 88 ± 79.93 grams) at eight weeks semi-intensive rearing period with *ad libitum* standard feed diet (Mahardhika & Daryono, 2019). Further selective breeding carried out in intercross between $F_1$ *Kambro* resulted in $F_2$ *Kambro* chicken. Accompanying selection for rapid growth, meat-type chicken exhibit an increase in physiological disorders such as obesity. Production performance and fitness traits were negatively correlated in chicken (Martin et al, 1990; Pinard et al, 1998). Multi-traits selection to simultaneously improve fitness and increase production is, therefore, difficult to achieve by conventional direct screening and selection.

Molecular Marker Assisted Selection (MAS) may be required and the integration of traditional genetic selection and modern molecular methods may be preferred for breeding chickens in the future (Li et al, 2003). Advances in molecular biotechnology enable fast and reliable methods for the accurate diagnosis of mutations responsible for different genetic defects (Alyethodi et al, 2016). These assist breeders to identify carriers at an early stage (Alyethodi et al, 2016).

The Amplification Refractory Mutation System (ARMS)-PCR (Newton et al, 1989) and tetra-primer PCR (Ye et al, 1992) were able to detect known sequence polymorphisms. The combination of aforesaid two techniques generated tetra-primer ARMS-PCR or T-ARMS-PCR technique (Ye et al, 2001). ARMS-PCR has several advantages, namely the need for samples is small, rapid and effective, efficient and simultaneous, the level of sensitivity and accuracy are high and consistent (Peng et al, 2017). LEPR gene polymorphism is correlated with the weight, BW gain and feed intake of meat-type chickens (El Moujahid et al, 2014; Kaczor et al, 2016).

The implementation of this method using Indonesian native chicken DNA samples to detect single nucleotide polymorphism (SNP) at exon 9 chicken (*Gallus gallus*) LEPR gene has never been studied. In this work, we investigated exon 9 LEPR gene SNP (NCBI Ref. Seq. **AY048693.1**: g. 127C>A) with T-ARMS-PCR to develop specific MAS in hybrid chicken *Kambro* selective breeding program. Association of exon 9 LEPR gene SNP with body weight (BWT) and inbreeding depression in hybrid chicken *Kambro* were investigated.

#### Material and Methods

##### *Ethical clearance*

This study was performed in accordance with the Animal Welfare Act of Indonesia and all procedures involving handling of animals were approved by the local office of occupational and technical safety.

*Sample collection*

Hybrid chickens ($F_1$ *Pelung*, *Kambro* and $F_2$ *Kambro*) and parental generation (*Pelung Blirik Hitam* and Broiler Cobb 500) reared at the semi-intensive system. $F_2$ *Kambro* produced through intercrossing of $F_1$ *Kambro*. $F_1$ *Pelung* was produced through crossbreeding of *Pelung Blirik Hitam*. Chicken of each population went through screening phase and then selected for molecular analysis (Table 1). Body weight (BWT) measurement and traits observation were conducted. Genomic DNA was isolated from whole blood sample by chelex based method (Ernanto et al, 2018). DNA concentration and purity were quantified with Spark® Reader spectrophotometer (TECAN). DNA later kept dissolved in TE buffer (pH 8.0) and stored in freezer at -20℃ for further use.

*T-ARMS-PCR primers design*

Single nucleotide polymorphism of exon 9 chicken LEPR gene were acquired from NCBI Ref.Seq. **AY048693.1**. BLAST alignment between **AY048693.1** and **AF222783.1** was conducted to detect SNP. Exon 9 chicken LEPR gene was found in **AY048693.1**: g. 127C>A. T-ARMS-PCR primers were designed with web-based software http://primer1.soton.ac.uk (Peng et al, 2017). Four primers were designed (FIA: forward inner primer A allele; RIC: reverse inner primer C allele; FOP: forward outer primer; ROP: reverse outer primer) enabling the specific amplification of normal and mutant alleles (Table 2). Allele-specific amplicons with different product lengths were separated by standard agarose gel electrophoresis. Designed primers produced by Integrated DNA Technologies (IDT) with third-party PT. Genetika Science Indonesia.

*In vitro amplification, visualization and analysis*

The composition of each Tetra ARMS PCR/T-ARMS PCR reaction mix consisted of 50-100 ng of genomic DNA with concentration range from 1-2% (260/280 nm), 200 µM of each dNTP at 1X and 0.5 U per 25 µL reaction, KAPA Taq DNA polymerase with buffer containing MgCl2 (1.5 mM at 1X) and stabilizers. The final reaction volume of 25 µL was made with nuclease-free water. Each PCR reaction were amplified with BIORAD T100™ PCR Thermal Cycler. The PCR products were detected by electrophoresis with Submarine Electrophoresis System (Mupid-EXU) device. Finally, observations were performed under ultraviolet light (λ = 260 nm) AnalytikJena™ gel imaging system and documented with GelDoc™ Documentation System.

A series of experiments were performed to validate the performance of Tetra ARMS PCR to detect chicken LEPR gene by changing primers concentration ratio, electrophoresis phase and PCR steps as can be seen in (Table 3). Images of electrophoresis gel were analyzed with ImageLab (V. 6.0.1) to identify each band based on base-pair length with BenchTop 100 bp ladder.

**Statistical analysis**

Body weight of each population was analyzed with one-way-ANOVA using IBM SPSS Statistics version 21. Inbreeding coefficient (Fx) and inbreeding rate (*F*) of inbred generation *Kambro* were calculated with the following formula:

Fx = inbreeding coefficient value,
Σ = sum, in case of multiple inbreeding on same or more common ascendants,
n = the number of connecting links between the two parents of X through common ancestors,
n' = number of generations between animal X and common ascendant A, maternal line,
FA = inbreeding coefficient of the common ancestor A.

(Telalbasic et al, 2007)

*F* = Inbreeding rate
Nm = number of male
Nf = number of female

(Perdamaian et al, 2017)

The frequencies of LEPR alleles and genotypes were determined and it was verified with HW *P*-value whether their distributions conformed to those expected according to the Hardy-Weinberg law.

**Results**

*Kambro body weight (BWT), phenotypic traits and inbreeding coefficient*

Intercrossing (inbreeding cross) of $F_1$ *Kambro* parental generation (age ± 1 year) under a semi-intensive rearing system used a ratio of 1 male: 2 female. Intercrossing of $F_1$ *Kambro* was initiated in the development of the new chicken breed to increase the allele homozygosity. Immunity depreciation in hybrid chicken generation can be caused by the presence of inbreeding or crossing of related individual (Oldenbroek & van der Waaij, 2014). Inbreeding cross of $F_1$ *Kambro* chickens aims to strengthen the inheritance of several traits, primarily fast-growth performance. Free-range chickens also are known locally as *Ayam Kampong* in Indonesia has been classified into 31 Indonesian native chicken breeds with *Pelung Blirik Hitam* as one of them. The *Mx*| *Hpy 81* gene demonstrated a high potential for use as a genetic marker for resistance to Avian Influenza and Newcastle Disease infection in Indonesian native chickens (Pagala et al, 2017). Fulton et al (2014) stated that polymorphisms of the *Mx* gene have been reported in multiple breeds of chickens, including Australorp, Fayoumi, Japanese native chickens, Indonesian native chickens, White Leghorns, Broilers and inbred laboratory lines. Pagala et al (2017) found that *Ayam Kampong* and *Tolki* chicken are resistant to virus attacks such as Avian Influenza (AI) and Newcastle Disease (ND) due to the flow of the A allele, which causes serine amino acid (AGT) changes to asparagine (AAT). *Kambro* line is expected to be a prominent meat-type breed with higher resistance to disease, fast-growth performance and more adapt to tropical climate.

In Table 4**.** BWT in chicken populations $F_1$K, $F_2$K, $F_1$P and $BC_5$ differed significantly [F (3, 53) = 68,896, p <.001, η2 = .796]. Significantly BWT $BC_5$ (M = 1706.82, SD = 262.54, p <.001) were superior to BWT $F_1$K, $F_2$K and $F_1$P, but $F_2$K (M = 753.36, SD = 155.31) did not differ significantly from $F_1$P (M = 602.88, SD = 79.93). The $F_1$K group (M = 1244.14, SD = 453.82, p <.001) was significantly superior to the $F_2$K group. Based on body weight (BWT) measurement, *Kambro* (1244.14 ± 453.82 grams) performed significantly (p<0.01) better than $F_1$ *Pelung* (602.88 ± 79.93 grams) at eight weeks semi-intensive rearing period with *ad libitum* standard feed diet (Mahardhika & Daryono, 2019). Inbred $F_2$ *Kambro* reached the average body weight of 753.36 ± 155.31 gram at eight weeks semi-intensive rearing period with *ad libitum* standard feed diet measured with digital scaling KrisChef® EK9350H as shown in Figure 1.

Based on ANOVA body weight of $F_2$ *Kambro* was insignificant compared to $F_1$ *Kambro* (p>0.001). Inbreeding depression had affected the body weight of inbred generation ($F_2$ *Kambro*). The calculation of the inbreeding coefficient (Fx) of $F_2$ *Kambro* was 25% and the inbreeding rate (*F*) was 4.925 %. Phenotypic traits parameter including neck feather color, dorsal/ back feather color, chest feather color, body feather color, femoral feather color, shank color, comb color, comb shape, and beak color were identified as visual data with the black background photo. Inbred *Kambro* has six phenotypic trait groups, classified as Pure White, Black-barred, White-Chocolate, Yellow-White, Yellow-Black, and Yellow (Figure 2**)**. Ratio of male and female chickens in $F_2$ *Kambro* population was 4: 7. In the $F_2$ *Kambro* chicken population the percentage per phenotype group is A (54.55%), B (9.09%), C (9.09%), D (9.09%), E (9.09%), F (9.09%).

*LEPR polymorphism and body weight association*

T-ARMS-PCR and Based on T-ARMS PCR method single nucleotide variant of g. 127C>A can be detected through series of optimization experiments. Exon 9 chicken LEPR gene (NCBI Ref.Seq. **AY048693.1**: g. 127C>A) with T-ARMS-PCR method could detect the single nucleotide variant on each population through series of optimization (Figure 3).

In total there were 11 experiments conducted to optimize the result of digested amplicons. One experiment showed a promising result with clear and distinctive bands shown. Specific detection protocol of exon 9 chicken LEPR gene (NCBI Ref.Seq. **AY048693.1**: g. 127C>A) in *Kambro* chicken line with T-ARMS PCR method requires inner primer (IP): outer primer (OP) ratio of 10:1 pmol / µM, chicken DNA template concentration of 100 ng / µL with annealing temperature of 55.7 ˚C / 30s (touchdown). Allele frequencies were calculated based on electrophoresis result as shown in Table 5.

The use of the T-ARMS-PCR method showed the single nucleotide variant of exon 9 chicken LEPR gene g. 127C>A in LEPR locus, allele A emerged with a high frequency of 0.416 (Table 5). The linear model used in the statistical analysis of the data included the line, sex, and LEPR, and the interactions between experimental factors. The chicken lines used in the study were fast-growing lines and no effect of the line on the production traits was found. No line X LEPR interaction was also found. Sex was found to have a statistically significant effect, but due to the lack of sex X LEPR interaction, the effect of this factor was ignored in the analysis of the results. In this preliminary study, we found the absent of heterozygosity and significant deviation from Hardy-Weinberg equilibrium within the population of Broiler Cobb 500, *Pelung* and *Kambro*. The use of the T-ARMS-PCR method showed the single nucleotide variant of exon 9 chicken LEPR gene g. 127C>A in LEPR locus. Allele A emerged with high frequency on the population of Broiler Cobb 500 (BC5), $F_1$ *Kambro* ($F_1$K) and $F_2$ *Kambro*�($F_2$K), each represented by allele A frequency of 0.75, 0.66 and 0.33, respectively (Table 6**).**

The population of *Pelung* ($F_1$P) showed monomorphic results with allele C frequency of 1. A similar result published about the study about exon 9-11 polymorphism in Mazandaran fowl (Abbasi et al, 2011). Abbasi et al (2011) concluded that Mazandaran fowl digestion product is monomorphic which showed allele B frequency of 1 from PCR-RFLP of LEPR gene exon 9-11.

Polymorphism of exon 9 chicken LEPR gene g. 127C>A showed a significant association with body weight (BWT). The highest BWT was found in the chicken population with higher allele A frequency respectively. Broiler Cobb 500 (BC5), $F_1$ *Kambro* ($F_1$K) and $F_2$ *Kambro* ($F_2$K) with allele A frequency of 0.75, 0.66 and 0.33 have represented significantly difference ($P<0.05$) body weight value of 1706.82 (± 262.54) grams, 1244.14 (± 453.82) grams and 753.36 (± 155.31) grams, respectively. The population of *Pelung* ($F_1$P) with allele A frequency of 0 showed a significantly lower ($P<0.05$) body weight value of 602.88 (± 79.93) grams, in comparison with $BC_5$ and $F_1$K but does not differ with $F_2$K.

#### Discussion

Tetra Amplification Refractory Mutation System Polymerase Chain Reaction (T-ARMS-PCR) has been widely used to detect single nucleotide polymorphism (SNP). Marker Assisted Selection (MAS) has been used in selective breeding to provide a faster, more accurate and reliable selection method. By combining T-ARMS-PCR and LEPR polymorphism we have developed MAS in the selection of meat-type chicken breed, *Kambro* in particular. Allele C frequency of 1 in *Pelung* showed a slow-growing broiler chicken, on the opposite Broiler Cobb 500 is known as fast-growing broiler chicken with allele A frequency of 0.5625 and allele C frequency of 0.0625. *Pelung Blirik Hitam* has several distinguished characters such as posture and higher body weight compare with other native Indonesian chicken breeds (Daryono et al. 2010, Mahardhika and Daryono 2019). Body weight of 1-year-old male *Pelung* chicken can reach 3.37 kg and female *Pelung* can reach 2.52 kg, under a semi-intensive rearing system (Daryono et al, 2010, Mahardhika & Daryono, 2019). Although *Pelung* has the potential to be a meat-type candidate, *Pelung* has a slow-growth and therefore must be improved. This improvement later carried out through a selective breeding program resulted in *Kambro,* an abbreviation for *Kampong*-Broiler. *Kambro* chicken line inherited the allele A and showed a distinct performance in growth compare to *Pelung.* Allele A frequency in $F_1$ *Kambro* and $F_2$ *Kambro* are 0.66 and 0.33, higher than *Pelung.* On the intra-population of *Kambro,* the F1 *Kambro* generation showed higher allele A frequency that the generation of F2 *Kambro*. Polymorphism of exon 9 chicken LEPR gene g. 127C>A with body weight was found to affect the body weight (BWT). The present effect of LEPR polymorphism on BWT is consistent with the result of studies with broiler chicken (Ross 308 and Hubbard Flex) (Kaczor et al, 2016).

Geneticists have made rapid genetic improvements through the use of intense selection for specific biological traits, leading to continuous improvements in body weight (BWT), growth rate, and meat yield in meat-type birds (Deeb & Lamont, 2002). In this study we purposely conducted an inbreeding to increase homozygosity of certain alleles related to the higher growth performance. Inbreeding is the main challenge in *Kambro* chicken line selective breeding program. The absent of heterozygosity detected in *Kambro* chicken based on LEPR polymorphism showed a promising opportunity to be further improved. Inbreeding depression had affected the body weight of inbred generation ($F_2$ *Kambro*). The inbreeding coefficient (Fx) of $F_2$ *Kambro* was 25% and the inbreeding rate (*F*) was 4.925 %. The inbreeding coefficient indicated an increase in allele homozygosity of inbred generation. (Habibah et al, 2018 *unpublished data*) stated that the tolerance level of the inbreeding coefficient is 37.5%. The inbreeding coefficient of $F_2$ *Kambro* was in the tolerance level although it had shown a decline in phenotypic performance. Perdamaian et al (2017) found that the inbreeding coefficient of 25% is classified as high amongst domesticated chicken. (Wakchaure & Ganguly, 2015) stated that inbreeding depression has the greatest effect on reproductive traits, such as fertility, followed by productive traits, growth and milk production, with little or no effect on carcass traits. Based on Hardy-Weinberg equilibrium *Kambro* chicken significantldeviated, thus showed a significant transformation of allelic distribution.

In comparison, a study conducted by (Abebe et al, 2015) about the genetic diversity of five local Swedish chicken breeds discussed the same result. All of the five breeds were significantly deviated from Hardy-Weinberg expectations and across all breeds, more than half of the loci showed significant deviation from Hardy-Weinberg (Abebe et al, 2015). Keeping small isolated flocks over many generations may result in loss of heterozygosity due to the high chances of random genetic drift and inbreeding (Abebe et al, 2015). Therefore, outbreeding can be a solution to this inbreeding problem, by introducing outer-generation parental from the same breed. Higher mortality rate also observed in $F_2$*Kambro*, declining performance in disease resistance and sudden death cause serious problem in the selective breeding. The increase in physiological disorders such as obesity, ascites, sudden death syndrome, and leg problems, as well as a reduction in overall immunocompetency have become important issues (Deeb & Lamont, 2002).

Leptin receptor gene (LEPR) have been located on neurons producing NeuroPeptide Y (NPY) and when activated by leptin binding, it is hypothesized to function in part by down-regulating the production of hypothalamic NPY (orexigenic effector) to inhibit ingestive behavior (Schwartz et al, 1997; Niv-Spector et al, 2005; Abbasi et al, 2011). Chicken (*Gallus gallus*) is an agriculturally important species and a model organism in developmental biology. Thus, the identification of chicken leptin and its possible role in metabolic regulation is of high interest (Seroussi et al, 2016). Seroussi et al (2016) found that both leptin (LEP) and leptin receptor (LEPR) in mammalian species play a critical and non-redundant role in the control of food intake and energy expenditure, affecting body weight, fat accumulation, thermogenesis, insulin sensitivity, and lipid metabolism, and besides, many other physiological processes such as puberty, reproductive cycle, immune response, bone growth and remodeling, and neural development. Recently the first genuine avian LEP were identified in the genomes of falcons (*Falco peregrinus* and *Falco cherrug*), Tibetan ground tit (*Pseudopodoces humilis*), zebra finch (*Taeniopygia guttata*), rock dove (*Columba livia*), bald eagle (*Haliaeetus leucocephalus*), downy woodpecker (Picoides pubescens), and budgerigar (*Melopsittacus undulatus*) (Seroussi et al, 2016). The identification approach in several studies regarding the LEP gene associated with several chicken traits was carried out by identifying the presence of single nucleotide polymorphisms (SNPs) or concentrations of leptin receptors and LEPR mRNA expressions using the PCR-SSCP method (Wang et al, 2006). The association of LEPR SNPs and feed conversion ratio (FCR) efficiency in broiler chicken was determined by four LEPR genes SNPs associated significantly with feed intake which correlated positively with the weight growth rate and broiler chicken FCR (El Moujahid et al, 2014). Genomic selection using a gene marker can be implemented to minimize the effect of inbreeding during the selection period. Genomic selection is a promising alternative to conventional breeding for genetic improvement of layer chickens (Wolc et al, 2015). Tetra Primer Amplification Refractory Mutation System Polymerase Chain Reaction (Tetra ARMS PCR or T-ARMS-PCR) is a genotyping method based on the principle that PCR amplification is inefficient or completely refractory if there is a mismatch between the 3' terminal nucleotide of a primer and its template sequence (Alyethodi et al, 2016; Peng et al, 2017). Tetra ARMS PCR incorporates 4 sets of primer, two outer primers (OF, OR) ensure the gene specificity and PCR efficiency, the inner-outer combination (OF/IR, IF/OR) ensures the allele specificity which can be visualized by simple gel electrophoresis procedure (Alyethodi et al, 2016). Tetra ARMS PCR introduced as a simple, effective, and economical SNP genotyping method, but on contrary, it requires a difficult procedure for optimization and on some occasions fails to distinguish the target allele in SNP genotyping (Medrano & de Oliveira, 2014; Tanha et al, 2015). Performance of Tetra ARMS PCR based on chicken DNA template requires further optimization to enhance its binding into the mutated site. During the optimization experiment, the results of experiment 9 were sufficient to be used in further detection of exon 9 SNP polymorphism of the chicken LEPR gene. Experiment 9 protocol provided a high-resolution DNA band with IP: OP ratio 10:1 pmol/µM, chicken DNA template concentration of 100 ng/µL with an annealing temperature of 55.7℃/30s.

Based on these results, a conclusion can be drawn from allele frequency and genotype frequency of Broiler Cobb 500, *Pelung,* $F_1$ *Kambro,* and $F_2$ *Kambro. Kambro* chicken breed offers a promising opportunity to be developed further in the selective breeding program. The inbreeding program conducted has reduced heterozygosity in *Kambro* chicken line. Further consideration to reduce inbreeding depression can be taken through outbreeding with different parental from the same breed of *Pelung.* The effort to develop a meat-type, fast-growth chicken from native Indonesian chicken as parental, in this case, *Pelung* chicken is possible. Marker Assisted Selection (MAS) of *Kambro* chicken with LEPR polymorphism could be used in future *Kambro* selective breeding program or broader meat-type chicken breeds.

**Conflict of Interest**

Authors have no conflict of interest to declare.

Wolc, A., Zhao, H.H., Arango, J., Settar, P., Fulton, J.E., O'Sullivan, N.P., Preisinger, R., Stricker, C., Habier, D., Fernando, R.L., Garrick, D.J., Lamont, S.J., & Dekker, J.C.M.

2015. Response and inbreeding from a genomic selection experiment in layer chickens. *Genomic Selection Evolution*. 47: 59.
https://doi.org/10.1186/s12711-015-0133-5

Ye, S., Humphries, S., & Green, F. 1992. Allele specific amplification by tetra primer PCR. *Nucleic Acid Research*. 20(5): 1152.
https://doi.org/10.1093/nar/20.5.1152

Ye, S., Dhillon, S., Ke, X., Collins, A.R., & Day, I.N.M. 2001. An efficient procedure for genotyping single nucleotide polymorphisms. *Nucleic Acid Res.* 29(17).
https://doi.org/10.1093/nar/29.17.e88

**Table 1**

Selected chicken of each population group sample tags for preliminary optimization of T-ARMS PCR.

| Population Group | | | | | | |
|---|---|---|---|---|---|---|
| $F_1$P (n:5) | | BC5 (n:4) | $F_1$K (n:6) | | $F_2$K (n:9) | |
| ♀ | ♂ | ♀ | ♀ | ♂ | ♀ | ♂ |
| 3p | PBH | 82b | ChipChip | Bjorn | CaoCao | Ragnar |
| 7p | | 90b | ChipChip2 | | Joy | Rollo |
| 8p | | 92b | ChipChip3 | | Hilda | Clyde |
| 10p | | 97b | ChipChip4 | | | Igor |
| | | | ChipChip5 | | | Odin |
| | | | | | | Satrio ( |

$F_1$P: *Pelung*; PBH: *Pelung Blirik Hitam;* BC5: Broiler Cobb 500; $F_1$K: first generation *Kambro*; $F_2$K: second generation *Kambro*

**Table 2**
T-ARMS-PCR primers design based on exon 9 *LEPR* gene SNP (NCBI Ref. Seq. **AY048693.1**) with web-based software http://primer1.soton.ac.uk

| Primers | Sequence (5'-3') | Melting temperature (℃) |
|---|---|---|
| FIA (A allele) | 251 TTACTCTTTTCAACTTGAAAGCAGCA 276 | 55,5 |
| RIC (C allele) | 304 CGTTATAGAAGAAACTTCCTCTAGGTCTG 276 | 55,9 |
| FOP (5' - 3') | 136 ATCTATAAAAACAAAACCCAGAGCGTA 162 | 54,8 |
| ROP (5' - 3') | 363 TACATATAATTCAGCGTATCTATGATGGC 335 | 54,6 |

| |
|---|
| Product size for A allele: 114 bp |
| Product size for C allele: 169 bp |
| Product size of two outer primers: 228 bp |

FIA: forward inner primer A allele; RIC: reverse inner primer C allele; FOP: forward outer primer; ROP: reverse outer primer

#### Table 3

Alteration and modification of primers concentration, electrophoresis phase and PCR configurations.

| Exp. | DNA | PC (pmol/µM) | PR (µL) | | | | PCR Steps | | | | | | Ag | Electrophoresis | |
|---|---|---|---|---|---|---|---|---|---|---|---|---|---|---|---|
| | (ng/ µL) | IP/OP | FIA | FIC | FOP | ROP | ID (℃/mins) | Cyl (X) | DS (℃/s) | A (℃/s) | E (℃/s) | FE (℃/mins) | (%) | V (volt) | D (mins) |
| 1 | 50 | 10/10 | .5 | .5 | .5 | .5 | 94/5 | 40 | 95/30 | Gradient/30 | 68/60 | 68/5 | 2 | 50 | 55 |
| 2 | 50 | 10/10 | .5 | .5 | .5 | .5 | 94/5 | 40 | 95/30 | 56.2/30 | 68/60 | 68/5 | 2 | 50 | 55 |
| 3a | 50 | 10/10 | .75 | .75 | .25 | .25 | 95/5 | 36 | 95/60 | 56.2/60 | 72/60 | 72/10 | 2 | 50 | 55 |
| 3b | 50 | 10/10 | .75 | .75 | .25 | .25 | 94/5 | 40 | 95/30 | 56.2/30 | 68/60 | 68/5 | 2 | 50 | 55 |
| 4 | 50 | 10/10 | .9 | .9 | .1 | .1 | 95/2 | 30 | 95/60 | 56.2/60 | 72/60 | 72/10 | 2 | 50 | 55 |
| 5 | 50 | 10/10 | 1.5 | 1.5 | 1 | 1 | 95/2 | 30 | 95/60 | 55.7/60 | 72/60 | 72/10 | 2 | 50 | 55 |
| 6a | 50 | 10/10 | 1.25 | 1.25 | .25 | .25 | 95/2 | 30 | 95/60 | 55.7/60 | 72/60 | 72/10 | 2 | 50 | 55 |
| 6b | 50 | 10/10 | .75 | .75 | .25 | .25 | 94/5 | 40 | 95/30 | 56.2/30 | 68/60 | 68/5 | 2 | 50 | 55 |
| 7 | 50 | 10/10 | .75 | .75 | .25 | .25 | 94/5 | 40 | 95/30 | 56.2/30 | 68/60 | 68/5 | 2 | 50 | 55 |
| 8 | 50 | 10/10 | .75 | .75 | .25 | .25 | 95/5 | 36 | 95/60 | 56.2/60 | 72/60 | 72/10 | 2 | 50 | 55 |
| 9 | 100 | 10/1 | .5 | .5 | .5 | .5 | 94/ 5 | 30 | 94/30 | 55.7/30 | 72/40 | 72/10 | 2.5 | 50 | 40 |

FIA: Forward Inner A Allele; FIC: Forward Inner C Allele; FOP: Forward Outer Primer; ROP: Reverse Outer Primer; PC: Primers Concentration; PR: Primers Ratio; IP/OP: Inner Primers/Outer Primers; ID: Initial Denaturation; Cyl: Cycle; DS: Denaturation Steps; A: Annealing; E: Extension; FE: Final Extension; Ag: Agarose; V: Voltage; D: Duration; Gradient: 53˚C, 53.3˚C, 53.8˚C, 54.6˚C, 55.5˚C, 56.2˚C, 56.7˚C and 57˚C; Exp: Experiment (1, 2, 3a, 3b, 4, 5, 6a, 6b, 7, 8 and 9).

#### Table 4

Bodyweight (BW) analysis on chicken population $F_1K$, $F_2K$, $F_1P$ and BC5.

| Traits | Chicken Group | | | | | |
|---|---|---|---|---|---|---|
| | $F_2K$ | BC5 | $F_1K$ | $F_1P$ | *F* | η2 |
| BW (gram) | 753.36a (± 155.31) | 1706.82b (± 262.54) | 1244.14ab (± 453.82) | 602.88a (± 79.93) | 68.896$^{***}$ | .796 |

BW: Bodyweight; F2K: F2 *Kambro*; F1K: F1 *Kambro*; F1P: F1 *Pelung*; BC5: Broiler Cobb 500.
*: p <0.05; **: p <0.01; ***: p <0.001; Sd: The Standard Deviation is located below the Mean. Mean with different letters in one line is significantly different based on Fisher's LSD and Tukey HSD (p <0.05)

#### Table 5

Frequency of LEPR alleles and genotypes in Broiler Cobb 500, *Pelung* and *Kambro.*

LEPR, leptin receptor gene (NCBI Ref.Seq. **AY048693.1**: g. 127C>A); PCR, polymerase chain reaction; SNP, single nucleotide polymorphism; in the analysed LEPR sequence, SNP were present: g. 127C>A; HW – Hardy–Weinberg equilibrium

| Locus | Frequency | | | | | HW *P*-value |
|---|---|---|---|---|---|---|
| | Allele | | Genotype | | | |
| g. 127C>A | C | A | CC | CA | AA | - |
| | 0.583 | 0.416 | 0.339 | 0.0 | 0.173 | 24.0569 |

HW *P*-value: *P* > 0.05; dF: 1; HW *P*-value (24.0569)> HW *P*-value (*P* (0.05)= 3.84).

#### Table 6

Frequency of SNP g.127C>A in Broiler Cobb 500, *Pelung,* $F_2$ *Kambro* and $F_1$ *Kambro.*

| | *LEPR* | | | | |
|---|---|---|---|---|---|
| | Allele | | Genotype | | |
| | C | A | CC | CA | AA |
| Broiler Cobb 500 | 0.25 | 0.75 | 0.0625 | 0.0 | 0.5625 |
| *Pelung* | 1 | 0 | 1 | 0.0 | 0 |
| F1 *Kambro* | 0.333 | 0.666 | 0.1089 | 0.0 | 0.4356 |
| F2 *Kambro* | 0.666 | 0.333 | 0.4356 | 0.0 | 0.1089 |
| Total | | | 0.8035 | 0.0 | 0.5535 |
