## Supplementary material for "Exon 9 *LEPR* Gene SNP Polymorphism of Hybrid Chickens F_2_ *Kambro* Crossbreeds of ♀ F_1_ *Kambro* with ♂ F_1_ *Kambro*": S1-2019-366819-complete.pdf

### POLIMORFISME *SINGLE NUCLEOTIDE POLYMORPHISM EXON* 9 GEN *LEPR* PADA AYAM HIBRIDA $F_2$ KAMBRO (*Gallus gallus gallus*, Linn.1758)

**SKRIPSI**

Disusun oleh:

**I Wayan Swarautama Mahardhika**

**14/366819/BI/09290**

**FAKULTAS BIOLOGI**
**UNIVERSITAS GADJAH MADA**
**YOGYAKARTA**
**2019**

**POLIMORFISME *SINGLE NUCLEOTIDE POLYMORPHISM EXON* 9 GEN *LEPR* PADA AYAM HIBRIDA $F_2$ KAMBRO (*Gallus gallus gallus*, Linn.1758)**

**SKRIPSI**

Untuk memenuhi sebagian persyaratan guna mencapai gelar Sarjana Sains
Program Studi Biologi

Disusun oleh:

**I Wayan Swarautama Mahardhika**

**14/366819/BI/09290**

Dosen Pembimbing:

**Prof. Dr. Budi Setiadi Daryono, M.Agr.Sc**

**FAKULTAS BIOLOGI**

**UNIVERSITAS GADJAH MADA**

**YOGYAKARTA**

**2019**

#### HALAMAN PENGESAHAN

**POLIMORFISME *SINGLE NUCLEOTIDE POLYMORPHISM EXON* 9 GEN *LEPR* PADA AYAM HIBRIDA $F_2$ KAMBRO (*Gallus gallus gallus*, Linn.1758)**

**SKRIPSI**

Disusun oleh:

**I Wayan Swarautama Mahardhika**

**14/366819/BI/09290**

Telah dipertahankan di depan Tim Penguji pada tanggal 17 Mei 2019 dan dinyatakan telah memenuhi syarat

Yogyakarta, 16 Juli 2019

Fakultas Biologi
Universitas Gadjah Mada

Tim Penguji
Pembimbing I/Penguji I

Prof. Dr. Budi Setiadi Daryono, M.Agr.Sc

Dekan

Prof. Dr. Budi Setiadi Daryono, M.Agr.Sc

Penguji II

Dr. med. vet. Hendry T.S.S.G. Saragih, M.P.

Penguji III

Dr. Slamet Widiyanto, S.Si., M.Sc.

#### PERNYATAAN BEBAS PLAGIASI

Saya yang bertanda tangan di bawah ini:

| | |
|---|---|
| Nama | : I Wayan Swarautama Mahardhika |
| NIM | : 14/366819/BI/09290 |
| Tahun terdaftar | : 2014 |
| Program Studi | : Biologi |
| Fakultas/Sekolah | : Biologi |

Menyatakan bahwa dalam dokumen ilmiah Tugas Akhir/ Skripsi/~~Tesis/Disertasi~~* ini tidak terdapat bagian dari karya ilmiah lain yang telah diajukan untuk memperoleh gelar akademik di suatu lembaga Pendidikan Tinggi, dan juga tidak terdapat karya atau pendapat yang pernah ditulis atau diterbitkan oleh orang/lembaga lain, kecuali yang secara tertulis disitasi dalam dokumen ini dan disebutkan secara lengkap dalam daftar pustaka.

Dengan demikian saya menyatakan bahwa dokumen ilmiah ini bebas dari unsur-unsur plagiasi dan apabila dokumen ilmiah Tugas Akhir/Skripsi/~~Tesis/Disertasi~~* ini di kemudian hari terbukti merupakan plagiasi dari hasil karya penulis lain dan/atau dengan sengaja mengajukan karya atau pendapat yang merupakan hasil karya penulis lain, maka penulis bersedia menerima sanksi akademik dan/atau sanksi hukum yang berlaku.

Yogyakarta, 12 Juli 2019

I Wayan Swarautama Mahardhika

14/366819/BI/09290

#### KATA PENGANTAR

Puji syukur kepada Tuhan Yang Maha Esa atas rahmat-Nya penulis dapat menyelesaikan naskah skripsi yang berjudul **"Polimorfisme *Single Nucleotide Polymorphism Exon* 9 Gen *LEPR* Pada Ayam Hibrida $F_2$ Kambro (*Gallus gallus gallus*, Linn.1758)".** Naskah ini disusun sebagai prasyarat tugas akhir di Fakultas Biologi Universitas Gadjah Mada. Tersusunnya laporan ini tentunya tidak lepas dari bimbingan dan bantuan dari beberapa pihak.

Untuk itu penulis menyampaikan terimakasih kepada:

1. Prof. Dr. Budi Setiadi Daryono, M.Agr.Sc selaku Dekan Fakultas Biologi Universitas Gadjah Mada, Dosen Pembimbing Skripsi, Kepala Gama Ayam Research Team yang selalu memberikan arahan dan nasihat selama masa pendidikan penulis,
2. Rina Sri Kasiamdari S.Si., Ph.D., selaku Wakil Dekan Bidang Akademik dan Kemahasiswaan Fakultas Biologi Universitas Gadjah Mada,
3. Donan Satria Yudha, S.Si., M.Sc. selaku Dosen Pembimbing Akademik yang telah memberikan dukungan, nasihat dan motivasi,
4. Dra. Siti Susanti, S.U. dan Dwi Umi Siswanti, S.Si., M.Sc. selaku dosen pengelola Skripsi,
5. Bapak Suryadi dan Bapak Triyanto yang telah membantu dalam pemeliharaan kandang dan ayam-ayam penelitian,
6. Seluruh rekan Formasigen dan Laboratorium Genetika dan Pemuliaan yang memfasilitasi dan memberikan dukungan moril kepada penulis dalam menyelesaikan penelitian ini,
7. Rekan dan Senior Tim Gama Ayam, Gama Melon dan Tim Pengabdian Kepada Masyarakat Fakultas Biologi UGM yang namanya tidak dapat disebutkan satu persatu,
8. Uyeevolutions, selaku kelompok studi yang senantiasa memberikan pandangan, dukungan moril dan canda tawa disela kepenatan mengerjakan naskah ini

9. Bapak dan Ibu penulis, I Wayan Sudi dan Eka Susiantin yang perannya tidak dapat dipertanyakan lagi,
10. Maria Caroline Samodra atas semangat dan senyumannya mampu memotivasi penulis,
11. Seluruh pihak yang telah berkontribusi dalam penyelesaian penelitian ini.

Laporan ini disusun dengan sebaik-baiknya dan diharapkan dapat memberi manfaat. Adapun kekurangan dan ketidaksempurnaan dari laporan ini hendaknya menjadi pembelajaran untuk penulis di masa mendatang. Terimakasih.

Yogyakarta, 12 Juli 2019

Penulis

### DAFTAR ISI

HALAMAN JUDUL ........ i
SAMPUL DALAM ........ ii
HALAMAN PENGESAHAN ........ iii
HALAMAN PERNYATAAN ........ iv
KATA PENGANTAR ........ v
DAFTAR ISI ........ vii
DAFTAR GAMBAR ........ viii
DAFTAR TABEL ........ ix
DAFTAR LAMPIRAN ........ x
ABSTRAK ........ xi
*ABSTRACT* ........ xii

### DAFTAR GAMBAR

### DAFTAR TABEL

**POLIMORFISME SINGLE NUCLEOTIDE POLYMORPHISM *EXON* 9 GEN *LEPR* PADA AYAM HIBRIDA $F_2$ KAMBRO (*Gallus gallus gallus*, Linn.1758)**

I Wayan Swarautama Mahardhika

(14/366819/BI/9290)

#### ABSTRAK

Implementasi metode T-ARMS PCR dalam deteksi *single nucleotide polymorphisms* (SNPs) gen *LEPR* sampel DNA ayam belum pernah dilakukan. Riset ini bertujuan dalam merancang protokol spesifik deteksi SNPs ekson 9 gen *LEPR* dan mendeteksi ekspresi gen *LEPR* atau SNPs *LEPR* pada sampel DNA ayam Pelung, $F_1$ Pelung, Layer, Broiler Cobb 500, ayam $F_1$ Kambro dan ayam $F_2$ Kambro menggunakan metode T-ARMS PCR. Penentuan derajat korelasi gen *LEPR* terhadap Bobot Tubuh (BT) dan Produtivitas Telur (PT) pada populasi ayam $F_1$ Kambro dan $F_2$ Kambro. Parameter fenotip kualitatif menunjukkan enam kelompok variasi fenotip tersegregasi dibandingkan ayam $F_1$ Kambro. Pertumbuhan bobot ayam $F_2$ Kambro mencapai 753,36 ± 155,31 gram dalam 7 minggu tidak signifikan terhadap ayam $F_1$ Kambro disebabkan adanya *inbreeding depression* (Fx = 25%, LI = 4,925%) dan mutasi transversi alel A *LEPR*. Protokol spesifik deteksi SNPs ekson 9 gen *LEPR* dengan metode T-ARMS PCR dapat mendeteksi mutasi C127A *LEPR* dengan rasio IP : OP 10:1 pmol/µM, konsentrasi template DNA ayam 100 ng/µL dengan temperatur *annealing* 55,7$^\circ$C/30s. Mutasi transversi alel A SNPs ekson 9 *LEPR* terdeteksi pada sampel DNA ayam betina $F_1$ Kambro (80%), ayam jantan $F_2$ Kambro (20%), ayam betina Broiler Cobb 500 (75%). Mutasi tersebut tidak terdeteksi pada sampel DNA ayam Layer, ayam Pelung Blirik Hitam dan ayam $F_1$ Pelung.

Kata Kunci: $F_2$ Kambro, $F_1$ Kambro, ARMS-PCR, *genotyping*, C127A

**POLYMORPHISM OF EXON 9 *LEPR* GENE SINGLE NUCLEOTIDE POLYMORPHISM IN HYBRID CHICKENS $F_2$ *KAMBRO***
**(*Gallus gallus gallus*, Linn.1758)**

I Wayan Swarautama Mahardhika

(14/366819/BI/09290)

#### ABSTRACT

The implementation of the T-ARMS PCR method in the detection of single nucleotide polymorphisms (SNPs) in the *LEPR* gene in chicken DNA samples has never been conducted. This research aims to design a specific protocol for exon 9 *LEPR* gene SNPs detection and detect *LEPR* gene expression or *LEPR* SNPs in *Pelung* chicken samples, $F_1$ *Pelung*, Layer, Broiler Cobb 500, $F_1$ *Kambro* chicken and $F_2$ *Kambro* chicken using the T-ARMS PCR method. Determination of *LEPR* gene correlation degree on Body Weight (BT) and Egg Productivity (PT) in $F_1$ Kambro population and $F_2$ *Kambro*. Qualitative phenotype parameters showed six groups of segregated phenotypes compared to $F_1$ *Kambro* chicken. Growth of $F_2$ *Kambro* chicken weight reached 753.36 ± 155.31 grams in 7 weeks was not significant for $F_1$ *Kambro* chicken due to inbreeding depression (Fx = 25%, LI = 4.925%) and transversion of A *LEPR* allele mutations. Specific protocol detection of exon 9 *LEPR* gene SNPs using the T-ARMS PCR method can detect C127A *LEPR* mutations with IP: OP ratio 10:1 pmol / µM, chicken DNA template concentration of 100 ng / µL with annealing temperature of 55.7° C / 30s. The transversion mutation of C127A of LEPR exon 9 SNP were detected in DNA samples of $F_1$ *Kambro* hens (80%), $F_2$ *Kambro* roosters (20%), Broiler Cobb 500 hens (75%). The mutations were not detected in Layer, *Pelung Blirik Hitam* chicken and $F_1$ *Pelung* populations.

Keywords: $F_2$ *Kambro*, $F_1$ *Kambro*, ARMS-PCR, genotyping, C127A

### BAB I
### PENDAHULUAN

#### A. Latar Belakang

Ayam pedaging dan petelur lokal asli Indonesia mengalami perkembangan yang signifikan dengan peningkatan minat dan keterlibatan komunitas peternak unggas dalam industri peternakan ayam lokal asli Indonesia (Iskandar, 2017). Peningkatan produktivitas dan kualitas daya saing ayam pedaging lokal asli dapat dicapai dengan persilangan selektif galur ayam lokal asli Indonesia. Persilangan selektif bertujuan menghasilkan galur ayam dengan kualitas fenotip tertentu selaras dengan kebutuhan manusia (Das *et al*., 2008; Cheng, 2010; Oldenbroek and van der waaij, 2014; Mariandayani *et al*., 2017; Sudrajat dan Isyanto, 2018).

Gama Ayam Research Team sendiri telah berhasil melakukan persilangan selektif antara Pelung Blirik Hitam dan Broiler Cobb 500 yang menghasilkan ayam hibrida $F_1$ Kambro. Identifikasi fenotip capaian sesuai rancangan seleksi menunjukkan Bobot Tubuh (BT) $F_1$ Kambro, 1.244,14 ± 453,82 gram (p <0,001) unggul signifikan terhadap $F_1$ Pelung BT, 602,88 ± 79,93 gram dalam periode 8 minggu dengan diet pakan standar *ad libitum* (Mahardhika and Daryono, 2019). Persilangan selektif mengimplementasikan konsep genetika Mendelian dan genetika molekular. Penerapan riset genetik mengakomodasi upaya evaluasi dan seleksi generasi parental dalam persilangan selektif dan memungkinkan pemulia menyeleksi dengan akurat setiap hibrida dengan karakter unggul. Dalam upaya menghasilkan galur ayam pedaging lokal dengan pertumbuhan cepat dan karakter fenotip unggul relatif terhadap Broiler Cobb 500 dilakukan investigasi beberapa gen, *LEPR* satu diantaranya.

Gen *leptin receptors* (*LEPR*) terlokalisasi pada neuron sekretoris NeuroPeptide Y (NPY) dan teraktivasi melalui pengikatan aktivator leptin, dihipotesiskan bahwa leptin dan reseptor leptin meregulasi produksi *hypothalamic* NPY (*orexigenic effector*) yang menginhibisi perilaku makan (Schwartz *et al*., 1997; Niv-Spector *et al*., 2005; Abbasi *et al*., 2011). Ayam (*Gallus gallus*) merupakan spesies agrikultural vital dan organisme model dalam bidang biologi perkembangan, karenanya identifikasi leptin otentik ayam dan peran regulasi metaboliknya bersifat substansial (Seroussi *et al.,* 2016).

Penerapan seleksi ayam tipe pedaging dengan pertumbuhan bobot atau karkas tubuh yang cepat mengakibatkan beberapa kelainan fisiologis seperti obesitas. Performa produksi dan kebugaran berkorelasi secara negatif (Martin *et al*., 1990; Pinard *et al*., 1998; Niv-Spector *et al.,* 2005). Seleksi multi-karakter dengan tujuan meningkatkan kebugaran dan produktivitas secara simultan tidak memungkinkan melalui seleksi langsung. Seleksi dengan penanda molekular atau *molecular marker-assisted selection* (MAS) dapat digunakan dan diintegrasikan dalam metode seleksi konvensional dan pendekatan molekular lebih diutamakan dalam pemuliabiakan ayam modern (Li *et al*., 2003; Wang *et al.,* 2006).

Kemajuan dalam bioteknologi molekular memungkinkan diagnosis akurat mutasi gen terkait deformitas atau kelainan genetik. Metode *Amplification Refractory Mutation System Polymerase Chain Reaction* (ARMS-PCR) (Newton *et al.,* 1989) dan tetra primer PCR (Ye *et al.,* 1992) dapat mendeteksi polimorfisme sekuen DNA (Alyethodi *et al.,* 2016). Kombinasi kedua teknik molekular tersebut menghasilkan tetra-primer ARMS PCR (T-ARMS PCR) (Ye *et al.,* 2001). ARMS-PCR membutuhkan jumlah sampel sedikit, cepat dan efektif, efisien dan simultan, sensitivitas dan akurasi tinggi serta konsisten (Peng *et al.,* 2017). Implementasi metode T-ARMS PCR dalam deteksi *single nucleotide polymorphisms* (SNPs) gen *LEPR* sampel DNA ayam belum pernah dilakukan. Riset ini bertujuan dalam merancang protokol spesifik deteksi SNPs ekson 9 gen *LEPR* dan mendeteksi ekspresi gen *LEPR* atau SNPs *LEPR* pada sampel DNA ayam Pelung, $F_1$ Pelung, Layer, Broiler Cobb 500, ayam $F_1$ Kambro dan ayam $F_2$ Kambro menggunakan metode T-ARMS PCR. Penentuan derajat korelasi gen *LEPR* terhadap Bobot Tubuh (BT) dan Produtivitas Telur (PT) pada populasi ayam $F_1$ Kambro dan $F_2$ Kambro.

#### B. Rumusan Permasalahan

Poin penelitian yang diamati adalah sebagai berikut:

1. Parameter fenotip kualitatif dan pertumbuhan bobot ayam $F_2$ Kambro.
2. Protokol spesifik deteksi SNPs ekson 9 gen *LEPR* dengan metode T-ARMS PCR.
3. Ekspresi SNPs *LEPR* pada ayam $F_1$ Kambro, ayam $F_2$ Kambro, ayam Broiler Cobb 500, ayam Layer, ayam Pelung Blirik Hitam dan ayam $F_1$ Pelung.

#### C. Tujuan Penelitian

Tujuan dari penelitian ini adalah untuk:

1. Menentukan parameter fenotip kualitatif dan pertumbuhan bobot ayam $F_2$ Kambro.
2. Menentukan protokol spesifik deteksi SNPs ekson 9 gen *LEPR* dengan metode T-ARMS PCR.
3. Mempelajari ekspresi SNPs *LEPR* pada ayam $F_1$ Kambro, ayam $F_2$ Kambro, ayam Broiler Cobb 500, ayam Layer, ayam Pelung Blirik Hitam dan ayam $F_1$ Pelung.

#### D. Manfaat Penelitian

Penelitian ini memiliki beberapa manfaat diantaranya sebagai berikut:

1. Korelasi antara SNPs ekson 9 gen *LEPR* terhadap bobot tubuh, produktivitas dan kualitas telur ayam $F_1$ Kambro dan ayam $F_2$ Kambro.
2. Panduan deteksi dan seleksi karakter fenotip molekular yang presisi dan konsisten dalam persilangan selektif ayam $F_1$ Kambro sebagai upaya mempersingkat periode persilangan dan meningkatkan efisiensi dan efektivitas persilangan pada generasi selanjutnya.
3. Data pendukung kontruksi *genomic map* mengenai ekspresi SNPs pada ekson 9 gen *LEPR* otentik *Gallus gallus domesticus*.

### BAB II
### TINJAUAN PUSTAKA DAN HIPOTESIS

#### A. Tinjauan Pustaka

##### 1. Karakter populasi ayam $F_1$ Kambro, ayam Pelung dan ayam Broiler

Konsumsi daging ayam tahun 2014 mencapai 4,48 kg/kapita/tahun akumulasi dari total konsumsi ayam ras pedaging, ayam ras petelur afkir dan pejantan serta ayam buras (Suwandi, 2015). Pemenuhan konsumsi produk unggas 67% didominasi oleh ayam ras dan 23% ayam lokal Indonesia, dengan total kontribusi peternakan unggas sebesar 60,73% terhadap produksi pangan hewani nasional (Suprijatna, 2010). Industri peternakan ayam pedaging dan petelur lokal Indonesia mengalami perkembangan pesat dengan meningkatnya minat dan keterlibatan masyarakat dalam industri peternakan ayam lokal atau ayam kampung (Iskandar, 2017). Produksi daging ayam lokal nasional sebesar 8,50 % atau sebesar 284,9 ribu ton dengan tingkat kontribusi 12,86 % terhadap produksi daging ayam nasional (Statistik Peternakan dan Kesehatan Hewan, 2017). Industri peternakan unggas ayam Indonesia masih didominasi ketergantungannya terhadap impor bibit ayam dan *grandparent stock* ayam ras pedaging atau broiler disebabkan masa produksi pendek dan profit yang cepat (Kartasudjana and Supriyatna, 2010; Henuk *et al.*, 2015; Nurfadillah *et al.,* 2018). Ayam lokal memiliki potensi besar sebagai sumber bibit unggul pedaging, petelur dan dwiguna dalam pemenuhan konsumsi pangan hewani (Nataamijaya, 2010; Kartika *et al.,* 2016). Ayam lokal asli Indonesia dikenal juga dengan ayam kampung atau ayam buras (bukan ras, *non-breed chickens*), sementara ayam ras komersil terdiri atas Cobb, Hubbar, Hybro, Isa Hyline dan Hisex (Henuk *et al.*, 2015). Identifikasi keanekaragaman ayam lokal asli Indonesia terkini menghasilkan 34 galur ayam yang terdiri atas Ayunai, Balenggek, Banten, Bangkok, Burgo, Bekisar, Cangehgar, Cemani, Ciparage, Gaok, Jepun, Kampung, Kasintu, Kedu (hitam dan putih), Pelung, Lamba, Maleo, Melayu, Merawang, Nagrak, Nunukan, Nusa Penida, Olagan, Rintit atau Walik, Sedayu, Sentul, Siem, Sumatera, Tolaki, Tukung, Wareng, Sabu, and Semau (Henuk dan Bailey, 2014). Sebanyak 11 galur ayam lokal tergolong sebagai ayam pedaging dan petelur potensial salah satunya

Pelung (Han, 2014; Henuk *et al.*, 2015). Potensi plasma nutfah ayam lokal asli Indonesia tersebut harus dapat dikelola dengan optimal guna menunjang peternakan ayam lokal Indonesia. Plasma nutfah ayam lokal asli Indonesia apabila dikelola dengan baik dapat menjadi solusi bagi pemenuhan konsumsi pangan yang selalu meningkat (Daryono dkk., 2010). Menurut Ningsih dan Prabowo (2017) berbagai permasalahan dihadapi oleh sektor perunggasan salah satunya adalah produktivitas dan daya saing produk perunggasan. Menurut Nurfadillah *et al.* (2018) masalah agribisnis peternakan ayam adalah efisiensi rendah yang mengakibatkan tingginya biaya produksi. Peningkatan efisiensi dan kualitas produk perunggasan ditentukan oleh penyediaan bibit unggul khususnya bibit unggul ayam pedaging lokal Indonesia (Anggitasari dkk., 2016). Peningkatan produktivitas dan kualitas daya saing ayam pedaging lokal dapat ditempuh dengan persilangan selektif galur ayam lokal asli Indonesia. Persilangan selektif bertujuan menghasilkan galur ayam dengan kualitas fenotip tertentu sesuai kebutuhan manusia (Oldenbroek and van der waaij, 2014; Cheng, 2010; Das *et al.,* 2008; Mariandayani dkk., 2017; Sudrajat dan Isyanto, 2018).

Pelung Blirik Hitam memiliki keunggulan yaitu postur dan bobot badan yang unggul dibandingkan ayam kampung lain (Nataamijaya, 2005). Bobot badan ayam pelung jantan dewasa umur 1 tahun dapat mencapai 3,37 kg, sedangkan ayam betina 2,52 kg (Daryono dkk*.,* 2010). Broiler Cobb 500 memiliki produktivitas dan laju pertumbuhan tinggi dalam fase *grower* (7-18 minggu) Broiler Cobb 500 jantan dan betina dapat mencapai bobot 1.599,17 gram dan 1.540,46 gram (Pratama dkk., 2015; Hassan *et al.,* 2016). Model seleksi ayam hibrida Kambro terdiri atas parameter estimasi bobot tubuh, pertumbuhan bobot tubuh, parameter bobot tubuh linear, tingkat mortalitas dan parameter vitalitas dan fenotip (Tamzil *et al.,* 2018; Mahardhika and Daryono, 2019). Pemberdayaan galur ayam pedaging lokal asli Indonesia dapat menunjang ketersediaan sumber pangan masyarakat dan mendukung konservasi plasma nutfah ayam lokal asli Indonesia (Suprijatna, 2010).

Persilangan parental I ♀ Broiler Cobb 500 dengan ♂ Pelung Blirik Hitam menghasilkan filial I Kambro sebanyak 18 ekor terdiri atas 9 jantan $F_1$ Kambro dan 9 betina $F_1$ Kambro (Gambar $1A_{1\text{-}2}$). Persilangan parental I ♀ Pelung Blirik

Hitam dengan ♂ Pelung Blirik Hitam menghasilkan filial I ($F_1$) Pelung sebanyak 22 ekor (Gambar $1B_{1-2}$).

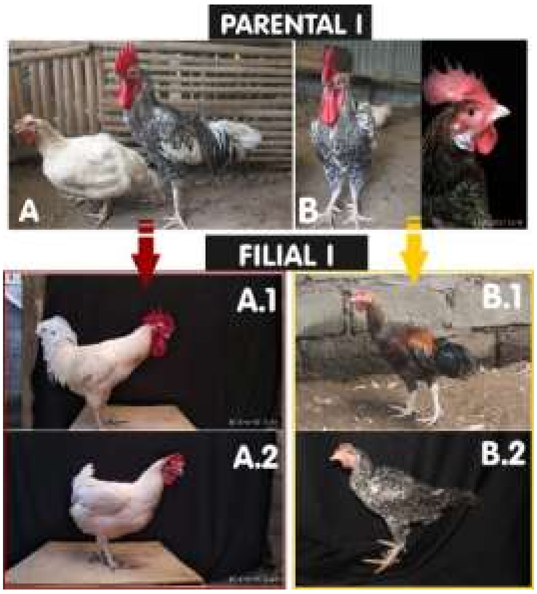

Gambar 1. Persilangan ayam (Dok. Pribadi, 2018)
**Parental I** (A: ♀ Broiler Cobb 500 dan ♂ Pelung Blirik Hitam; B: ♀ Pelung Blirik Hitam dan ♂ Pelung Blirik Hitam) dan **Filial I** (**A.1.**: ♂ Kambro; **A.2.**: ♀ Kambro; **B.1.**: ♂ $F_1$ Pelung; **B.2.**: ♀ $F_1$ Pelung) (Dok. Pribadi, 2017)

Populasi DOC kontrol $F_1$ Pelung dan Broiler Cobb 500 masing-masing sebanyak 22 ekor. Tingkat mortalitas populasi grup III ($F_1$ Kambro), grup II (Broiler Cobb 500) dan grup I ($F_1$ Pelung) secara berurutan sebesar 5,5%, 0% dan 68,2%. Tingkat mortalitas grup I lebih tinggi dibandingkan kelompok II dan III. Kematian ayam termuda grup I pada minggu kedua sementara pada kelompok III terjadi pada minggu keenam. Kemungkinan besar penyebab kematian pada grup I dan grup III disebabkan serangan *infectious coryza* (snot) melalui hasil pengamatan harian. Penyakit *infectious coryza* (snot) disebabkan oleh bakteri gram negatif *Haemophilus paragallinarum* dengan simptom berupa infeksi cepat dan morbiditas tinggi, penurunan produksi telur, *oculonasal conjunctivitis,* pembengkakan wajah dan eksudasi kantung conjuncivital (Ali *et al.,* 2013; Iskandar, 2017). Absennya vaksinasi merupakan perlakuan untuk menguji ketahanan tiap grup ayam. Dari data mortalitas tiap grup ayam tersebut dapat disimpulkan bahwa daya tahan dari kelompok III lebih tinggi dibandingkan kelompok I. Tingkat mortalitas yang tinggi dapat diakibatkan oleh absennya vaksinasi ayam grup I dan III. Grup II telah divaksinasi semenjak menetas oleh produsen DOC. Ayam kampung memiliki ketahanan tubuh lebih baik dibandingkan ayam pedaging lain di wilayah tropis dan ekspresi gen antivirus *Mx+* tertinggi (Diwyanto dan Prijono, 2007; Nuroso, 2010; Kartika *et al.,* 2016; Nurhuda, 2017). Kambro memiliki ketahanan tubuh lebih tinggi mengindikasikan peningkatan mutu genetik ayam lokal melalui persilangan dalam sistem perkandangan semi-intensif didukung oleh beberapa faktor seperti manajemen dan lingkungan. Parameter bobot tubuh linear terdiri atas TA (tinggi ayam), TB (tinggi badan), PB (panjang badan), LB (lebar badan), PPu (panjang punggung), PL (panjang leher), PS (panjang sayap) dan LD (lingkar dada). Parameter vitalitas meliputi TJ (tinggi jengger), PJ (panjang jengger), PK (panjang kepala), LK (lebar kepala), PP (panjang paruh) dan LP (lebar paruh). Parameter fenotip kualitatif meliputi warna bulu leher, warna bulu punggung, warna bulu dada, warna bulu tubuh, warna bulu femur, warna ceker atau *shank*, warna jengger, bentuk jengger dan warna paruh (Wilson, 2010). Parameter fenotip ayam hibrida diidentifikasi sebagai data visual dengan foto berlatar hitam. Rekam data mingguan meliputi pertumbuhan bobot ayam dan panjang femur-tibia (PPa-PBe). Data dianalisis dengan korelasi, regresi, *one way anova* dan *independent sample t-test* menggunakan IBM© SPSS© *Statistics version* 21. Analisis *t-test* digunakan untuk membandingkan rerata bobot tubuh, pertumbuhan tubuh, asupan pakan, konversi pakan dan mortalitas pada dua populasi ayam (Walpole, 1995; Darwati *et al.*, 2016). Korelasi antara panjang femur-tibia dan parameter bobot tubuh linear terhadap pertumbuhan bobot tubuh dianalisis dengan metode korelasi Pearson, regresi linear berganda dan *Analysis of Covariance* (ANCOVA). Parameter fenotip dianalisis dengan metode skoring observasi visual warna berbasis foto.

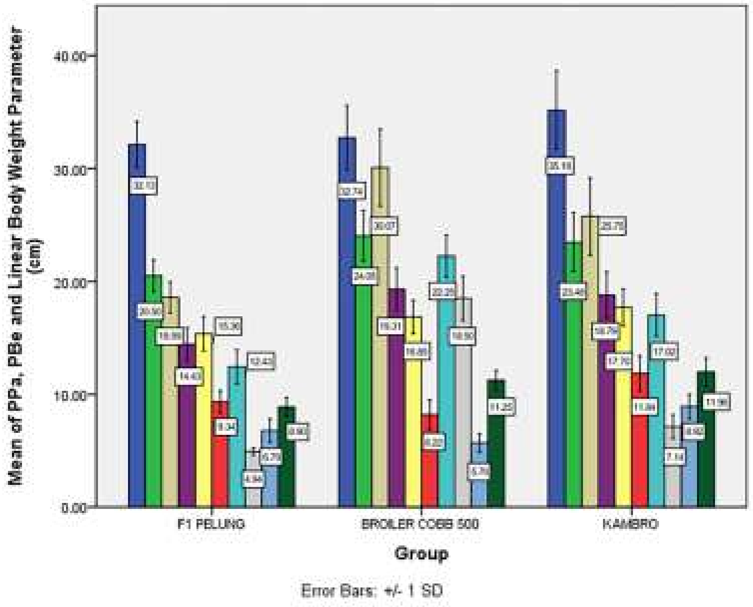

**(A)**

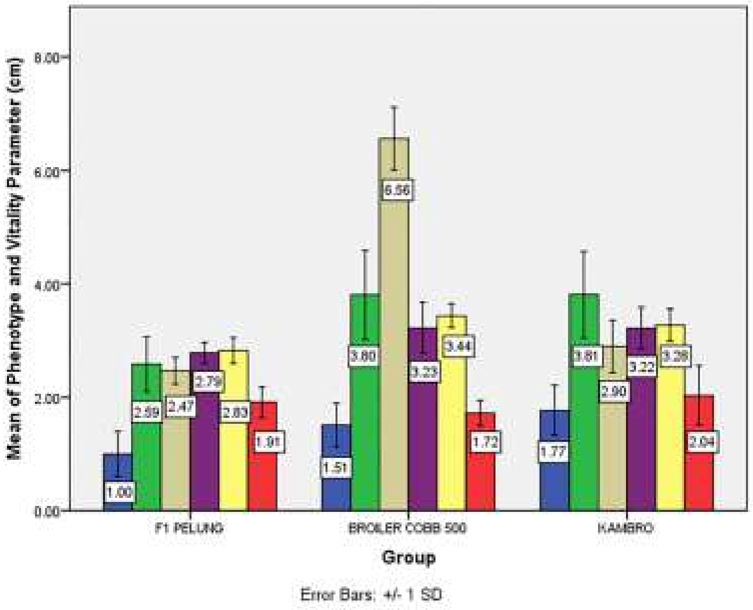

**(B)**

Gambar 2. Rerata PPa, PBe dan parameter bobot tubuh linear
(Olahan Pribadi, 2019)

**(A)** Rerata PPa, PBe dan parameter bobot tubuh linear grup ayam I, II dan III pada umur 8 minggu; **(B)** Rerata parameter vitalitas dan fenotip grup ayam I, II dan III pada umur 8 minggu. Standar deviasi dilambangkan dengan T-bar.

Pada grafik 1 (**A**) tiap parameter dilambangkan warna dengan susunan: **TA** ■; **TB** ■; **PB** ■; **LB** ■; **LD** ■; **PPu** ■; **PS** ■; **PL** ■; **PBe** ■; **PPa** ■

Pada grafik II (**B**) tiap parameter dilambangkan warna dengan susunan: **LP** ■; **PP** ■; **PK** ■; **LK** ■; **TJ** ■; **PJ** ■

Laju koleksi dan penetasan telur Kambro sebanyak 10 hingga 20 butir per minggu selama kurun waktu enam bulan (Desember 2016 s/d Mei 2017). Tingkat produksi telur per minggu sangat rendah sekitar 20 hingga 22 telur pada puncak siklus bertelur Broiler Cobb 500. Daya tetas telur ayam betina Broiler Cobb 500 (± 6 bulan) hanya mencapai 25% / tetasan. Faktor yang mempengaruhi fluktuasi produksi telur Kambro diantaranya faktor asupan nutrisi, tingkat stress, tingkat fertilitas sperma dan fertilitas telur. Broiler petelur produktif pada umur 23 minggu (± 6 bulan) (Rahman *et al.,* 2015). Fluktuasi produktivitas telur betina Broiler Cobb 500 dalam penelitian ini dapat dipengaruhi oleh umur betina. Menurut Hameed *et al.* (2016) berat telur dan daya tetas telur dapat dipengaruhi oleh peningkatan umur betina, penurunan daya tetas telur mencapai 15% pada betina Broiler umur 30 minggu ke atas dengan berat telur kurang dari 60 gram. Faktor utama fluktuasi produksi telur dapat diakibatkan oleh pola diet secara *ad libitum.* Menurut Rahman *et al.* (2015) pola diet *ad libitum* dapat menurunkan produksi telur, daya tetas rendah dan tingkat mortalitas tinggi. Restriksi diet pakan diharuskan untuk membatasi pertambahan bobot tubuh, memaksimalkan produksi telur dan fertilitas betina Broiler Cobb 500 (Rahman *et al.,* 2015).

Tabel 1. Analisis *One Way Anova* PPa, PBe dan BT Grup Ayam I, II dan III

| | Grup Ayam | | | | |
|---|---|---|---|---|---|
| | I (n = 7) | II (n = 22) | III (n =17) | *F* | η2 |
| PPa (cm) | 6,79a (1,03) | 5,69b (0,82) | 8,92ab (1,08) | $55{,}09^{***}$ | 0,719 |
| PBe (cm) | 8,9a (0,82) | 11,25b (0,85) | 11,96ab (1,2) | $22{,}87^{***}$ | 0,515 |
| BT (gram) | 602,88a (79,93) | 1.706,82b (262,54) | 1.244,14ab (453,82) | $62{,}09^{***}$ | 0,743 |

**Ket**: **PPa**: panjang femur; **PBe**: panjang betis; **BT**: bobot tubuh. * = p < 0,05, *** = p < 0,001. Standar deviasi dicantumkan dibawah rerata. Rerata dengan subskrip berbeda dalam kolom yang sama berbeda signifikan (p < 0,05) berdasarkan Fisher's LSD *post hoc* (Olahan Pribadi, 2019)

Pada Tabel 1 ditunjukkan PPa, PBe dan BT tiap grup ayam berbeda secara signifikan (p<0,001). BT berbeda sangat signifikan (p<0,001) pada ketiga grup ayam [F (2, 43) = 62,09, p<0,001, η2 = 0,743]. PPa berbeda sangat signifikan (p<0,001) pada ketiga grup ayam [F (2, 43) = 55,09, p<0,001, η2 = 0,719]. PBe berbeda sangat signifikan (p<0,001) pada ketiga grup ayam [F (2, 43) = 22,87, p<0,001, η2 = 0,515]. Analisis *post hoc* dengan Fisher's LSD mengindikasikan perbedaan signifikan PPa, PBe dan BT pada setiap grup ayam. PPa grup I (M = 6,79, SD = 1,03) signifikan terhadap grup II (M = 5,69, SD = 0,82) dan grup III

(M = 8,92, SD = 1,08). PBe grup I (M = 8,9, SD = 0,82) signifikan terhadap grup II (M = 11,25, SD = 0,85) dan grup III (M = 11,96, SD = 1,2). BT grup I (M = 602,88, SD = 79,93) signifikan terhadap grup II (M = 1.706,82, Sd = 262,54) dan grup III (M = 1.244,14, SD = 453,82). Dapat disimpulkan grup III mengungguli BT, PPa dan PBe mengungguli grup I (Gambar 2A). BT grup III (1.244,14 ± 453,82 gram) mendekati BT grup II (1.706,82 ± 262,54 gram) pada umur 8 minggu. Analisis *one way anova* terhadap PPa, PBe dan BT diperkuat dengan *independent sample t-test* (Tabel 2). PPa grup III (M = 8,92, SD = 1,08) lebih panjang secara signifikan terhadap grup I (M = 6,79, SD = 1,03), t (22) = 4,446, p<0,001). PPa grup III (M = 8,92, SD = 1,08) lebih panjang secara signifikan terhadap grup II (M = 6,79, SD = 1,03), t (37) = 10,62, p<0,001). PBe grup III (M = 8,92, SD = 1,08) lebih panjang secara signifikan terhadap grup I (M = 6,79, SD = 1,03), t (22) = 5,956, p<0,001). PBe grup III (M = 8,92, SD = 1,08) lebih panjang secara signifikan terhadap grup I (M = 6,79, SD =1,03), t (37) = 2,139, p<0,05). BT grup III (M = 8,92, SD = 1,08) lebih tinggi secara seginifikan terhadap grup I (M = 6,79, SD = 1,03), t (21,66) = 9,88, p<0,001). Uji keseragaman *Levene's test* pada BT grup III-I mengindikasikan adanya ketidakseragaman (F = 11,11, *p* = 0,003), sehingga derajat kebebasan disesuaikan dari 22 menjadi 21,66. Rerata BT ayam $F_1$ Kambro umur 8 minggu mengungguli hasil persilangan ayam Sentul dengan rataan bobot per 75 hari sebesar sebesar 896,34 ± 55,46 gram (ayam Sentul jantan) dan 736,00 ± 46,63 gram (ayam Sentul betina) (Solikin dkk., 2016; Sudrajat dan Isyanto, 2018). Mariandayani *et al.* (2013) dilaporkan bahwa bobot badan ayam lokal umur 8 minggu yaitu ayam pelung (jantan 458,23 g dan betina 420,11 g), ayam sentul (jantan 406,36 g dan betina 355,98 g), kampung (jantan 411,56 g dan betina 358,74 g). Dapat disimpulkan bahwa Kambro memiliki BT lebih tinggi dibandingkan ayam pedaging lokal asli murni. Dalam penelitian Hasyim (2015) ayam hibrida persilangan ayam jantan Kampung dan betina Broiler pada umur 12 minggu dapat mencapai bobot 2.335 gram (jantan) dan 1.833 gram (betina). Pertumbuhan bobot tubuh Kambro belum mencapai titik infleksi pada umur 8 minggu sehingga proyeksi pertumbuhan BT Kambro diperkirakan lebih tinggi pada minggu selanjutnya. Titik infleksi merupakan titik maksimum pertumbuhan bobot badan, pada titik tersebut terjadi peralihan perubahan yang asalnya percepatan pertumbuhan menjadi perlambatan. Pertumbuhan dapat terus berlangsung pada minggu selanjutnya karena ayam belum mencapai dewasa kelamin (Sogindor, 2017). Menurut Suprijatna (2010) dewasa kelamin ayam Pelung yaitu 165 hari dengan capaian bobot 12 minggu sebesar 669 gram/ekor.

Tabel 2. Analisis *Independent Sample t-Test* Parameter Bobot Tubuh, Parameter Vitalitas dan Fenotip, PPa - PBe dan BT Grup Ayam I, II dan III pada Umur 8 Minggu

| Parameter | Grup Ayam III-I | | *t* | *df* | Grup Ayam III-II | | *t* | *df* | Grup Ayam II-I | | *t* | *df* |
|---|---|---|---|---|---|---|---|---|---|---|---|---|
| | III | I | | | III | II | | | II | I | | |
| PPa (cm) | 8,92a (1,08) | 6,79b (1,03) | 4,446$^{***}$ | 22 | 8,92a (1,08) | 5,69b (0,82) | 10,62$^{***}$ | 37 | 5,69a (0,82) | 6,79b (1,03) | -2,886$^{*}$ | 27 |
| PBe (cm) | 11,96a (1,2) | 8,9b (0,82) | 5,956$^{***}$ | 22 | 11,96a (1,2) | 11,25b (0,85) | 2,139* | 37 | 11,25a (0,85) | 8,9b (0,82) | 6,404$^{***}$ | 27 |
| BT (gram) | 1.244,1a (453,82) | 602,88b (79,93) | 9,88$^{***}$ | 21,66 | 1.244,14a (453,82) | 1.706,82b (262,54) | -5,691$^{***}$ | 37 | 1.706,82a (262,54) | 602,88b (79,93) | 10,844$^{***}$ | 27 |
| PJ (cm) | 3,81a (0,76) | 2,59b (0,48) | 3,917$^{***}$ | 22 | 3,8a (0,76) | 3,81a (0,785) | 0,029$^{ns}$ | 37 | 3,81a (0,785) | 2,59b (0,49) | 3,854$^{***}$ | 27 |
| TJ (cm) | 1,77a (0,44) | 1b (0,40) | 3,967$^{***}$ | 22 | 1,77a (0,44) | 1,5a (0,39) | 1,925$^{ns}$ | 37 | 1,51a (0,388) | 1b (0,4) | 3,028$^{*}$ | 27 |
| PK (cm) | 2,9a (0,46) | 2,5b (0,24) | 2,310$^{*}$ | 22 | 2,9a (0,46) | 6,56b (0,56) | -21,921$^{***}$ | 37 | 6,56a (0,56) | 2,47b (0,24) | 18,756$^{***}$ | 27 |
| LK (cm) | 3,22a (0,36) | 2,79b (0,187) | 3,003$^{*}$ | 22 | 3,22a (0,363) | 3,23a (0,45) | -0,028$^{ns}$ | 37 | 3,23a (0,45) | 2,79b (0,19) | 3,729$^{***}$ | 24.53 |
| PP (cm) | 3,28a (0,28) | 2,83b (0,23) | 3,693$^{***}$ | 22 | 3,28 (0,28) | 3,44 (0,21) | -2,096* | 37 | 3,44a (0,21) | 2,83b (0,23) | 6,673$^{***}$ | 27 |
| LP (cm) | 2,04a (0,53) | 1,9a (0,27) | 0,571$^{ns}$ | 22 | 2,04a (0,528) | 1,7b (0,23) | 2,507$^{*}$ | 37 | 1,72a (0,22) | 1,91a (0,27) | -1,868$^{ns}$ | 27 |
| TA (cm) | 35,18a (3,45) | 32,13b (2,03) | 2,173$^{*}$ | 22 | 35,18a (3,45) | 32,74b (2,874) | 2,411$^{*}$ | 37 | 32,74a (2,87) | 32,13a (2,03) | 0,517$^{ns}$ | 27 |
| TB (cm) | 23,48a (2,6) | 20,5b (1,4) | 2,825$^{*}$ | 22 | 23,48a (2,61) | 24,05a (2,23) | -0,745$^{ns}$ | 37 | 24,05a (2,23) | 20,5b (1,41) | 3,938$^{***}$ | 27 |
| LD (cm) | 18,79a (2,06) | 14,43b (1,48) | 5,306$^{***}$ | 22 | 18,79a (2,06) | 19,31b (1,91) | -3,916$^{***}$ | 37 | 19,31a (1,91) | 14,43b (1,48) | 8,578$^{***}$ | 27 |
| PPu (cm) | 25,75a (3,42) | 18,58b (1,38) | 5,063$^{***}$ | 22 | 25,75a (3,42) | 30,07a (3,42) | -0,817$^{ns}$ | 37 | 30,073a (3,42) | 18,59b (1,38) | 6,178$^{***}$ | 27 |
| PS (cm) | 7,14a (1,05) | 4,9b (0,33) | 3,211$^{*}$ | 22 | 7,14a (1,05) | 18,51a (1,98) | 1,695$^{ns}$ | 37 | 18,51a (1,98) | 4,94b (0,33) | 2,292$^{*}$ | 27 |
| PL (cm) | 17,02a (1,89) | 12,43b (1,51) | 3,952$^{***}$ | 22 | 17,02a (1,89) | 22,25b (1,87) | 7,954$^{***}$ | 37 | 22,26a (1,87) | 12,43b (1,51) | -2,091$^{*}$ | 27 |
| PB (cm) | 17,7a (1,66) | 15,36b (1,5) | 5,706$^{***}$ | 22 | 17,7a (1,66) | 16,85b (1,49) | -8,653$^{***}$ | 37 | 16,85a (1,49) | 15,36b (1,54) | 12,635$^{***}$ | 27 |
| LB (cm) | 11,8a (1,54) | 9,34b (0,97) | 7,803$^{***}$ | 21,29 | 11,84a (1,54) | 8,2b (1,3) | - 23,109$^{***}$ | 33,23 | 8,22a (1,3) | 9,34b (0,97) | 30,889$^{***}$ | 24.13 |

* = p <0,05, *** = p < 0,001, ns = *non-significant*. Standar deviasi dicantumkan dibawah rerata (Olahan Pribadi, 2019)

Menurut Nurhuda (2017) kombinasi komponen genetik berpengaruh terhadap BT ayam hasil crossbreeding dengan hibrida memiliki performa lebih baik dibandingkan performa parental atau indukannya pada sifat tertentu. Rerata BT Kambro mencapai 1.244, 14 ± 453,82 gram lebih rendah dibandingkan BT Broiler Cobb 500 yang mencapai 1.706, 82 ± 262,54 gram pada umur 8 minggu karena hanya mewarisi 50% komponen genetik ayam Broiler Cobb 500. BT F1 Pelung murni hanya mencapai 602, 88 ± 79,93 gram pada umur 8 minggu.

PPa, PBe dan beberapa parameter bobot tubuh linear berkorelasi dengan bobot tubuh ayam (Ukwu *et al.,* 2014). Parameter bobot tubuh linear terdiri atas panjang ceker (*shank*), lingkar dada, panjang betis, panjang leher, panjang punggung dan panjang paha (Ukwu *et al.,* 2014). Parameter bobot tubuh linear dalam penelitian ini yaitu TA, TB, LB, PL, PS, LD dan PPu. Pengaruh parameter bobot tubuh linear penting di dalam program pemuliaan dan seleksi, selain itu sebagai indikator pertumbuhan bobot tubuh ayam dan daya tarik pasar (Ukwu *et al.,* 2014; Assan, 2015). LD dan PB grup III unggul secara signifikan (p<0,001) terhadap grup II, sementara TA, PL dan LB grup III unggul terhadap grup II (Tabel 2). Peningkatan performa hibrida Kambro terhadap Pelung ditunjukkan oleh hasil signifikan parameter bobot tubuh linear grup III terhadap grup I. Korelasi PPa, PBe dan parameter bobot tubuh linear terhadap BT dirangkum dalam Tabel 3.

Tabel 3. Korelasi Parameter Bobot Tubuh Linear, PPa dan PBe terhadap BT Grup Ayam I, II dan III

| Parameter (cm) | | Grup Ayam I (n=7) | II (n=22) | III (n=17) |
|---|---|---|---|---|
| Parameter Bobot Tubuh | TA | -0,374$^{ns}$ | 0,444$^{*}$ | 0,553$^{*}$ |
| | TB | -0,091$^{ns}$ | 0,380$^{†}$ | 0,633$^{**}$ |
| | LB | 0,344$^{ns}$ | 0,216$^{ns}$ | 0,629$^{**}$ |
| | PB | 0,150$^{ns}$ | 0,005$^{ns}$ | 0,478$^{†}$ |
| | PL | -0,454$^{ns}$ | 0,361$^{†}$ | 0,152$^{ns}$ |
| | PS | 0,792$^{*}$ | 0,179$^{ns}$ | 0,606$^{**}$ |
| | LD | 0,131$^{ns}$ | -0,398$^{†}$ | 0,396$^{ns}$ |
| | PPu | 0,431$^{ns}$ | 0,349$^{ns}$ | 0,299$^{ns}$ |
| PPa | | 0,975$^{***}$ | 0,932$^{***}$ | 0,965$^{***}$ |
| PBe | | 0,298$^{ns}$ | -0,064$^{ns}$ | 0,567$^{*}$ |

† = p < 0,10, *= p < 0,05, **= p < 0,01, ***= p < 0,001. † *very slightly significant*, ns = *non-significant* (Olahan Pribadi, 2019)

Analisis korelasi Pearson’s mengindikasikan korelasi positif signifikan antara PPa dan PBe terhadap BT grup III (PPa $r$ (17) = 0,965, $p$<0,001; PBe $r$ (17) = 0,567, $p$<0,001). Pada grup I dan grup II, BT hanya berkorelasi positif dengan

PPa (grup I $r$ (7) = 0,975, $p$<0,001; grup II $r$ (22) = 0,932, $p$<0,001). Pada grup III TA (0,553), TB (0,633), LB (0,629) dan PS (0,606) berkorelasi positif signifikan (p<0,05) terhadap BT. Pada grup II TA (0,444) berkorelasi positif signifikan (p<0,05) terhadap BT. Pada grup I PS (0,792) berkorelasi positif (P<0,05) terhadap BT. Beberapa parameter bobot tubuh linear pada tiap grup berkorelasi positif lemah terhadap BT sementara PPa berkorelasi positif kuat (r > 0,90) terhadap BT pada setiap grup. Dapat disimpulkan bahwa PPa dapat digunakan sebagai model standar estimasi BT pada ketiga grup ayam. Analisis regresi digunakan untuk memperkuat kesimpulan ditunjukkan pada Gambar 3.

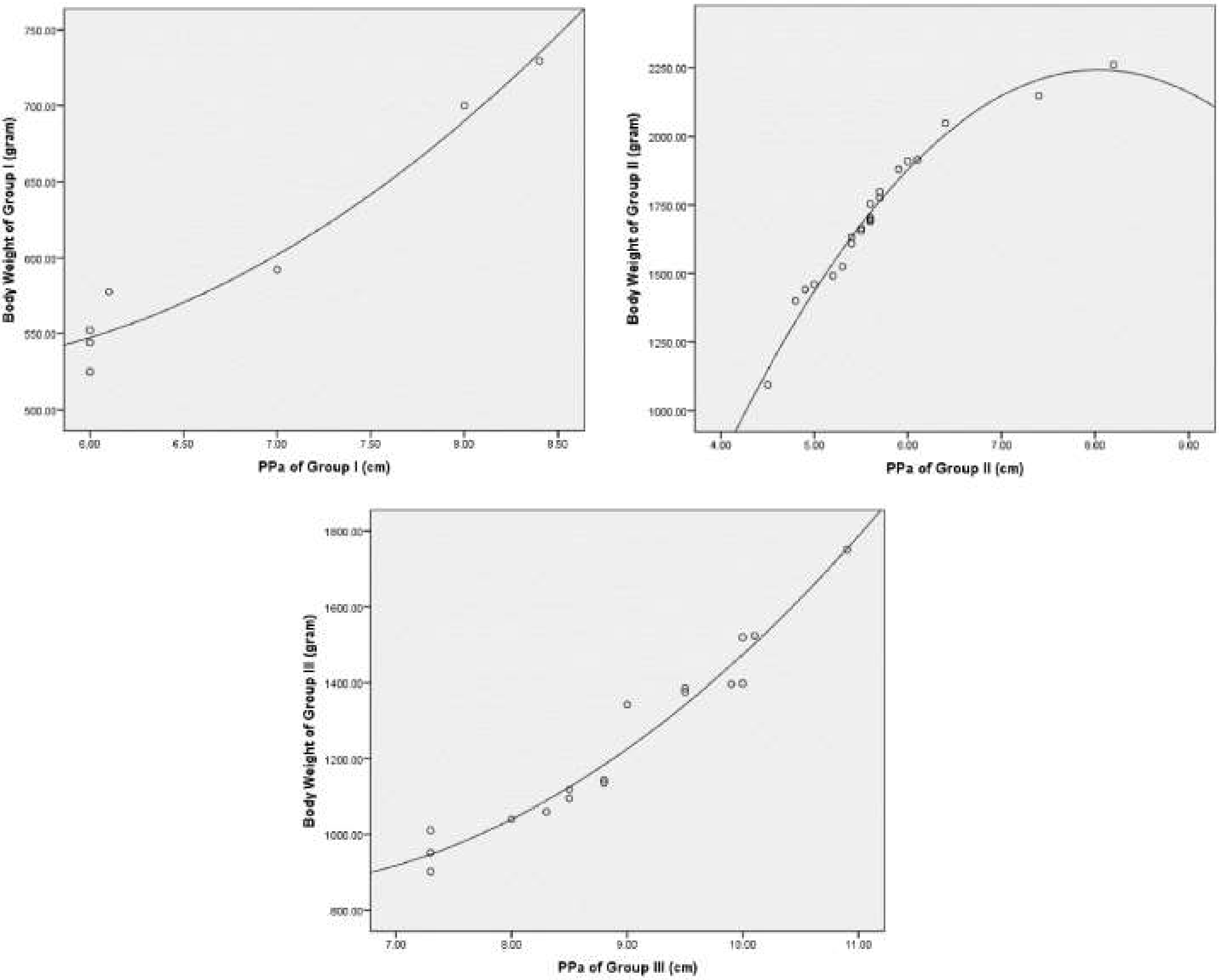

Gambar 3. Model curvilinear quadratic PPa terhadap BT grup ayam I, II dan III (Olahan Pribadi, 2019)

Model regresi non-linear yang digunakan yaitu curvilinear quadratic karena peningkatan nilai $R^2$ lebih tinggi dibandingkan $R^2$ model regresi linear sederhana (Tabel 4). PPa merupakan parameter konstruksi model prediksi yang sesuai digunakan di dalam proyeksi regresi non-linear BT grup ayam I, II dan III.

Tabel 4. Model prediksi bobot tubuh (BT) ayam umur 8 minggu berdasarkan parameter PPa

| Model | Equation | | | | | | | | |
|---|---|---|---|---|---|---|---|---|---|
| | Group I | $R^2$ | sig. | Group II | $R^2$ | sig. | Group III | $R^2$ | sig. |
| Quadratic | 9,23E2±1,63E2*x+16,75*$x^2$ | 0,962 | *** | -3,41E3 + 1,41E3*x ±87,62*$x^2$ | 0,978 | *** | 1,84E3 ± 3,54E2*x+31,73*$x^2$ | 0,956 | *** |
| Linear | 92,81 + 75,17*x | 0,95 | *** | -0,04 + 3E2*x | 0,869 | *** | -6,39E2+2,11E2*x | 0,931 | *** |

*** = P<0,001, variabel x = PPa (Olahan Pribadi, 2019)

Model prediksi BT menurut parameter bobot tubuh linear dengan korelasi positif lemah dianalisis dengan ANCOVA dalam Tabel 5.

Tabel 5. Analisis ANCOVA antara Grup Ayam x Faktor Parameter Bobot Tubuh Linear (FAC1_1)

| Sumber | *Df* | *F* | η2 | *p* |
|---|---|---|---|---|
| Grup | 1 | 7,205 | 0,265 | 0,014 |
| FAC1_1 | 1 | 2,508 | 0,111 | 0,129 |
| Grup*FAC1_1 | 1 | 0,482 | 0,024 | 0,482 |
| Error (within groups) | 20 | | | |

FAC1_1: TA, TB, LB, LD, PL, PS, PPu dan PB; p<0,05 (Olahan Pribadi, 2019)

Analisis ANCOVA antara faktor dan subjek [Grup Ayam (I, II dan III); kovariat: FAC1_1] menunjukkan efek signifikan grup F (1, 20) = 7,205, p = 0,014, η2 = 0,265, sementara FAC1_1, F (1, 20) = 2,508, p = 0,129, η2 = 0,111 tidak menunjukkan efek signifikan, dan tidak ada interaksi antara grup dan FAC1_1, F (1, 20) = 0,482, p = 0,482, η2 = 0,024. ANCOVA memperkuat PPa sebagai parameter model prediksi bobot tubuh ayam $F_1$ Kambro umur 8 minggu. Dalam penelitian Semakula *et al.* (2011) disimpulkan bahwa bobot hidup ayam lokal asli Danau Victoria berkorelasi terhadap ketebalan dada ayam. Dalam penelitian tersebut model prediksi bobot tubuh hidup ayam dan lingkar dada yang sesuai adalah regresi non-linear dengan nilai $R^2$ tertinggi pada model *power* ($0{,}001G^{2.417}$) (Semakula *et al.,* 2011). Dalam penelitian Ukwu *et al.* (2014) disimpulkan bahwa parameter bobot tubuh linear yaitu panjang ceker atau *shank* dapat digunakan sebagai parameter model prediksi bobot tubuh ayam lokal asli Nigeria. Dalam penelitian Mabelebele *et al.* (2017) disimpulkan bahwa ayam Broiler Ross 308 memiliki panjang femur dan tibia mengungguli ayam Venda, ayam lokal asli Afrika Selatan. Fenomena yang sama dapat diamati pada ayam

Pelung dengan PPa lebih pendek dibandingkan PPa Broiler Cobb 500, namun PBe ayam Pelung unggul. Ayam $F_1$ Kambro memiliki PPa dan PBe mengungguli kedua parentalnya (Gambar 2A). Dalam penelitian Mabelebele *et al.* (2017) regresi polinomial fungsi bobot karkas ayam Ross 308 dipengaruhi oleh 97% panjang femur dan 94% tibia, sementara ayam Venda dipengaruhi oleh 89% panjang tibia dan 37% panjang femur. Dalam penelitian ini regresi non-linear quadratic fungsi bobot tubuh ayam Broiler Cobb 500 dipengaruhi oleh 97,8% PPa, ayam $F_1$ Pelung 96,2% PPa dan ayam $F_1$ Kambro 95,6% PPa. Dapat disimpulkan bahwa pertumbuhan panjang PPa Kambro mengikuti pertumbuhan BT Kambro. Fungsi PPa yang lebih tinggi pada Broiler Cobb 500 dibandingkan $F_1$ Pelung dapat diakibatkan oleh sistem pemeliharaan intensif. Ayam Pelung umumnya dipelihara secara ekstensif dengan diet pakan yang variatif, keterbatasan lokomosi dapat menjadi penyebab perlambatan pertumbuhan tulang pada ayam $F_1$ Pelung. Dalam penelitian Henuk *et al.* (2015) ditemukan bahwa sistem ekstensif menurunkan produktivitas ayam lokal asli Indonesia dengan efisiensi pakan sangat rendah dan jangka waktu tumbuh ayam 90 hari/1kg. Pertumbuhan PPa yang mengikuti pertumbuhan BT pada Broiler Cobb 500 dengan sistem ekstensif dan diet pakan tidak ketat dapat berdampak negatif terhadap performa tumbuh ayam ras pedaging "*fast-growing*"(Pauwels *et al.,* 2015). Hasil regresi PPa grup II menunjukkan pertambahan panjang PPa diikuti penurunan BT grup II (Gambar 3). Dapat disimpulkan bahwa peningkatan lokomosi Broiler dipengaruhi BT dan PPa. Diet ketat mengakibatkan penundaan tumbuh sistem muskoskeletal broiler yang menimbulkan stress otot saat berdiri dan bergerak (Paxton *et al.,* 2014). Menurut Shim *et al.* (2012) tulang *fast-growing* broiler umur 6 minggu lebih panjang, lebar, berat, kuat, padat dan tinggi kadar kalsiumnya dibanding *slow-growing* broiler umur yang sama. Menurut Han *et al.* (2015) tibia merupakan bagian terpanjang dan terberat dibandingkan femur dan metatarsus, dengan femur sebagai tulang dengan diameter terpanjang. Performa dan tingkat mortalitas ayam pedaging dipengaruhi oleh struktur tulang. Abnormalitas pertumbuhan tulang dapat dipengaruhi oleh beberapa faktor diantaranya periode pencahayaan. Dalam penelitian van der Pol (2015) disimpulkan pencahayaan minimum menurunkan stress ayam terhadap lingkungan, dimana pencahayaan terang-gelap ekstrim meningkatkan pertumbuhan tulang asimetrik ayam broiler. Fungsi PPa ayam $F_1$ Kambro lebih rendah dibandingkan ayam $F_1$ Pelung dan Broiler Cobb 500 sehingga dapat disimpulkan bahwa sistem semi-intensif dan diet pakan kombinasi yang lebih fleksibel dapat menjadi standar pemeliharaan ayam $F_1$ Kambro.

Penilaian pasar dan seleksi persilangan bergantung terhadap kenampakan parameter fenotip visual (Frame, 2009; Semakula *et al.,* 2011; Assan, 2015) Metode visual dapat secara cepat dan mudah menentukan kualitas dari ayam.

Tabel 6. Parameter fenotip populasi ayam $F_1$ Kambro 8 minggu dengan metode skoring observasi visual

| **Parameter fenotip** | **Karakter** | **Frekuensi gen (%) ♂/♀ (n=17)** | **Lokus** | **Gen** |
|---|---|---|---|---|
| Warna bulu leher | Putih | 100 | *I-i* | $q^I$-$q^i$ |
| Warna bulu punggung | Putih dengan helaian hitam, cokelat dan abu-abu | 100 | *I-i/ E-e+-e* | $q^I$-$q^i$/$q^E$-$q^{e+}$-$q^e$ |
| Warna bulu dada | Putih | 100 | *I-i* | $q^I$-$q^i$ |
| Warna bulu tubuh | Putih dengan helaian hitam, cokelat dan abu-abu | 100 | *I-i/ E-e+-e/B-b* | $q^I$-$q^i$/ $q^E$-$q^{e+}$-$q^e$/$q^B$-$q^b$ |
| Warna bulu femur | Putih | 52,95 | *I-i* | $q^I$- |
| | Putih pola hitam atau abu-abu | 47,05 | *E-e+-e/B-b* | $q^E$-$q^{e+}$-$q^e$/$q^B$-$q^b$ |
| Warna *shank* atau ceker | Putih | 52,95 | *Id- id* | $q^{Id}$/$q^{id}$ |
| | Putih dengan pola hitam atau abu-abu | 41,17 | *Id- id* | $q^{Id}$/$q^{id}$ |
| | Hitam | 5,88 | *Id- id* | $q^{Id}$/$q^{id}$ |
| Warna jengger | Merah | 58,82 | - | - |
| | Merah jambu | 41,18 | - | |
| Bentuk jengger | *Single* | 100 | *P-p* | $q^P$/$q^p$ |
| Warna paruh | Putih gading | 70,58 | - | - |
| | Putih pola hitam | 29,42 | - | - |

(Olahan Pribadi, 2019)

Menurut Navara *et al.* (2012) kenampakan fenotip menjadi penentu produktivitas dan suksesi genetik ayam. LP Grup III tidak signifikan (p>0,05) terhadap grup I (Tabel 2). PJ, TJ dan LK Grup III tidak signifikan (p>0,05) terhadap grup II (Gambar 2B). Warna jengger pada grup III didominasi warna merah terang dengan persentase 58,82% dan warna merah jambu 41,18% (Tabel 6) berbentuk *single* (100%). Menurut Navara *et al.* (2012) warna jengger berkorelasi signifikan positif terhadap fungsi sperma pejantan, namun ukuran jengger berkorelasi signifikan negatif. Temuan ini berbanding terbalik dengan temuan yang mengatakan ukuran jengger berkorelasi signifikan positif terhadap vitalitas, fungsi sperma dan sinyal seksual pejantan (Gebriel *et al.,* 2009; El Ghany *et al.,* 2011; Udeh *et al.,* 2011). Pejantan dominan memiliki dimensi (PJ+TJ) lebih besar dengan warna merah terang tetapi dengan motilitas sperma rendah. Kecenderungan betina memilih pejantan dominan dapat mengakibatkan penurunan kualitas sperma generasi filial (Navara *et al.,* 2012). Menurut Frame (2009) warna jengger berperan sebagai indikator periode bertelur pada betina dengan warna pucat menandakan awal masa bertelur dan afkir sementara merah terang menandakan periode aktif bertelur. Rerata PJ Kambro umur 8 minggu (3,81 ± 0,76 cm) lebih pendek dibandingkan beberapa galur ayam lain seperti *white leghorn* (10-16 cm), *red junglefowl* (6-12 cm) dan broiler (8-14 cm) (Navara *et al.,* 2012). Identifikasi warna jengger, PJ dan TJ pada Kambro menjadi pedoman seleksi indukan, untuk menghindari pejantan dengan motilitas sperma rendah maka dalam persilangan berikutnya akan diseleksi jantan dengan PJ dan TJ terkecil dan betina dengan jengger merah terang. Parameter PK, LK, PP dan LP (Gambar 2B) merupakan indikator pola diet dan laju konsumsi pada ayam dan hubungannya dengan BT telah diselidiki pada berbagai studi (Joller *et al.,* 2018; Fayeye *et al.,* 2013; Fahey *et al.,* 2007; Yakubu *et al.,*2009). Deformitas paruh diketahui untuk mempengaruhi pola makan dan bobot tubuh ayam (Joller *et al.,* 2018) dan dipengaruhi oleh gen *DEGs* (Bai *et al.,* 2014).

Warna paruh didominasi oleh warna putih gading (70,58%) disusul warna putih pola hitam (29,42%). Menurut Frame (2009) pudarnya warna paruh dari putih menjadi kusam atau gading mengindikasikan umur ayam antara 4-6 minggu. Warna ceker atau *shank* pada ayam memiliki lokus *Id-id* dan *W-w* dengan *Id-* mengekspresikan warna kuning atau putih dan *idid* mengekspresikan warna hitam, abu-abu atau kehijauan diekspresikan oleh gen *GRAMD3* pada jaringan dermal *shank* (Sartika dkk., 2010; Xu *et al.,* 2017). Menurut Frame (2009) depigmentasi warna *shank* merupakan indikasi produktivitas telur pada ayam betina selama 15-20 minggu.

Pelung Blirik Hitam memiliki bulu dengan genotipe $Z^BZ^b$ (blirik) dan alel $Z^bZ^b$ (polos) dan Broiler Cobb 500 memiliki bulu dengan genotipe $Z^bW$ (polos) berdasarkan panduan dalam Sartika *et al.* (2016). Warna bulu tubuh pada ayam broiler tergolong dalam warna *dominant white* yang diamati pada ayam *white leghorn* dengan beberapa variasi yaitu *smoky* / keabu-abuan (*I*S*) dan *dun*/ keputihan (*I*D/i*) (Kerje *et al.*, 2004). Kambro jantan (n=9) dan betina (n=9) memiliki frekuensi gen warna bulu tubuh dengan pola helai hitam, cokelat atau abu-abu sebesar 100% ($Z^B Z^b / Z^B W$). Pelung Blirik Hitam dan Broiler Cobb 500 memiliki genotipe ceker atau *shank* yaitu *IdId/Id_* (putih/kuning) dan *idid* (hitam/abu/hijau). Kambro jantan (n=9) dan betina (n=9) memiliki frekuensi gen warna ceker atau *shank* yaitu *IdId/*putih (52,95%), *Idid/*putih pola hitam atau abu-abu (41,17%) dan *idid/*hitam (5,88%). Variasi bulu tubuh dan warna ceker Kambro mengindikasikan adanya segregasi. Menurut Duguma (2006) warna bulu tubuh terang atau putih nilai komersialnya lebih tinggi dan memenuhi standar pasar. Semakula *et al.* (2011) yang menunjukkan bahwa penilaian visual berpengaruh terhadap nilai jual dengan kecenderungan permintaan yang lebih tinggi terhadap ayam lokal asli Uganda. Studi dalam negeri oleh Suprijatna (2010) menunjukkan adanya *niche* pasar ayam lokal asli Indonesia dan kecenderungan masyarakat dalam memilih ayam lokal asli berdasarkan cita rasa yang khas dan kenampakan fenotip. Berdasarkan parameter vitalitas dan fenotip ayam $F_1$ Kambro secara signifikan unggul terhadap Broiler dan Pelung. BT Kambro 1.244,14 ± 453,82 gram secara signifikan ($p<0,001$) unggul terhadap $F_1$ Pelung 602,88 ± 79,93 gram pada umur 8 minggu dengan diet *ad libitum.* Peningkatan performa Kambro signifikan terhadap $F_1$ Pelung berdasarkan parameter bobot tubuh linear, parameter vitalitas, PPa, PBe dan parameter fenotip. Kambro memiliki perpaduan karakteristik fenotip indukannya berdasarkan observasi parameter fenotip. Parameter PPa merupakan model estimasi BT Kambro berdasarkan regresi non-linear quadratic ($r = 0,956$). Perbedaan grup signifikan ($p = 0,014$) terhadap BT dan tidak terdapat interaksi antara grup dan parameter bobot tubuh linear berdasarkan ANCOVA. Tingkat mortalitas Kambro lebih rendah dibandingkan $F_1$ Pelung tanpa vaksinasi dengan sistem pemeliharaan semi-intensif.

#### 2. Persilangan dan Galur

*Trait* atau galur merupakan suatu karakter fenotip determinatif yang terkait dengan suatu individu spesies tertentu (Oldenbroek and van der Waaij, 2014). Domestikasi merupakan kegiatan konversi hewan liar menjadi hewan ternak atau hewan domestik sebagai sumber makanan (Oldenbroek and van der Waaij, 2014). Ayam merupakan hewan hasil domestikasi *Gallus gallus domesticus* yang berasal dari India atau Asia Tenggara pada kurun waktu 6000 BC (Oldenbroek and van der Waaij, 2014). Pemuliabiakan hewan atau *animal breeding* dapat diartikan sebagai upaya peningkatan kualitas karakter fenotip tertentu yang terekspresikan oleh *trait* atau karakter fenotip determinatif pada individu atau spesies hewan domestikasi tertentu dengan mengubah kemampuan genetik *trait* unggulan hewan tersebut melalui persilangan selektif atau *selective breeding* (Oldenbroek and van der Waaij, 2014). Hal ini dilakukan dengan melakukan investigasi mengenai karakter unggulan pada hewan tertentu dan pengukuran performa setiap karakter unggulan yang dikehendaki dapat dilihat pada Gambar 4. Persilangan dilakukan antara individu jantan dan betina yang telah melalui proses seleksi kualitas fenotip tertentu untuk meningkatkan dan mewariskan kualitas fenotip yang diinginkan tersebut pada generasi populasi hibrida selanjutnya. Evaluasi karakter fenotip populasi hibrida dilakukan untuk menentukan derajat peningkatan dan pewarisan fenotip diinginkan dari persilangan yang dilakukan.

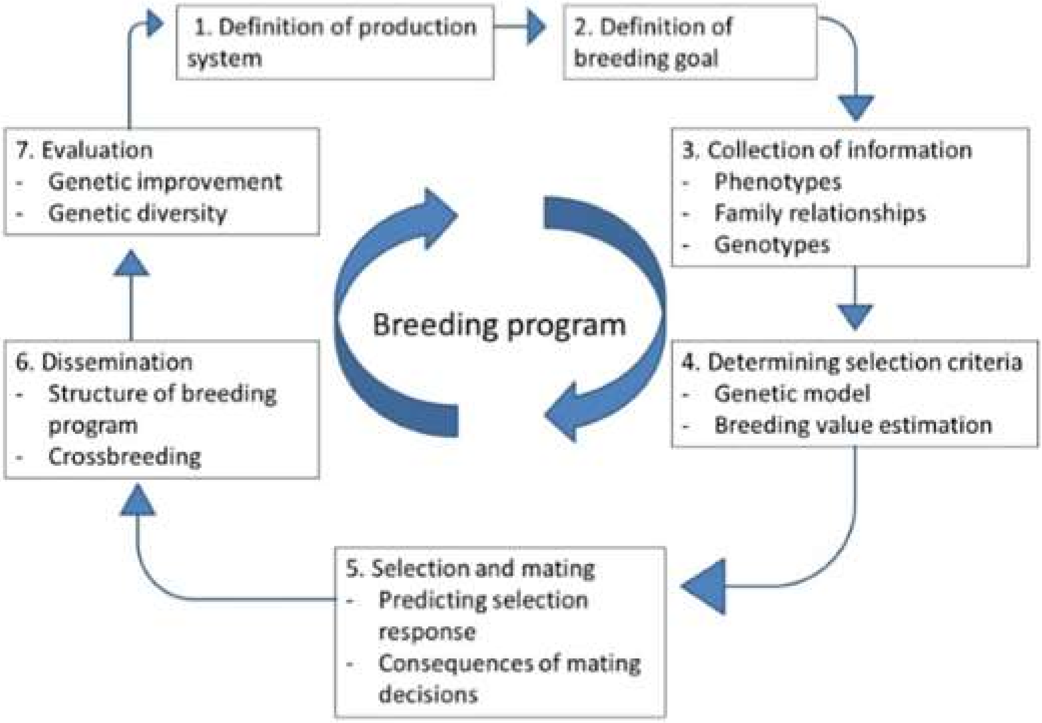

Gambar 4. Alur program *selective breeding* pada persilangan hewan domestikasi. Adaptasi dari: Oldenbroek and van der Waaij (2014)

Ayam $F_1$ Kambro merupakan hibrida hasil persilangan ayam Broiler Cobb 500 dan ayam Pelung Blirik Hitam. Karakter fenotip unggulan yang diharapkan pada ayam $F_1$ Kambro yaitu adanya performa peningkatan bobot tubuh yang cepat, postur tegap, persentase karkas yang lebih tinggi dibandingkan lemak, kemampuan bertelur seperti ayam Pelung dan kemampuan untuk menghasilkan hibrida $F_2$ Kambro yang fertil untuk mendukung proses *selective breeding* dan peningkatan performa *trait* dari kedua parental. Persilangan dapat dilakukan hingga beberapa generasi dengan pertimbangan atau pergeseran karakter tertentu yang hendak dieliminasi atau dipreservasi pada hibrida generasi selanjutnya (Oldenbroek and van der Waaij, 2014). Dalam kegiatan menghasilkan galur baru (*breed*) perlu dilakukan persilangan hingga beberapa generasi hingga mendapatkan hibrida dengan karakter fenotip yang terekspresikan seragam baik dari segi performa, kenampakan dan sejarah seleksi (Oldenbroek and van der Waaij, 2014). Pengaruh *selective breeding* pada hewan domestikasi seperti ayam dapat ditinjau melalui peningkatan performa ayam broiler umur 84 hari yang pada tahun 1957 hanya dapat mencapai bobot 1.907 gram, sementara pada tahun 2001 dapat mencapai 5.958 gram dengan komposisi pakan yang identik (Havenstein *et al.,* 2003). Peningkatan performa akibat *selective breeding* yang sama juga dapat diamati pada ayam Layer dari periode tahun 1950 hingga 1993, ayam Layer dapat mulai bertelur 28 hari lebih cepat dari sebelumnya dengan peningkatan bobot telur sebesar 7 gram dan penurunan kebutuhan nutrisi pakan sebesar 10%. Proses *animal breeding* memiliki beberapa kriteria yang harus dipenuhi sebagai berikut:

1. *Heritable trait* atau kualitas genetik tertentu yang dapat diturunkan contohnya kemampuan bertelur atau postur tubuh yang gagah pada ayam.
2. Latar belakang genetik atau galur yang berbeda dalam satu spesies sehingga dapat dilakukan perkawinan intraspesies atau dalam satu spesies.
3. Seleksi karakter fenotip pada populasi hibrida pada setiap generasi ($F_1$ - $F_n$)
4. Derajat efektivitas persilangan diamati pada tingkat populasi hibrida dengan melakukan perhitungan pergeseran baik kuantitatif dan kualitatif karakter yang diinginkan dari setiap generasi populasi hibrida.
5. Tingkat kesuksesan *selective breeding* dapat diamati pada hasil pengukuran karakter kumulatif populasi hibrida dari keseluruhan generasi.

(Oldenbroek and van der Waaij, 2014)

Dalam kegiatan persilangan hibrida dapat diamati adanya fenomena *inbreeding* atau perkawinan dua individu yang sekerabat (Oldenbroek and van der

Waaij, 2014). *Inbreeding* dapat mengakibatkan penurunan kesehatan hibrida dan kemampuan reproduktif sehingga dapat digolongkan ke dalam efek negatif dalam persilangan. *Inbreeding* dapat menyebabkan penurunan persentase sperma normal dan peningkatan abnormalitas sperma (Oldenbroek and van der Waaij, 2014). *Inbreeding* dapat mengakibatkan efek negatif sebab rendahnya tingkat keragaman genetik atau *genetic diversity* pada populasi hibrida. *Genetic diversity* merupakan tingkat perbedaan genetik antara individu satu spesies, antar generasi atau di dalam suatu generasi tertentu (Oldenbroek and van der Waaij, 2014). *Inbreeding* dapat diukur dengan menentukan seberapa tingginya tingkat *inbreeding coefficient* pada populasi hibrida tertentu. *Inbreeding coefficient* mengindikasikan probabilitas suatu individu menerima alel yang sama dari parentalnya yang sekerabat. *Inbreeding coefficient* memiliki nilai antara 0 (*not inbred*) hingga 1 (*fully inbred*). Perkawinan antara parental yang sekerabat akan meningkatkan tingkat homozigositas alel pada filial akibat *non random mating* (Eldik *et al.*, 2006). Pengaruh *inbreeding* dapat dicegah dengan melakukan *crossbreeding*. *Crossbreeding* merupakan persilangan antara individu hewan satu spesies yang berasal dari galur yang berbeda atau generasi keturunan yang berbeda (Oldenbroek and van der Waaij, 2014). Hibrida hasil *crossbreeding* dari galur atau garis keturunan yang berbeda selama beberapa generasi akan mempertahankan *trait* unggulan tertentu dan meningkatkan heterosis hibrida *crossbred* yang dihasilkan. Heterosis atau *hybrid vigour* merupakan suatu rentang performa hibrida hasil *crossbreeding* dengan satu atau lebih *trait* yang mengungguli rerata performa parentalnya pada *trait* yang diuji (Oldenbroek and van der Waaij, 2014). Heterosis digambarkan pada Gambar 5 yang menunjukkan adanya dominansi oleh satu lokus alel. Dominansi merupakan keadaan ketika suatu lokus alel berada pada kondisi non-aditif yang mengakibatkan nilai lokus heterozigotik dari suatu *trait* berbeda dengan rerata kedua alel homozigotik (Oldenbroek and van der Waaij, 2014). Pada Gambar 5 dicontohkan keadaan dimana nilai lokus homozigotik BB =125 dan bb = 115. Lokus heterozigotik Bb memiliki nilai 122. Efek aditif B terhadap b sebesar (125-115/2) 5, sementara efek dominansi Bb adalah 122-120 (rerata nilai BB dan bb) = 2.

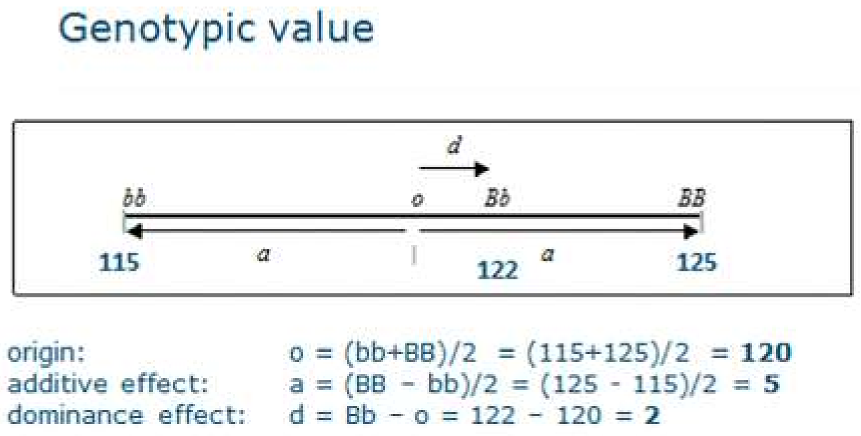

Gambar 5. Skema heterosis pada parental dan hibrida yang mengalami *crossbreeding*. Adaptasi dari: Oldenbroek and van der Waaij (2014)

Heterosis memiliki dampak yang positif terhadap populasi hibrida sebab meningkatkan persentase alel heterozigotik dan mengeliminasi pengaruh alel resesif homozigot pada populasi hibrida. Jumlah heterosis yang diharapkan pada karakteristik tertentu persilangan dua galur ayam tergantung kepada jumlah lokus yang terlibat dan derajat perbedaan persentase lokus yang relevan pada populasi kedua galur (Oldenbroek and van der Waaij, 2014).

##### 3. *Leptin receptor gene* (*LEPR*) *Gallus gallus domesticus*

Evaluasi molekuler melibatkan beberapa marker gen atau *molecular marker-assisted selection* (MAS) yang bertujuan mengidentifikasi alel atau ekspresi gen tertentu dalam populasi ayam (Li *et al*., 2003; Wang *et al*., 2006; Schmid *et al*., 2015). Identifikasi ini menjadi dasar penentuan seleksi dengan tingkat ketepatan tinggi dalam mendeteksi beberapa gen yang berkorelasi signifikan terhadap produktivitas ayam. Penelitian Sun *et al*. (2013) melaporkan bahwa terdapat 14 gen yang berkorelasi terhadap kualitas daging ayam pada ayam $F_2$ hibrida ayam Beijing-You dan Cobb-Vantress. Dalam Nie *et al*., (2005) dilaporkan 12 gen yang berkorelasi terhadap produktivitas daging ayam yaitu *GH, GHR, ghrelin, GHSR, IGF-cn, IGF-co, IGFBP-2, insulin, LEPR, PIT-1, SS* dan *TSH-* $\beta$. Secara khusus terdapat gen yang mengekpresikan leptin dengan implikasi terhadap perilaku makan pada ayam yaitu *LEP* (Zhang *et al*., 1994; Klok *et al.,* 2007; Ohkubo and Adachi, 2008; Ohkubo, 2014; Seroussi *et al.,* 2016; Seroussi *et al*., 2017).

*Leptin receptor gene* (*LEPR*) merupakan gen yang meregulasi translasi reseptor leptin yang memediasi pengikatan leptin dan terintegrasi secara fisiologis termasuk asupan pakan dan metabolisme lipid antara lain persentase dan bobot lemak abdominal pada aves termasuk ayam *broiler* (Wang *et al*., 2006; Bamidele *et al.,* 2012; El Moujahid *et al.,* 2014).

*LEPR* atau disebut juga *OB-R* pada ayam terletak pada kromosom nomor 8 QTL tersusun atas 20 ekson (Wang *et al*., 2006; Adachi *et al*., 2012). Ditemukan korelasi positif antara pembentukan folikel ovarium dengan persentase tingkat leptin dan reseptor leptin pada jaringan folikular melalui steroidogenesis. Ekspresi mRNA *LEPR* tertinggi ditemukan pada lapisan theca folikel kuning (F3-F1) ovarium ayam betina broiler, dimana administrasi eksternal mempengaruhi periode bertelur dengan (a) penundaan penghentian peneluran, (b) penipisan folikel kuning, (c) pengubahan steroidogenesis dan (d) penghilangan apoptosis pada F3-F1 folikel kuning ovarium (Paczoska-Eliasiewicz *et al.,* 2003; Bamidele *et al*., 2012).

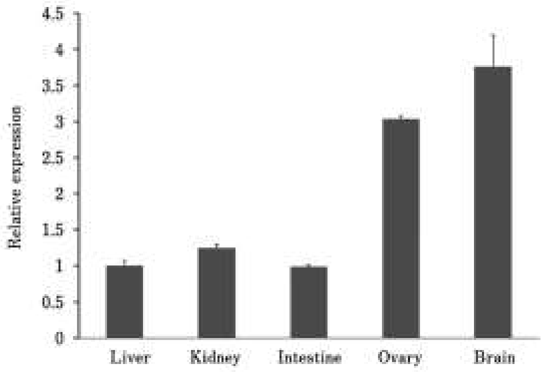

Gambar 6. Ekspresi *LEPR* mRNA pada jaringan tubuh ayam betina umur 5 minggu. **Sumber:** Adaptasi dari Ohkubo (2014). *Relative expression* (weeks): Ekspresi relatif (minggu); *Liver*: Hati; *Kidney*: Ginjal; *Intestine*: Usus; *Ovary*: Ovarium; *Brain*: Otak.

Ekspresi mRNA *LEPR* dalam persentase tinggi terdapat di otak dan ovarium (El Moujahid *et al.,* 2014), persentase kecil terdapat di usus, ginjal dan hati (Mrazova *et al.,* 2012) lihat Gambar 6. Leptin merupakan hormon neuropeptida berukuran 16 kDa dengan yang terekspresikan oleh ekson 9-11 kromosom 8 (Abbasi *et al*., 2011). *Leptin receptor gene* (*LEPR*) terletak di *NeuroPeptide Y* (*NPY*) dan teraktivasi oleh pengikatan leptin yang meregulasi produksi *hypohypothalamic NPY* berkontribusi pada inhibisi perilaku makan unggas (Abbasi *et al*., 2011). cDNA *LEPR* aves pertama kali diidentifikasi pada ayam (Horev *et al*., 2000; Ohkubo *et al*., 2000) dan selanjutnya diidentifikasi pada kalkun (Richards and Poch, 2003) dan bebek (Wang *et al*., 2011). *LEPR* ayam memiliki berat molekul 180 kDa dan dapat diekspresikan pada sel mammalia dan jaringan ayam (Ohkubo *et al*., 2007). Sebagai tambahan, struktur isoform pendek cDNA *chicken LEPR* (*chLEPR*) berhasil dikloning dengan teknik *splicing* menggunakan struktur ekson pendek spesifik antara ekson 19 dan 20 (Liu *et al*., 2007). Gen pengkode *LEPR* pada ayam terdapat pada kromosom nomor 8 (*Accession* No. NC_011472.1) (Dunn *et al*., 2000).

**4. *Single Nucleotide Polymorphisms* (SNPs) dan *Tetra-Primer Amplification Refractory Mutation System-PCR* (T-ARMS PCR)**

Penelitian polimorfisme *LEPR* pada ayam (*Gallus gallus domesticus*) menggunakan *single nucleotide polymorphisms* (SNPs) sebagai pendekatan investigatif. SNPs ayam seperti dijelaskan dalam (Schmid *et al*., 2000; Schmid *et al*., 2005; Schmid *et al*., 2015) menerangkan penggunaan SNPs dalam memahami genotip pada ayam dan unggas domestik. SNPs sendiri merupakan komponen penting konstruksi *Quantitative Trait Loci* (QTL) dengan fungsi aplikatif dalam bidang peternakan dengan memungkinkan analisis genotipik mengenai suatu galur atau *trait*. Informasi genotipik diaplikasikan dalam upaya peningkatan performa dan kualitas suatu galur unggas termasuk ayam. SNPs diterapkan pada genom ayam sebagai spesies domestikasi pertama yang disekuensing (*International Chicken Genome Sequencing Consortium*, 2004). SNPs memudahkan seleksi genomik populasi yang secara aplikatif dapat meningkatkan akurasi estimasi frekuensi genotip suatu *trait* secara efisien tanpa membutuhkan catatan fenotipdari setiap keturunan atau hibrid serta memotong interval generasi *selective breeding* dan memudahkan analisis biodiversitas genotip pada suatu populasi galur ayam. Perkembangan teknik SNPs terbaru dapat menentukan korelasi antara alterasi atau mutasi suatu gen terhadap variasi struktur (SVs, *structural variants*) yang terdiri atas insersi dan delesi (*InDels*) sejumlah pasang basa. SNP juga mendeskripsikan adanya variasi replikasi (CNVs, *Copy Number Variations*) yang terdiri atas sejumlah duplikasi dan delesi segmen DNA berukuran 50 bp atau lebih. CNVs memainkan peran penting di dalam variasi fenotip dan *trait* pada ayam (Schmid *et al.,* 2015). Sebagai contoh CNVs berupa duplikasi parsial 180 kb fragmen gen *PRLR* dan *SPEF2* pada kromosom *Z* dapat menyebabkan terlambatnya pembentukan bulu pada ayam dan jengger *Pea* dipengaruhi oleh CNVs pada gen *SOX5* (Schmid *et al.,* 2015). Dalam studi CNVs terdapat tiga metode yang digunakan untuk mendeteksi CNVs yaitu (1) *medium-high density SNPs, (2) high-density arrays of genomic probes* (*aCGH*) dan *(3) next-generation sequencing* (Schmid *et al.,* 2015). SNPs umumnya berstruktur bi-alelik atau mono-alelik pada region fungsional (promoter dan ekson) dan non-fungsional (Gambar 7) digunakan sebagai marker DNA untuk menyelidiki polimorfisme pada ayam (Schmid *et al.,* 2000). Adanya perubahan basa secara *synonymous* dan *non-synonymous* SNPs merupakan marker penentu dalam mengkonstruksi peta QTL (*Quantitative Trait Loci*).

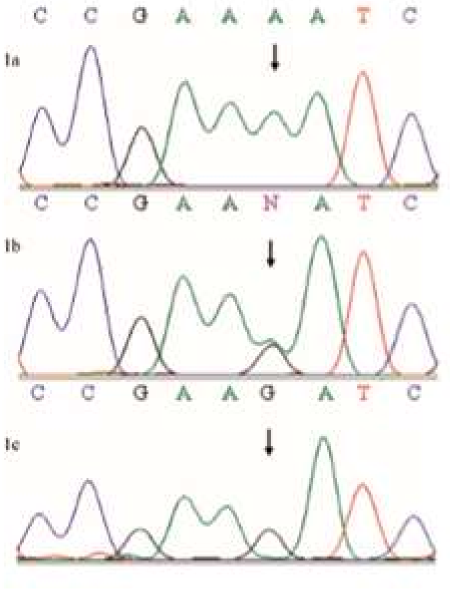

Gambar 7. Sekuen SNPs hasil *alignment* pada tiga individu
1a: homozigot AA; 1b: heterozigot AG; 1c: homozigot GG. Adaptasi dari: Schmid *et al.* (2005)

Penentuan SNPs dilakukan dengan metode PCR sekuen hasil amplifikasi DNA yang dianalisis dengan menggunakan *software* untuk menentukan perubahan frekuensi alel atau gen pada tingkat populasi. Proses untuk menganalisis SNPs pada suatu spesies dapat menggunakan *nuclear SNPs* dan *mitochondrial SNPs* (Schmid *et al.,* 2005). Teknik yang digunakan pada beberapa studi untuk menentukan SNPs diantaranya *oligonucleotide chips,* MALDI-TOF *Mass Spectrometry* dan *pyrosequencing* (Schmid *et al.,* 2005). Analisis SNPs pada broiler dan layer menunjukkan perbedaan sebesar 33% dengan SNPs berupa transisi dari C-T dan A-T. Sebanyak 35% dari 16.630 bp, 26 cDNA SNPs antara broiler dan layer menunjukkan adanya segmen *non-synonymous.* Sebanyak 7.930 bp menunjukkan 55 SNPs dengan range antara 0 sampai 9 per sekuen, menunjukkan satu SNP setiap 144 bp. SNPs yang diamati merupakan transisi pada region pengkode dan region non-pengkode. Pada *coding sequence* persentase transisi sebesar 85% dibandingkan *non-coding region* sebesar 53%. Database genomik ayam memiliki 2,8 juta SNPs menurut laporan *International Chicken Genome Sequencing Consortium*, 2004). Penggunaan *Next Generation Sequencing* (NGS) menunjukkan hasil analisis SNPs pada ayam secara lebih mendetail dan komprehensif. Dalam studi Gheyas *et al.* (2015) disebutkan jumlah SNPs pada ayam sebesar 15 juta SNPs pada genom ayam menggunakan metode NGS. Metode NGS pada genom ayam menunjukkan SNPs dengan potensi variasi fungsional (*pfVars*) yang signifikan terhadap karakter fenotip ayam (Gheyas *et al.,* 2015). Secara garis besar identifikasi SNPs secara signifikan dapat meningkatkan pendeteksian gen terlibat dalan variasi genetik *trait* penting serta memahami fungsi dan perannya dan pemahaman biodiversitas ayam, secara filogenetik terkait lokasi dan spesies yang terlibat dalam domestikasi ayam (Schmid *et al.,* 2005). Identifikasi SNPs pada gen *LEPR* ayam menunjukkan 9 SNPs dengan komposisi 3 SNPs bersifat *synonymous* dan 6 SNPs tergolong intron SNPs dapat dilihat pada Tabel 7 (Nie *et al.,* 2005). Polimorfisme SNPs *LEPR* pada ekson 9 didasari oleh asosiasi yang signifikan antara ekspresi mRNA *LEPR* region tersebut dengan deposisi lemak, metabolisme dan asupan pakan ayam pedaging (Nie *et al.,* 2005; Wang *et al.,* 2006; Bamidele *et al.,* 2012). Beberapa SNPs yang telah dianalisis yaitu AF222783 dengan tipe tri-allelik SNP (T/G/A berukuran 885 nukleotida) pada region sepanjang 1.070 bp *LEPR* (Nie *et al.,* 2005).

Tabel 7. Identifikasi 283 SNPs pada 12 gen terkait pertumbuhan pada ayam (Nie *et al.,* 2005)

| Gene | Chrom[1] | Bps scanned | Primer pairs | Total SNP | SNP numbers[2] | | | |
|---|---|---|---|---|---|---|---|---|
| | | | | | 5'UTR | Syn/non- | Intron | 3'UTR |
| *GH* | 1 | 3945 | 9 | 46 | 4 | 3/2 | 36 | 1 |
| *GHR* | Z | 4007 | 11 | 33 | 0 | 3/5 | 17 | 8 |
| *ghrelin* | 7 | 2536 | 7 | 25 | 1 | 1/1 | 21 | 1 |
| *GHSR* | 9 | 3628 | 7 | 27 | 0 | 9/2 | 25 | 1 |
| *IGF-I* | 1 | 3578 | 10 | 15 | 3 | 0/0 | 1 | 11 |
| *IGF-II* | 5 | 1681 | 3 | 4 | 0 | 1/0 | 3 | 0 |
| *IGFBP-2* | 7 | 4311 | 9 | 35 | 0 | 4/1 | 18 | 12 |
| *insulin* | 5 | 1793 | 4 | 24 | 1 | 0/0 | 22 | 1 |
| *LEPR* | 8 | 1070 | 2 | 9 | 0 | 3/0 | 6 | 0 |
| *PIT-1* | 1 | 2400 | 7 | 23 | 0 | 2/2 | 16 | 3 |
| *SS* | 9 | 944 | 2 | 11 | 0 | 1/2 | 3 | 5 |
| *TSH-β* | 26 | 2004 | 4 | 31 | 0 | 5/0 | 26 | 0 |
| In total | - | 31897 | 75 | 283 | 9 | 32/15 | 194 | 43 |

Penelitian ini menggunakan metode *Tetra Primer Amplification Refractory Mutation System-Polymerase Chain Reaction* (ARMS-PCR). T-ARMS PCR memiliki beberapa keunggulan yaitu kebutuhan sampel sedikit, rapid dan efektif, efisien dan simultan, tingkat sensitivitas dan akurasi yang tinggi dan konsisten (Peng *et al.*, 2017). T-ARMS PCR terdiri atas tiga tahapan yaitu ektraksi DNA genom sampel darah, DNA genom diamplifikasi dengan T-ARMS PCR menggunakan primer spesifik dan penentuan genotip (*genotyping*) produk T-ARMS PCR dengan elektroforesis DNA reguler (Gambar 8) (Peng *et al.,* 2017).

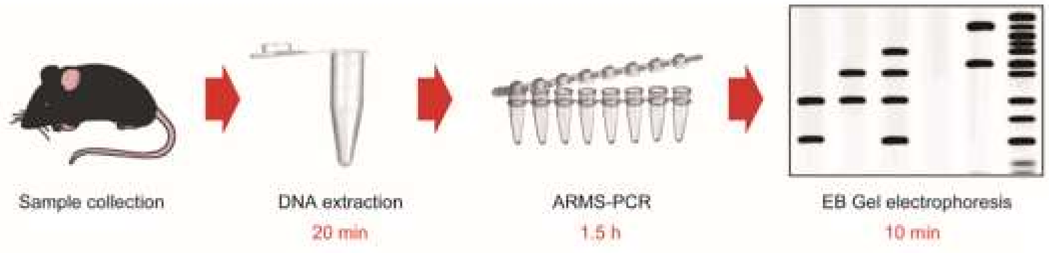

Gambar 8. Skema prosedur *genotyping* dengan ARMS-PCR dan MADGE
Adaptasi dari: Peng *et al.* (2017)

T-ARMS PCR menggunakan empat (tetra) jenis primer yang mengamplifikasi dua alel SNP dalam satu proses PCR (Ye *et al.,* 2001). Primer yang digunakan terdiri atas (Gambar 9) empat jenis yaitu *forward inner primer*, *reverse inner primer, forward outer primer* dan *reverse outer primer*.

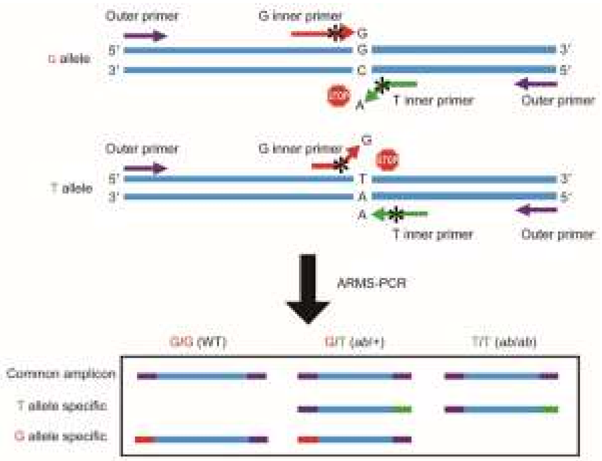

Gambar 9. Skema metode *Tetra Primer Amplification Refractory Mutation System Polymerase Chain Reaction* (ARMS-PCR). Adaptasi dari: Peng *et al.* (2017)
**Ket**: SNP pada gambar merupakan substitusi nukleotida G→T, Dua amplikon alel spesifik dihasilkan dengan menggunakan dua pasang primer, satu pasang mengamplifikasi alel G (panah merah dan hijau) dan satu pasang lain mengamplifikasi alel T (panah ungu). Spesifitas alel ditentukan dengan *mismatch* ujung 3' antara *inner primer* dan template DNA. Peningkatan spesifitas dilakukan dengan *second mismatch* (asteriks *) antara inner primer dan template DNA pada posisi -2 dari ujung 3'. Primer berukuran 26 nukleotida (nt) atau lebih dengan prioritas laju ekstensi dibandingkan hibridisasi. Dua *Outer Primer* berbeda ukuran sehingga amplikon DNA memiliki ukuran berbeda dan dapat dibedakan.

T-ARMS PCR merupakan inovasi dari dua metode yang mendasarinya yaitu *Tetra Primer PCR* dan *Bi-PASA* dengan perbandingan pada Tabel 8. Metode ARMS-PCR menggunakan dua *mismatch* alel spesifik dengan posisi pada basa ujung 3' dan -2 dari ujung 3' dari *inner primer* (Ye *et al.,* 2001).

Tabel 8. Perbandingan ARMS-PCR, *Tetra Primer PCR* dan *Bi-PASA* (Ye *et al.,* 2001)

| | Tetra-primer ARMS-PCR | Tetra-primer PCR | Bi-PASA |
|---|---|---|---|
| Inner primers | | | |
| Allele-specific mismatch | At 3'-terminal base | At centre of primer | At 3'-terminal base |
| Additional mismatch | Yes, at position −2 from 3'-terminus | No | No |
| Length | ~28 bases | ~15 bases | ~20 bases exclusive of tail |
| Tail | No | No | Yes |
| Inner/outer primer ratio | 10 | 1 | 1 |
| Annealing temperature | Constant or touchdown | Higher in early cycles | Constant |

*Mismatch* yang dirancang menggunakan aturan sebagai berikut: *strong mismatch* (G/A atau C/T) dan *weak mismatch* (C/A atau G/T) atau dua *medium mismatch* (A/A, C/C, G/G dan T/T) (Ye *et al.,* 2001). *Outer primer* memiliki panjang nukleotida yang berbeda satu dengan yang lain sehingga amplikon DNA memiliki panjang yang berbeda. Perbedaan ukuran tersebut memungkinkan identifikasi amplikon pada *Microplate Array Diagonal Gel Electrophoresis* (MADGE) (Ye *et al.,* 2001). Desain primer metode ARMS-PCR menggunakan software berbasis web http://primer1.soton.ac.uk (Ye *et al.,* 2001).

#### B. Hipotesis

Hipotesis dari penelitian ini adalah sebagai berikut:

1. Parameter fenotip kualitatif dan pertumbuhan bobot ayam $F_2$ Kambro ekuivalen dengan $F_1$ Kambro dengan depresiasi diakibatkan adanya *inbreeding*.
2. Protokol spesifik deteksi SNPs ekson 9 gen *LEPR* dengan metode T-ARMS PCR mampu mendeteksi keberadaan SNPs *LEPR* otentik *Gallus gallus domesticus*.
3. Ekspresi SNPs *LEPR* pada ayam $F_1$ Kambro, ayam $F_2$ Kambro, ayam Broiler Cobb 500, ayam Layer, ayam Pelung Blirik Hitam dan ayam $F_1$ Pelung akan dapat diukur dan tervisualisasi.

### BAB III
### METODE PENELITIAN

#### A. Waktu dan Tempat Penelitian

Penelitian berlangsung perbulan Desember 2016 hingga Februari 2019 di Pusat Inovasi Agroteknologi (PIAT) UGM, Penetasan Unggas HTN Yogyakarta dan Laboratorium Genetika dan Pemuliaan Fakultas Biologi UGM.

#### B. Bahan dan Alat Penelitian

##### 1. Populasi Ayam

Ayam hibrida $F_1$ Pelung, $F_1$ Kambro dan $F_2$ Kambro dan generasi parental ayam Pelung Blirik Hitam, Broiler Cobb 500 dan ayam Layer dimuliabiakan di Pusat Inovasi Agroteknologi (PIAT) UGM Kali Tirto, Berbah, Sleman, Yogyakarta (Tabel 9).

Tabel 9. Populasi ayam dan penanda individu ayam

| Grup Ayam | | | | | | | |
|---|---|---|---|---|---|---|---|
| **$F_1$ *Pelung* (n=6)** | | **Broiler Cobb 500 (n=4)** | **$F_1$ Kambro (n=6)** | | **$F_2$ Kambro (n=9)** | | **Layer (n=1)** |
| ♀ | ♂ | ♀ | ♀ | ♂ | ♀ | ♂ | ♀ |
| 3p | Pelung Blirik Hitam (11) | 82b | ChipChip (1) | Bjorn (Bj) | CaoCao (5) | Ragnar (3) | Ly |
| 7p | | 90b | ChipChip2 (12) | | Joy (9) | Rollo (4) | |
| 8p | | 92b | ChipChip3 (2) | | Hilda (15) | Clyde (6) | |
| 10p | | 97b | ChipChip4 (14) | | | Igor (7) | |
| | | | ChipChip5 (8) | | | Odin (10) | |
| | | | | | | Satrio (13) | |

♀: betina; ♂: jantan

##### 2. Alat dan Bahan

##### 2.1. PIAT dan HTN Yogyakarta

Timbangan digital (0,01 gram) KrisChef® EK9350H, jangka sorong Verni caliper (0,01 cm), kandang bambu DOC (0-4 minggu), kandang aluminium DOC (4-7 minggu), bohlam 15 watt, koran bekas, wadah minum dan pakan, kalkulator, kamera digital, *metline*, alat tulis, *eggtray*, mesin tetas, kotak pengeram, jerami, kandang persilangan indukan K.4-K.5 Gama Ayam. Beberapa stimulan dan vitamin seperti Egg Stimulant®, VitaChick®, VitaStress®, Tetrachlor® dan

StrongnFit® dan pakan konsentrat standar ayam jenis BR-1 (0-7 minggu) dan pakan AD-II (>7 minggu).

##### 2.2. Laboratorium Genetika dan Pemuliaan UGM

Sampel darah, alkohol 70 %, kertas tisu, es batu, glove, masker, *syringe* 1 ml, EDTA *tube* Vaculab®, larutan chelex, *pipette tip* (biru, kuning dan putih), proteinase K, DTT 0,05 M, buffer TE, PCR tube, agarosa, TAE IX dan 0,5X, FloroSafe™ DNA Stain 1st Base, $ddH_2O$, 2X Kapa Taq DNA Polymerase, primer T-ARMS PCR (FOP, ROP, FIA dan RIC) dan *BenchTop* 100bp Ladder. Alat yang digunakan yaitu *tube* 1,5 ml, mikropipet, *vortex*, *waterbath*, termometer, *microwave*, *microsentrifuge* GyroSpin™, elektroforator Mupid-exU™, BIORAD T100™ PCR Thermal Cycler, inkubator MENMERT, AnalytikJena™ gel imaging system dan GelDoc™ Documentation System.

#### C. Cara Kerja

##### 1. Persilangan ♀ $F_1$ Kambro x ♂ $F_1$ Kambro

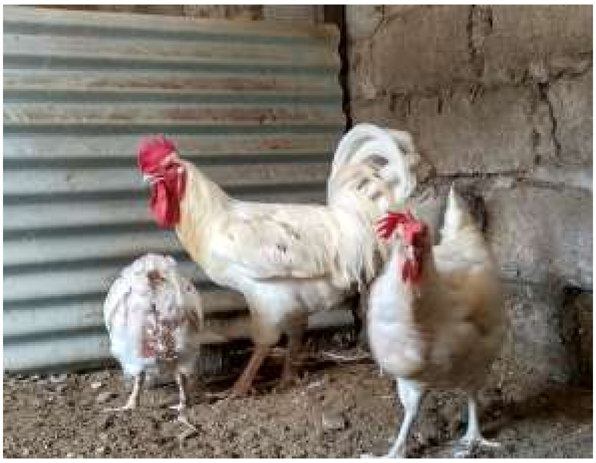

Gambar 10. Persilangan ♀ $F_1$ Kambro x ♂ $F_1$ Kambro (Arsip, 2018).
Persilangan dilakukan di kandang persilangan parental
Gama Ayam berukuran 8 $m^2$

Diet pakan standar AD-II secara *ad libitum* disertai dengan pemberian vitamin melalui air minum berupa *Egg Stimulant®* dan *Tetrachlor®*. Rasio Kambro jantan: betina, 1:2 masing-masing berumur satu tahun. Koleksi telur per hari dengan menggunakan eggtray dan disimpan pada lemari koleksi. Penetasan telur dilakukan per minggu bekerjasama dengan Penetasan HTN Yogyakarta.

**2. Pemeliharaan *Day Old Chicken* (DOC) $F_2$ Kambro**

DOC $F_2$ Kambro hasil tetasan dipelihara secara intensif selama 4 minggu dalam kandang bambu terinsulasi dengan perlengkapan berupa lampu pijar 15 watt, wadah pakan dan minum DOC. Diet pakan standar BR-1 secara *ad libitum* diberikan kepada DOC hingga berumur 7 minggu beserta beberapa vitamin seperti VitaChick® dan VitaStress®. DOC berumur lebih dari 4 minggu dipindahkan ke kandang aluminium dan ayam berumur 7 minggu dipindahkan ke kandang persilangan berukuran 8 $m^2$.

**3. Pengukuran pertambahan bobot ayam dan produktivitas telur $F_1$ Kambro dan $F_2$ Kambro**

Populasi ayam $F_2$ Kambro ditimbang dengan timbangan digital KrisChef® (akurasi .01 gram) per minggu hingga berumur 7 minggu. Produktivitas telur ayam $F_1$ Kambro dan $F_2$ Kambro diukur dengan rekam data koleksi telur per hari dalam periode 4 bulan (April 2018 s/d Juli 2018). Koleksi telur 4 ekor ayam $F_1$ Kambro (1 ± 6 bulan) dan 4 ekor ayam $F_2$ Kambro (± 6 bulan) ditimbang dengan timbangan digital KrisChef© dan morfometri telur diukur dengan jangka sorong Verni caliper. Morfometri telur meliputi lingkar horizontal (LH) dan lingkar vertikal (LV) telur. Dokumentasi dan observasi skoring fenotip visual populasi ayam $F_2$ Kambro meliputi foto berlatar hitam setiap individu ayam $F_2$ Kambro dan telur ayam $F_2$ Kambro dan ayam $F_1$ Kambro.

**4. Analisis molekuler ayam $F_1$ Kambro, $F_2$ Kambro, Broiler Cobb 500 dan $F_1$ Pelung, ayam Layer dan ayam Pelung Blirik Hitam**

**4.1. Koleksi darah**

Sampel darah (1 ml) vena aksiler ayam $F_2$ Kambro, ayam $F_1$ Kambro, ayam Broiler Cobb 500, ayam $F_1$ Pelung, ayam layer dan ayam Pelung Blirik Hitam dikoleksi dengan menggunakan *syringe*. Sampel darah ditransfer dalam *vacutainer* dengan antikoagulan *Ethylenediaminetetraacetic* (EDTA) atau EDTA *tube* merk Vaculab®. EDTA *tube* disimpan dalam lemari es Laboratorium Genetika dan Pemuliaan dengan temperatur -20°C.

**4.2. Isolasi DNA sampel darah ayam**

Isolasi DNA ayam berbasis metode chelex sesuai Ernanto *et al.* (2018) dengan modifikasi. Sampel darah ayam (10 µl) ditransfer ke dalam *microtube*

(1,5 ml) dengan *syringe* lalu dengan mikropipet (tip putih) untuk diencerkan menggunakan TE *buffer* (1 ml, 10 µM, pH 8). Mikrotub disentrifugasi dengan kecepatan 13.000 rpm selama 3 menit. Supernatan dipisahkan dengan menggunakan mikropipet (tip kuning), 200 µl chelex 5%, 18 µl DTT 0,05 M dan 2 µl proteinase K 10mg/ml ditambahkan ke dalam mikrotub dan dihomogenisasi dengan vortex. Inkubasi tahap I dilakukan dengan inkubator merk MENMERT bertemperatur 56℃ selama 1 jam dimana setiap 15 menit dilakukan vortex (total 4 x vortex). Inkubasi tahap II dilakukan dengan waterbath bertemperatur 100℃ selama 8 menit. Mikrotub kemudian disentrifugasi dengan kecepatan 13.000 rpm selama 3 menit. Fase supernatan berisi isolat DNA (±150 µl) ditransfer dengan mikropipet (tip putih) ke mikrotub baru dan dipreservasi dalam temperatur -20℃.

##### 4.3. Desain primer T-ARMS PCR gen *LEPR* ayam berbasis web

Desain primer menggunakan sekuen gen *LEPR* (GenBank *accession number* AY048693.1) dalam Gu *et al.* (2001) dengan panjang 174 bp partial *complete DNA sequence* (CDS) ekson 9 (*Gallus gallus*). Proses BLAST *alignment* antara AY048693.1 dan AF222783.1 bertujuan menemukan *single nucleotide polymorphisms* (SNPs) ekson 9 *LEPR*. Deteksi SNPs *LEPR* dideteksi pada posisi basa 127 yang merupakan transisi basa C/A (C127A) pada Tabel 10.

Tabel 10. Desain primer SNPs *LEPR* dengan Primer BLAST *alignment*

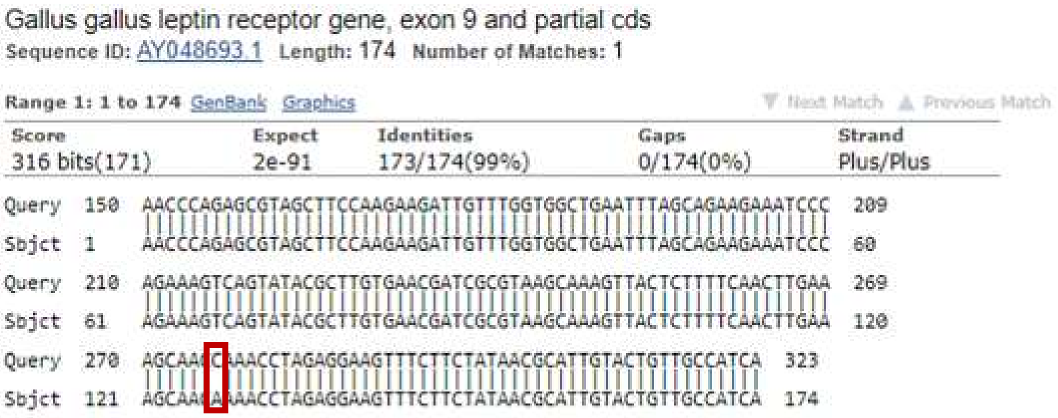

Desain primer berbasis web berdasarkan Peng *et al.* (2017) http://primer1.soton.ac.uk menghasilkan primer pada Tabel 11. Primer terdiri atas empat tipe yaitu *forward outer primer* (FOP) 5’-ATCTATAAAAACAAAACCCAGAGCGTA dan *reverse outer primer* (ROP) 5’TACATATAATTCAGCGTATCTATGATGGC secara spesifik mengamplifikasi *LEPR* dengan amplikon sepanjang 228 bp. Alel mutan A *LEPR* dideteksi dengan *forward inner primer* (FIA) 5'-TTACTCTTTTCAACTTGAAAGCAGCA menghasilkan amplikon sepanjang 114 bp. Alel *wild type* C SNPs *LEPR* dideteksi dengan *reverse inner primer* (RIC) 5'CGTTATAGAAGAAACTTCCTCTAGGTCTG menghasilkan amplikon sepanjang 169 bp. Desain primer T-ARMS PCR diproduksi oleh Integrated DNA Technologies (IDT) dengan perantara PT. GenetikaScience Indonesia.

Tabel 11. Desain primer T-ARMS PCR SNPs *LEPR* ayam berbasis web http://primer1.soton.ac.uk

| Primers | Sequence (5'-3') | Melting Temperature (℃) |
|---|---|---|
| FIA (A allele) | 251 TTACTCTTTTCAACTTGAAAGCAGCA 276 | 55,5 |
| RIC (C allele) | 304 CGTTATAGAAGAAACTTCCTCTAGGTCTG 276 | 55,9 |
| FOP (5' - 3') | 136 ATCTATAAAAACAAAACCCAGAGCGTA 162 | 54,8 |
| ROP (5' - 3') | 363 TACATATAATTCAGCGTATCTATGATGGC 335 | 54,6 |

Product size for A allele: 114 bp
Product size for C allele: 169 bp
Product size of two outer primers: 228 bp

FIA: *forward inner primer* A *allele*; RIC: *reverse inner primer* C *allele*; FOP: *forward outer primer*; ROP: *reverse outer primer*. (Peng *et al.*, 2017)

##### 4.4. *Tetra-Primer Amplification Refractory Mutation System-PCR* (T-ARMS PCR)

T-ARMS PCR mix (25µl) terdiri atas DNA *template*, *nuclease-free water*, KAPA Taq DNA Polymerase dan primer T-ARMS PCR. DNA genomik (50-100 ng) memiliki rentang purifikasi 1-2% (260/280nm) berdasarkan hasil kuantifikasi Spark® Reader spectrophotometer (TECAN®). Primer T-ARMS PCR (FOP, ROP, FIP dan RIP) dilarutkan (1X) dengan TE sebagai larutan stok dan dilarutkan lebih lanjut (10X) untuk digunakan. KAPA Taq DNA Polymerase (200 µM dNTP 1X, 0,5 U/25 µl larutan reaksi) mengandung *buffer* $MgCl_2$ (1,5 mM 1X) dan *stabilizers* (Kapa Biosystems, 2016). Amplifikasi T-ARMS PCR mix menggunakan BIORAD T100™ PCR Thermal Cycler.

##### 4.5. Visualisasi amplikon dan diagnosis SNPs *LEPR* ayam

Amplikon didiagnosis dengan metode elektroforesis gel agarosa dan divisualisasi dengan sinar UV menggunakan mesin AnalytikJena™ gel imaging system dan didokumentasikan dengan GelDoc™ Documentation System. Alel *wild type* C dan alel mutan A dideferensiasikan dengan penanda ukuran amplikon (bp) berdasarkan metode elektroforesis. Gel agarosa 2% dengan melarutkan 0,8 gram agarosa menggunakan 40 ml TBE 1X dalam erlenmeyer, lalu dipanaskan dengan microwave selama 1 menit. Erlenmeyer didinginkan selama 5 menit kemudian ditambahkan FloroSafe™ DNA Stain 1st Base sebanyak 4 µl dan dimasukkan dalam cetakan gel dengan sisir untuk membentuk sumuran. Gel agarosa dibiarkan menjendal (20 menit) lalu ditempatkan dalam wadah elektroforator dan direndam dengan TAE 0,5X. Proses *loading* sampel PCR dan ladder DNA dilakukan dengan volume 5 µl. Elektroforator selanjutnya ditentukan lama dan voltase yang digunakan. Hasil elektroforesis kemudian divisualisasi dengan mesin AnalytikJena™ gel imaging system dan didokumentasikan dengan GelDoc™ Documentation System.

#### D. Analisis Data

Analisis data dalam penelitian ini terdiri atas:

1. Analisis *One Way Anova* dan *Independent Sample t-Test* IBM SPSS Statistics version 21 pertumbuhan bobot tubuh dan parameter kualitas telur populasi ayam $F_1$ Kambro dan $F_2$ Kambro
2. Analisis korelasi Pearson dan regresi linear IBM SPSS Statistics version 21 parameter kualitas telur $F_1$ Kambro dan $F_2$ Kambro
3. Analisis ImageLab v.6.0.1 data dokumentasi elektroforesis sampel PCR DNA ayam $F_1$ Kambro, $F_2$ Kambro, $F_1$ Pelung, ayam jantan Pelung Blirik Hitam, ayam Broiler Cobb 500 dan ayam Layer
4. Perhitungan koefisien *inbreeding* (Fx) dan laju *inbreeding* (LI) $F_2$ Kambro
5. Perhitungan frekuensi genotipe dan frekuensi alel populasi $F_2$ Kambro
6. Analisis deskriptif berbasis perangkat lunak ImageLab V 6.0.1 tes *genotyping* T-ARMS PCR SNPs ekson 9 gen *LEPR* ayam $F_2$ Kambro dengan perbandingan jumlah dan ukuran pita DNA
7. Analisis skoring observasi visual parameter fenotip kualitatif ayam $F_2$ Kambro

### BAB IV
### HASIL DAN PEMBAHASAN

#### A. Parameter Fenotip Kualitatif dan Pertumbuhan Bobot Tubuh Ayam $F_2$ Kambro

Persilangan *inbreeding* generasi parental Kambro (± 1 tahun) ♀ $F_1$ Kambro x ♂ $F_1$ Kambro dengan metode *semi-intensive rearing* menggunakan rasio 1♂ : 2♀. Persilangan tersebut menghasilkan sebelas ekor $F_2$ Kambro viabel terdiri atas 4 betina dan 7 jantan (Gambar 11) tingkat mortalitas hibrida $F_2$ Kambro sebesar 50 % selama periode April hingga Desember 2018. Tingkat mortalitas $F_2$ Kambro lebih tinggi dibandingkan $F_1$ Kambro yang hanya sebesar 5,5%. Kematian termuda populasi $F_2$ Kambro tercatat pada minggu pertama sementara $F_1$ Kambro pada minggu keenam (Mahardhika and Daryono, 2019).

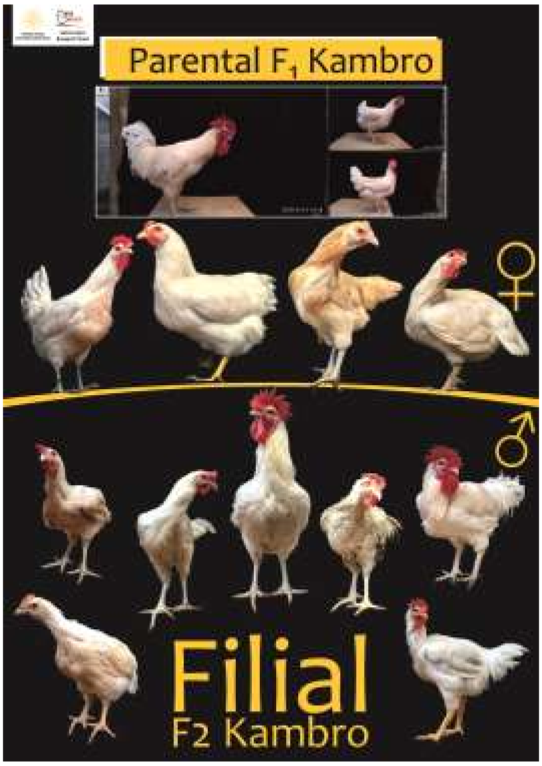

Gambar 11. Persilangan parental $F_1$ Kambro dan populasi $F_2$ Kambro.
♀: 4 ekor betina $F_2$ Kambro; ♂: 7 ekor jantan $F_2$ Kambro (Dok. Pribadi, 2018)

Secara umum penyebab kematian populasi ayam $F_2$ Kambro diakibatkan oleh infeksi coryza atau *snot* berdasarkan observasi harian. Penyakit *infectious coryza* (snot) disebabkan oleh bakteri gram negatif *Haemophilus paragallinarum.*

Beberapa simptom yang ditunjukkan berupa infeksi cepat dan morbiditas tinggi, penurunan produksi telur, *oculonasal conjunctivitis,* pembengkakan wajah dan eksudasi kantung *conjuncivital* (Ali *et al.,* 2013; Iskandar, 2017). Dalam proses pemeliharaan vaksinasi tidak diberikan sebagai bentuk perlakuan untuk mengukur imunitas ayam $F_2$ Kambro. Beberapa individu viabel dalam populasi $F_2$ Kambro dapat secara langsung mewarisi gen *Mx+* dari Pelung Blirik Hitam. Ayam kampung memiliki ketahanan tubuh lebih baik dibandingkan ayam pedaging lain di wilayah tropis dan ekspresi gen antivirus *Mx+* tertinggi (Diwyanto dan Prijono, 2007; Nuroso, 2010; Kartika *et al.,* 2016; Nurhuda, 2017). $F_2$ Kambro memiliki ketahanan tubuh lebih tinggi dibandingkan $F_1$ Pelung yang mencapai 68,2% (Mahardhika and Daryono, 2019). Hal ini mengindikasikan peningkatan mutu genetik ayam lokal melalui persilangan dalam sistem perkandangan semi-intensif didukung oleh beberapa faktor seperti manajemen dan lingkungan.

Dalam Oldenbroek and van der Waaij (2014) depresiasi imunitas pada generasi ayam hibrida dapat disebabkan oleh adanya *inbreeding* atau perkawinan individu sekerabat. Kegiatan persilangan selektif ayam $F_1$ Kambro secara *inbreeding* bertujuan memperkuat pewarisan beberapa *trait* atau karakter unggulan pada generasi $F_1$ Kambro. Pertumbuhan bobot tubuh, parameter bobot tubuh linear, parameter fenotip, parameter vitalitas dan panjang femur-tibia populasi ayam $F_1$ Kambro secara signifikan unggul terhadap populasi ayam $F_1$ Pelung (Mahardhika and Daryono, 2019).

Tabel 12. Analisis BW pada populasi ayam $F_1$K, $F_2$K, $F_1$P and BC5

| | Chicken Group | | | | | |
|---|---|---|---|---|---|---|
| | $F_2$K (n= 11) | BC5 (n = 22) | $F_1$K (n =17) | $F_1$P (n= 7) | *F* | η2 |
| BW (gram) | 753,36a (± 155,31) | 1.706,82b (± 262,54) | 1.244,14c (± 453,82) | 602,88a (± 79,93) | 68,896*** | 0,796 |

BW: *Body Weight*; $F_2$K: $F_2$ Kambro; $F_1$K: $F_1$ Kambro; $F_1$P: $F_1$ Pelung; BC5: Broiler Cobb 500. *: p<0,05; **: p<0,01; ***: p<0,001; Sd: Standar Deviasi terletak dibawah Mean. Mean dengan huruf berbeda dalam satu baris berbeda secara signifkan berdasarkan Fisher's LSD and Tukey HSD (p<0,05) (Olahan Pribadi, 2019)

Dalam Tabel 12 populasi ayam $F_2$ Kambro dapat mencapai bobot tubuh (BW, *body weight*) 753,36 ± 155,31 gram dalam periode 8 minggu. Dalam Mahardhika and Daryono (2019) populasi ayam $F_1$ Kambro dapat mencapai bobot tubuh 1.244,14 ± 453,82 gram, $F_1$ Pelung 602,88 ± 79,93 gram dan Broiler Cobb

500 1.706,82 ± 262,54 gram. Berdasarkan parameter BW populasi ayam $F_1K$, $F_2K$, $F_1P$ and BC5 berbeda secara signifikan [F (3, 53) = 68,896, p<0,001, η2 = 0,796]. Secara signifikan BW BC5 (M = 1.706,82, SD = 262,54, p<0,001) unggul terhadap BW $F_1K$, $F_2K$ dan $F_1P$, namun $F_2K$ (M = 753,36, SD = 155,31) tidak berbeda secara signifikan terhadap $F_1P$ (M = 602,88, SD = 79,93). Grup $F_1K$ (M = 1.244,14, SD = 453,82, p<0,001) unggul secara signifikan terhadap grup $F_2K$.

Capaian BW / 8 minggu grup $F_2K$ lebih unggul dibandingkan beberapa galur ayam hibrida lain. Rerata BW grup $F_2K$ umur 8 minggu (56 hari) mengungguli hasil persilangan ayam Sentul dengan rataan bobot per 75 hari sebesar sebesar 896,34 ± 55,46 gram (ayam Sentul jantan) dan 736,00 ± 46,63 gram (ayam Sentul betina) (Solikin dkk., 2016; Sudrajat dan Isyanto, 2018). Mariandayani *et al*. (2013) dilaporkan bahwa bobot badan ayam lokal umur 8 minggu yaitu ayam pelung (jantan 458,23 g dan betina 420,11 g), ayam sentul (jantan 406,36 g dan betina 355,98 g), kampung (jantan 411,56 g dan betina 358,74 g).

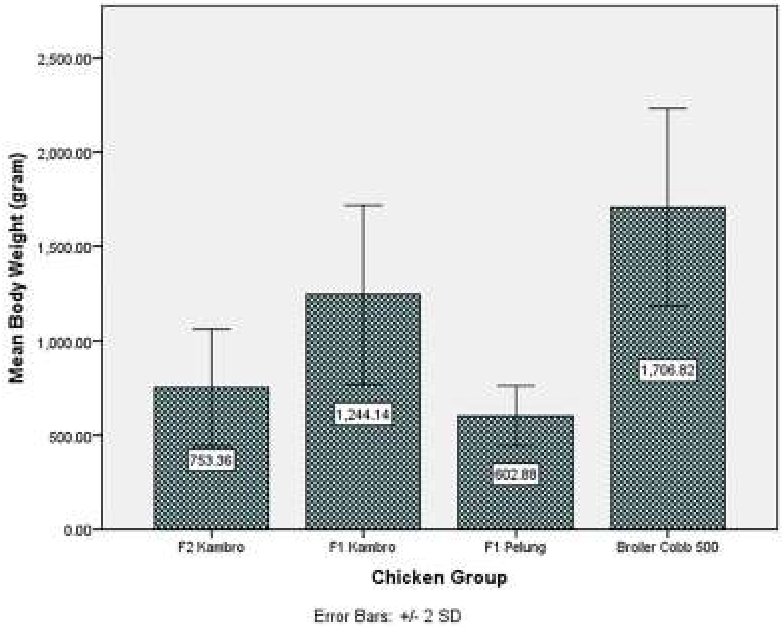

Gambar 12. Bobot Tubuh (BW) $F_2K$, $F_1K$, $F_1P$ dan BC5.
$F_1K$: $F_1$ Kambro; $F_2K$: $F_2$ Kambro; $F_1P$: $F_1$ Pelung; BC5: Broiler Cobb 500
(Olahan Pribadi, 2019)

Capaian BW $F_2$ Kambro mencapai 753,36 ± 155,31 gram lebih rendah terhadap $F_1$ Kambro 1.244, 14 ± 453,82 gram dapat dilihat pada Gambar 12.

Capaian BW $F_2$ Kambro jauh lebih rendah dibandingkan capaian BW Broiler Cobb 500 yang mencapai 1.706, 82 ± 262,54 gram pada umur 8 minggu karena hanya mewarisi 25% komponen genetik ayam Broiler Cobb 500. Pertumbuhan bobot tubuh ayam $F_2$ Kambro belum mencapai titik infleksi pada umur 8 minggu sehingga proyeksi pertumbuhan BW Kambro diperkirakan lebih tinggi pada minggu selanjutnya. Titik infleksi atau dewasa kelamin merupakan titik maksimum pertumbuhan bobot badan, pada titik tersebut terjadi peralihan perubahan yang asalnya percepatan pertumbuhan menjadi perlambatan. Pertumbuhan dapat terus berlangsung pada minggu selanjutnya karena ayam belum mencapai dewasa kelamin (Sogindor, 2017). Menurut Suprijatna (2010) dewasa kelamin ayam Pelung yaitu 165 hari dengan capaian bobot 12 minggu sebesar 669 gram/ekor. Menurut Nurhuda (2017) kombinasi komponen genetik berpengaruh terhadap BW ayam hasil *crossbreeding* dengan hibrida memiliki performa lebih baik dibandingkan performa parental atau indukannya pada sifat tertentu.

Parameter fenotip kualitatif yang diamati pada populasi ayam $F_2$ Kambro meliputi warna bulu leher, warna bulu punggung, warna bulu dada, warna bulu tubuh, warna bulu femur, warna ceker atau *shank*, warna jengger, bentuk jengger dan warna paruh (Wilson, 2010; Liyanage *et al.,* 2015).

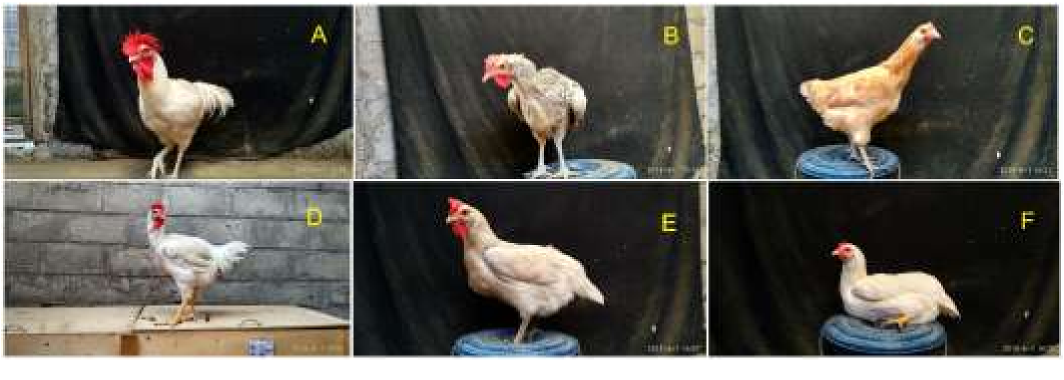

Gambar 13. Golongan fenotip ayam $F_2$ Kambro (Arsip Pribadi, 2018).
A: *Pure White*; B: Blirik Hitam; C: Cokelat-Putih; D: Kuning-Putih; E: Kuning-Hitam; F: Kuning

Observasi parameter fenotip kualitatif ayam $F_2$ Kambro menghasilkan enam golongan fenotip dapat dilihat pada Gambar 13. Parameter fenotip kualitatif merupakan sekumpulan karakter yang secara Mendelian ditentukan oleh satu atau sedikit gen (Habibah, 2018). Dalam Hill (2010) parameter kuantitatif atau *genetic* *complex traits*, dipengaruhi secara poligenik dan sekumpulan faktor non-genetik. Ayam $F_2$ Kambro memiliki enam golongan fenotip yaitu *pure white,* Blirik Hitam, Cokelat Putih, Kuning-Putih, Kuning-Hitam dan Kuning. Golongan A memiliki beberapa determinator yaitu warna bulu tubuh, warna paruh dan warna ceker atau *shank* dominan putih. Golongan B memiliki determinator yaitu warna bulu tubuh blirik hitam dengan warna paruh pola hitam dan warna *shank* putih. Golongan C memiliki determinator warna bulu tubuh cokelat dengan warna paruh dan warna ceker putih. Golongan D memiliki determinator warna bulu tubuh putih dengan warna paruh dan dan warna ceker kuning pada jantan. Golongan E memiliki determinator warna bulu tubuh putih dengan warna paruh kuning dan warna ceker hitam. Golongan F memiliki determinator warna bulu tubuh putih dengan warna paruh dan warna ceker kuning pada betina (Tabel 13).

Tabel 13. Parameter kualitatif ayam $F_2$ Kambro berdasarkan metode skoring observasi visual

| **Parameter fenotipe** | **Karakter** | **Frekuensi gen (%) ♂/♀ (n=11)** | **Lokus** | **Gen** |
|---|---|---|---|---|
| Warna bulu leher | Putih | 81,82 | *I-i* | $q^I$-$q^i$/ $q^I$-$q^I$ |
| | Cokelat | 9,09 | *E-e+-e* | $q^E$-$q^{e+}$-$q^e$ |
| | Putih dengan helaian hitam dan abu-abu | 9,09 | *E-e+-e* | $q^E$-$q^{e+}$-$q^e$ |
| Warna bulu punggung | Putih | 81,82 | *I-i* | $q^I$-$q^i$ |
| | Cokelat | 9,09 | *E-e+-e* | $q^E$-$q^{e+}$-$q^e$ |
| | Putih dengan helaian hitam dan abu-abu | 9,09 | *E-e+-e* | $q^E$-$q^{e+}$-$q^e$ |
| Warna bulu dada | Putih | 81,82 | *I-i* | $q^I$-$q^i$ |
| | Cokelat | 9,09 | *E-e+-e* | $q^E$-$q^{e+}$-$q^e$ |
| | Putih dengan helaian hitam dan abu-abu | 9,09 | *E-e+-e* | $q^E$-$q^{e+}$-$q^e$ |
| Warna bulu tubuh | Putih | 81,82 | *I-i* | $q^I$-$q^i$ |
| | Cokelat | 9,09 | *E-e+-e/B-b* | $q^E$-$q^{e+}$-$q^e$/$q^B$-$q^b$ |
| | Putih dengan helaian hitam dan abu-abu | 9,09 | *E-e+-e/B-b* | $q^E$-$q^{e+}$-$q^e$/$q^B$-$q^b$ |
| Warna bulu femur | Putih | 81,82 | *I-i* | $q^I$- |
| | Cokelat | 9,09 | *I-i* | $q^I$- |
| | Putih pola hitam atau abu-abu | 9,09 | *E-e+-e/B-b* | $q^E$-$q^{e+}$-$q^e$/$q^B$-$q^b$ |
| Warna *shank* atau ceker | Putih | 63,64 | *Id- id* | $q^{Id}$/$q^{id}$ |
| | Kuning | 18,18 | *Id- id* | $q^{Id}$/$q^{id}$ |
| | Putih dengan pola hitam atau abu-abu | 9,09 | *Id- id* | $q^{id}$/$q^{id}$ |
| | Hitam | 9,09 | *Id- id* | $q^{id}$/$q^{id}$ |
| Warna jengger | Merah | 81,81 | - | - |
| | Merah jambu | 18,18 | - | |
| Bentuk jengger | *Single* | 100 | *P-p* | $q^P$/$q^p$ |
| Warna paruh | Putih gading | 72,73 | - | - |
| | Kuning | 27,27 | - | - |
| | Putih pola hitam | 9,09 | - | - |

(Olahan Pribadi, 2019)

Perbandingan jantan dan betina ayam $F_2$ Kambro yaitu 4:7. Dalam populasi ayam $F_2$ Kambro (n = 11) persentase per golongan yaitu A (54,55%), B (9,09%), C (9,09%), D (9,09%), E (9,09%), F (9,09%). Ayam $F_2$ Kambro memiliki bentuk jengger *single* (100%) dengan warna jengger merah (81,81%) dan merah jambu (18,18%). Warna jengger pada ayam $F_1$ Kambro didominasi warna merah terang dengan persentase 58,82% dan warna merah jambu 41,18% berbentuk *single* (100%). Menurut Navara *et al.* (2012) kenampakan fenotipe menjadi penentu produktivitas dan suksesi genetik ayam. Menurut Navara *et al.* (2012) warna jengger berkorelasi signifikan positif terhadap fungsi sperma pejantan, namun ukuran jengger berkorelasi signifikan negatif. Temuan ini berbanding terbalik dengan temuan yang mengatakan ukuran jengger berkorelasi signifikan positif terhadap vitalitas, fungsi sperma dan sinyal seksual pejantan (Gebriel *et al.,* 2009; El Ghany *et al.,* 2011; Udeh *et al.,* 2011). Pejantan dominan memiliki jengger berwarna merah terang tetapi dengan motilitas sperma rendah. Kecenderungan betina memilih pejantan dominan dapat mengakibatkan penurunan kualitas sperma generasi filial (Navara *et al.,* 2012). Menurut Frame (2009) warna jengger berperan sebagai indikator periode bertelur pada betina dengan warna pucat menandakan awal masa bertelur dan afkir sementara merah terang menandakan periode aktif bertelur. Identifikasi warna jengger dan bentuk jengger ayam $F_2$ Kambro menjadi pedoman seleksi indukan, untuk menghindari pejantan dengan motilitas sperma rendah maka dalam persilangan berikutnya.

Warna paruh $F_1$ Kambro didominasi oleh warna putih gading (70,58%) disusul warna putih pola hitam (29,42%). Warna paruh $F_2$ Kambro didominasi oleh warna putih gading (72,73%), kuning (27,27%) dan putih pola hitam (9,09%). Menurut Frame (2009) pudarnya warna paruh dari putih menjadi kusam atau gading mengindikasikan umur ayam antara 4-6 minggu. Warna ceker atau *shank* pada ayam memiliki lokus *Id-id* dan *W-w* dengan *Id-* mengekspresikan warna kuning atau putih dan *idid* mengekspresikan warna hitam, abu-abu atau kehijauan diekspresikan oleh gen *GRAMD3* pada jaringan dermal *shank* (Sartika dkk., 2010; Xu *et al.,* 2017). Menurut Frame (2009) depigmentasi warna *shank* merupakan indikasi produktivitas telur pada ayam betina selama 15-20 minggu.

Pelung Blirik Hitam memiliki bulu dengan genotipe $Z^BZ^b$ (blirik) dan alel $Z^bZ^b$ (polos) dan Broiler Cobb 500 memiliki bulu dengan genotipe $Z^bW$ (polos) berdasarkan panduan dalam Sartika *et al.* (2016). Warna bulu tubuh pada ayam broiler tergolong dalam warna *dominant white* yang diamati pada ayam *white leghorn* dengan beberapa variasi yaitu *smoky* / keabu-abuan (*I*S*) dan *dun* / keputihan (*I*D/i*) (Kerje *et al.*, 2004). Ayam $F_1$ Kambro memiliki frekuensi gen warna bulu tubuh dengan pola helai hitam, cokelat atau abu-abu sebesar 100% ($Z^BZ^b/Z^BW$). Dalam populasi ayam $F_2$ Kambro frekuensi gen blirik hitam (9,09%), cokelat (9,09%) dan putih (81,81%) menandakan adanya segregasi alel dalam kerangka persilangan galur ayam Kambro. Dalam Habibah (2018) terdapat asosiasi positif antara ekspresi warna bulu putih (*recessive white*) pada ayam $F_2$ Golden Kamper terhadap mutasi fragmen gen *cTYR* dalam bentuk insersi sekuen lengkap retroviral intron 4.

Pelung Blirik Hitam dan Broiler Cobb 500 memiliki genotipe ceker atau *shank* yaitu *IdId/Id_* (putih/kuning) dan *idid* (hitam/abu/hijau). Ayam $F_1$ Kambro memiliki frekuensi gen warna ceker atau *shank* yaitu *IdId/*putih (52,95%), *Idid*/putih pola hitam atau abu-abu (41,17%) dan *idid*/hitam (5,88%). Ayam $F_2$ Kambro memiliki frekuensi gen warna ceker atau *shank* yaitu *IdId/*putih (63,64%), *Idid/*kuning (18,18%), *Idid*/putih pola hitam atau abu-abu (9,09%) dan *idid*/hitam (9,09%). Variasi bulu tubuh dan warna ceker ayam $F_2$ Kambro mengindikasikan adanya segregasi.

Menurut Duguma (2006) warna bulu tubuh terang atau putih nilai komersialnya lebih tinggi dan memenuhi standar pasar. Semakula *et al.* (2011) yang menunjukkan bahwa penilaian visual berpengaruh terhadap nilai jual dengan kecenderungan permintaan yang lebih tinggi terhadap ayam lokal asli Uganda. Studi dalam negeri oleh Suprijatna (2010) menunjukkan adanya *niche* pasar ayam lokal asli Indonesia dan kecenderungan masyarakat dalam memilih ayam lokal asli berdasarkan cita rasa yang khas dan kenampakan fenotipe. Berdasarkan parameter fenotipe kualitatif ayam $F_2$ Kambro lebih unggul terhadap Broiler dan Pelung.

#### B. Parameter Kualitas Eksterior, Produktivitas Telur dan Fenotip Telur Ayam $F_2$ Kambro

Produktivitas telur merupakan aspek kuantitatif yang penting dalam kegiatan persilangan selektif. Produktivitas telur yang baik dapat mendukung kegiatan seleksi dan persilangan dalam upaya membentuk suatu galur ayam dengan fenotip tertentu. Kegiatan pendataan telur berlangsung selama ± 270 hari dengan menggunakan masing-masing 4 ekor betina pada grup $F_1$ Kambro ($F_1$K) dan $F_2$ Kambro ($F_2$K). Produktivitas telur dapat diukur dengan menggunakan *Hen Day Production* (HDP) yaitu persentase jumlah produksi telur harian per jumlah betina (Setiawati dkk., 2016). Produktivitas telur 4 ekor ayam betina $F_2$K (± 6 bulan) mencapai 66 butir telur (HDP, 16,5%) dan 4 ekor ayam betina $F_1$K (1 ± 6 bulan) mencapai 96 butir telur (HDP, 24%) yang merangkap total produksi telur 162 butir telur/ ± 270 hari. Berdasarkan data yang diperoleh maka HDP galur ayam Kambro yaitu 20,25%.

Dengan memperhitungkan faktor *inbreeding depression* maka nilai heterosis dapat dihitung pada generasi $F_2$K. Heterosis merupakan lawan dari *inbreeding depression* yaitu nilai peningkatan performa suatu karakter hibrida terhadap indukannya (Oldenbroek and van der Waaij, 2014). Heterosis pada $F_2$K sebesar -31,25% ((16,5-24)/24*100) sehingga dapat diasumsikan bahwa terdapat *inbreeding depression* terhadap penurunan produktivitas telur dalam kondisi kandang dan pemeliharaan yang identik antar grup $F_1$K dan $F_2$K. Hal ini tidak dapat dipastikan sebab individu tiap grup terpaut jauh yaitu 52 minggu sehingga dapat diasumsikan pula bahwa grup $F_2$K belum mencapai titik infleksi dan masa puncak bertelur. Dalam penelitian Sogindor (2017) dijelaskan bahwa produksi telur pada ayam dimulai ketika ayam mencapai dewasa kelamin (titik infleksi).

Produksi telur pada ayam *breeder* dimulai pada saat ayam berumur 24 minggu. Produksi tersebut dapat digambarkan ke dalam suatu kurva. Secara matematis, kurva produksi ini dapat dibagi ke dalam 3 tahap, yaitu : awal produksi - puncak (peningkatan kemiringan), puncak produksi, dan puncak-akhir produksi (penurunan kemiringan). Pada permulaan produksi telur, persentase produksi *hen day* sekitar 5 %. Persentase tersebut meningkat dengan cepat pada 8 minggu Pada saat ayam berumur 31-32 minggu, produksi telur mencapai puncaknya dengan persentase produksi *hen day* lebih dari 80 %. Produksi telah mencapai puncaknya apabila selama 5 hari berturut-turut produksi telur tidak meningkat. Setelah mencapai puncaknya, persentase produksi *hen day* menurun secara konstan dengan laju penurunan sebesar 1% per minggu (Anang dkk., 2007) Dalam pemeliharan ayam $F_1$K dan $F_2$K diterapkan sistem *litter cage* dengan diet pakan standar secara *ad libitum*. Dalam penelitian Setiawati dkk. (2016) diketahui bahwa antara sistem pemeliharaan *cage* dan *litter cage* tidak berkorelasi signifikan terhadap produktivitas telur dan kualitas telur ayam.

Produktivitas telur ayam Kambro yang mencapai 20,25% secara deskriptif unggul dibandingkan generasi parentalnya Broiler Cobb 500 pada minggu ke-24. Produktivitas telur (HDP) ayam *parent stock* Broiler Cobb 500 umur 24 minggu hanya mencapai 1,45% jauh lebih rendah dibandingkan standar Cobb-Vantress yaitu 5% (Anang dkk., 2007). Dalam Anang dkk. (2007) ditemukan bahwa masa puncak produksi *parent stock* Broiler Cobb 500 adalah minggu ke-31 sebesar 83,61%. Dengan memperhitungkan adanya pewarisan alel asosiatif terhadap produktivitas telur maka ayam Kambro memiliki puncak masa bertelur yang sama dengan tetuanya Broiler Cobb 500.

Produktivitas telur ayam Kambro (162 butir/± 270 hari; 20,25%) secara deskriptif lebih rendah dibandingkan ayam lokal asli yaitu Pelung, Kampung dan Lurik. Dalam penelitian Darwati (2000) dilaporkan bahwa produktivitas telur ayam Pelung mencapai 20,36% - 43,37% dengan rataan 31,28 ± 6,34%, sementara ayam Kampung mencapai 26,05 - 47,90% dengan rataan 33,8 ± 7,49%. Dalam Depison (2009) produktivitas telur ayam Lurik mencapai 200 - 250 butir telur/tahun dengan hibrida Pelung x Lurik dapat mencapai 39,57%. Produktivitas telur ayam Kambro secara deskriptif lebih rendah dibandingkan produktivitas ayam $F_1$ Kamper yang mencapai 140,37 butir telur/300 hari, pelung 56,40 butir telur/300 hari dan layer 195,07 butir/300 hari (Ernanto, 2017). Produktivitas telur ayam Lohmann LSL-Classic Layer pada masa puncak bertelur dapat mencapai 94 - 96% (Lohmann Tierzucht, 2019). Berdasarkan data maka disimpulkan perlu adanya analisis molekuler guna menentukan alel asosiatif terhadap produktivitas telur. Faktor manajemen dan sistem pemeliharaan tidak berkorelasi signifikan terhadap produktivitas telur sementara terdapat korelasi signifikan antara umur dewasa kelamin dan *inbreeding depression.*

Pengukuran parameter morfometrik telur ayam yaitu EW (*egg weight*), EI (*Eggshape Index*) dan beberapa parameter lain menentukan kualitas telur, grading, performa reproduktif dan perkembangan embrio ayam (Kabir *et al.,* 2012; Duman *et al.,* 2016). Dalam beberapa studi parameter kualitas telur berpengaruh signifikan dalam kegiatan komersil seperti distribusi telur, kegiatan penetasan *day old chicken* (DOC) dan ketertarikan konsumen (Abanikannda and Leigh, 2012; Ikegwu *et al.,* 2016). Parameter kualitas telur sendiri ditentukan oleh beberapa faktor seperti usia ayam petelur, genotip, nutrisi, sistem pemeliharaan dan waktu oviposisi (Duman *et al.,* 2016). Dalam Setiawati dkk. (2016) ditemukan bahwa terdapat korelasi signifikan antara sistem pemeliharaan (*cage* dan *litter cage*) dengan bobot telur (EW).

Pengukuran kualitas telur ayam $F_1$K dan $F_2$K dihitung berdasarkan beberapa parameter eksternal telur. Parameter kualitas telur eksternal yang digunakan dalam mengidentifikasi telur populasi $F_1$K dan $F_2$K terdiri atas:

1. EW: *Egg Weight* (gram)
2. ESA: *Egg Surface Area* ($cm^2$) ($3{,}9782xEW^{0.7056}$) (Duman *et al.,* 2016)
3. HV: *Hypothetical Volume* ($cm^3$) ($(0{,}6057-0{,}0018B)LB^2$) (Zhou *et al.,* 2009)
4. GMD: *Geometric Mean Diameter* (cm) ($Dg = (LW^2)^{1/3}$) (Ikegwu *et al.,* 2016)
5. Sp: *Sphericity* (cm) ((Dg/L)x100) (Ikegwu *et al.,* 2016)
6. EI: *Eggshape Index* (cm) ((W/L)x100 (Duman *et al.,* 2016)

Tabel 14. Deskripsi statistik produktivitas telur dan *Eggshape Index* (EI) $F_1$K dan $F_2$K

| Descriptive statistics for Eggshape Index (EI) of $F_1$K (n = 4) | | | | | |
|---|---|---|---|---|---|
| Shape Index | N (96) | Min (cm) | Max (cm) | Mean (cm) | Sd |
| Sharp egg | 51 | 60,45 | 71,99 | 70,19 | ±1,784 |
| Standard egg | 40 | 72,09 | 75,91 | 73,45 | ±1,007 |
| Round egg | 5 | 76,07 | 78,89 | 77,62 | ±1,329 |
| Descriptive statistics for Eggshape Index (EI) of $F_2$K (n = 4) | | | | | |
| Shape Index | N (66) | Min (cm) | Max (cm) | Mean (cm) | Sd |
| Sharp egg | 8 | 69,60 | 71,72 | 70,941 | ±0,785 |
| Standard egg | 15 | 72,06 | 75,74 | 73,96 | ±1,309 |
| Round egg | 43 | 76,20 | 81,87 | 78,654 | ±1,487 |

$F_1$K: $F_1$ Kambro; $F_2$K: $F_2$ Kambro (Olahan Pribadi, 2019)

Telur dapat diklasifikasi ke dalam 3 kategori berdasarkan *eggshape index* (EI) yaitu *sharp egg* (EI<72), *standard egg* (EI = 72-76) dan *round egg* (EI>76) (Duman *et al.,* 2016). Dalam Tabel 14 ditunjukkan rerata ukuran (EI, *Eggshape*

*Index*) pada populasi ayam $F_2K$ (n = 66) yaitu *sharp* (70,941 ± 0,278 cm), *standard* (73,96 ± 0,338 cm) dan *round* (78,654 ± 0,227 cm). Rerata EI populasi ayam $F_1K$ (n = 96) yaitu *sharp* (70,19 ± 0,249 cm), *standard* (73,45 ± 0,159 cm) dan *round* (77,62 ± 0,595 cm). Secara deskriptif ditemukan bahwa 53,125% telur ayam $F_1K$ (n = 96) tergolong *sharp egg*, sementara 65,15% telur $F_2K$ (n = 66) tergolong *round egg*.

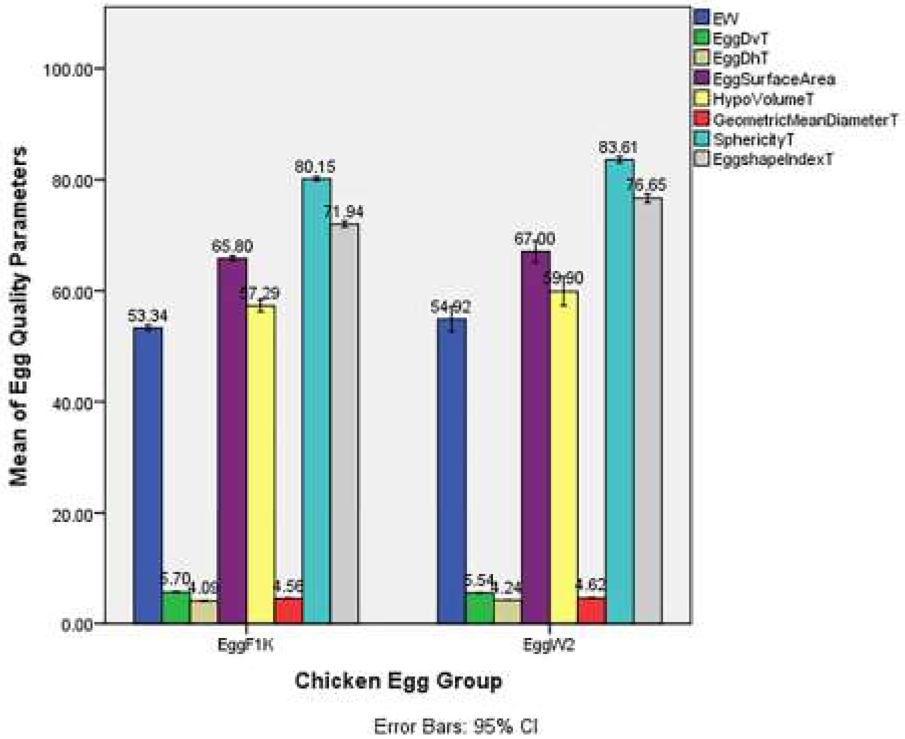

Gambar 14. Parameter kualitas telur Egg$F_1K$ dan EggW2 (Olahan Pribadi, 2019).
**Ket**: Egg$F_1K$: $F_1$ Kambro; EggW2: $F_2$ Kambro; EW: *Egg Weight*; EggDvT: lingkar vertikal; EggDhT: lingkar horizontal

Pembagian grade telur dibagi menjadi tiga yaitu AA (*perfect/standard egg*), A/B (*nearly perfect/sharp egg*) dan AB (*round egg*) (Duman *et al.,* 2016; Ikegwu *et al.,* 2016). Dalam Setiawati dkk. (2016) bentuk telur terdiri atas *biconical* (kedua ujungnya runcing), *elliptical* (elips), *oval* (bentuk terbaik) dan *spherical* (hampir bulat). Bentuk telur dipengaruhi oleh faktor genetik dan tidak ditemukan korelasi antara sistem pemeliharaan dan suhu terhadap bentuk telur (Setiawati dkk., 2016).

Pada Gambar 14 ditunjukkan perbandingan beberapa parameter kualitas telur eksternal ayam $F_1K$ dan $F_2K$. Berdasarkan parameter EW, EggDhT, *EggSurfaceArea*, *Sphericity*, *HypoVolume*, *GeometricMeanDiameter* dan

*Eggshape Index* maka ayam $F_2$K unggul dibandingkan ayam $F_1$K. Pada parameter EggDvT ayam $F_2$K memiliki rerata 5,54 cm lebih rendah dibandingkan ayam $F_1$K dengan rerata 5,7 cm. Berdasarkan hal ini maka bentuk telur pada ayam $F_2$K mendekati *round egg* dan $F_1$K mendekati *standard egg.* Dalam rangka menguji fenomena tersebut maka dilakukan analisis *independent sample t-test* pada telur ayam $F_1$K dan $F_2$K (Tabel 15).

Tabel 15. Efek EI terhadap parameter kualitas telur eksternal telur $F_1$K dan $F_2$K

| Quality Characteristics | Shape Index $F_1$K | | | | |
|---|---|---|---|---|---|
| | Sharp | Standard | Round | Sd | *p* |
| EW (gram) | 53,65a (± 1,958) | 53,06b (± 2,731) | 52,8a (± 2,588) | 2,34 | ns |
| ESA ($cm^2$) | 66,07a (± 1,703) | 65,52b (± 2,385) | 65,33a (± 2,25) | 2,04 | ns |
| HV ($cm^3$) | 57,723a (± 5,103) | 56,27a (± 4,475) | 60,976a (± 11,508) | 5,36 | ns |
| GMD (cm) | 4,575a (± 0,151) | 4,54a (± 0,124) | 4,652a (± 0,303) | 0,15 | ns |
| Sp ($cm^3$) | 78,86a (± 1,356) | 81,28a (± 0,742) | 84,32a (± 0,96) | 1,89 | *** |

| Eggshape Index of $F_1$K (T-Test) | | | | | | |
|---|---|---|---|---|---|---|
| Standard-Sharp | | Round- Standard | | Round- Sharp | | *p* |
| t | df | t | df | t | df | |
| 10,321*** | 89 | 8,421*** | 43 | 9,024*** | 54 | *** |

| Quality Characteristics | Shape Index $F_2$K | | | | |
|---|---|---|---|---|---|
| | Sharp | Standard | Round | Sd | *p* |
| EW (gram) | 59,25a (± 14,77) | 53,73a (± 8,713) | 54,535a (± 8,163) | 9,254 | ns |
| ESA ($cm^2$) | 70,486a (± 12,33) | 66a (± 7,308) | 66,705a (± 6,896) | 7,771 | ns |
| HV ($cm^3$) | 67,74a (± 12,59) | 63,65a (± 13,84) | 57,13b (± 7,013) | 10,29 | * |
| GMD (cm) | 4,82a (± 0,294) | 4,71a (± 0,331) | 4,56b (± 0,182) | 0,252 | * |
| Sp ($cm^3$) | 79,42a (± 0,586) | 81,65a (± 0,965) | 85,08a (± 1,073) | 2,34 | ns |

| Eggshape Index of $F_2$K (T-Test) | | | | | | |
|---|---|---|---|---|---|---|
| Standard-Sharp | | Round- Standard | | Round- Sharp | | *p* |
| t | df | t | df | t | df | |
| 5,931*** | 21 | 10,846*** | 56 | 14,224*** | 49 | *** |

EW: *Egg Weight*; ESA: *Egg Surface Area*; HV: *Hypothetical Volume*; GMD: *Geometric Mean Diameter*; Sp: *Sphericity*. ns: *non-significant*; *: p<0,05; **: p<0,01; ***: p<0,001. Sd: Standar Deviasi terletak dibawah Mean; SEM: *Standard Error of Mean*. Mean dengan huruf berbeda dalam satu baris berbeda secara signifkan berdasarkan Fisher's LSD and Tukey HSD (p<0,05) (Olahan Pribadi, 2019).

Analisis *independent sample t-test* dilakukan pada EI ayam $F_1$K dan ayam $F_2$K (Tabel 15). Pada grup ayam $F_1$K *round egg* unggul secara signifikan terhadap *standard egg* (t (43) = 8,421, p<0,001). Pada grup ayam $F_2$K *round egg* unggul secara signifikan terhadap *standard egg* (t (56) = 10,846, p<0,001). Berdasarkan hasil uji *t-test* pada bentuk telur $F_1$K dan $F_2$K maka disimpulkan bahwa bentuk telur grup ayam $F_1K$ dan $F_2K$ didominasi oleh *round egg*. Generasi parental ayam hibrida $F_1K$ yaitu Broiler Cobb 500 memiliki nilai EI 69,79 ± 1,95 cm (*sharp egg*) dan Pelung Blirik Hitam memiliki nilai EI 78,10 ± 7,11 cm (*round egg*) (Kabir *et al.,* 2012). Nilai EI ayam Layer ISA Brown dan ISA White yaitu 79,90 ± 2,83 cm (*round* egg) dan 72,08 ± 2,7 cm (*standard egg*) (Kabir *et al.,* 2012). Berdasarkan hasil perbandingan maka dapat diasumsikan bahwa *sharp egg* dan *round egg* pada ayam $F_1K$ dan $F_2K$ dipengaruhi oleh pewarisan alel tetuanya dan dapat menunjukkan adanya fenomena segregasi alel pada populasi hibrida tersebut.

Nilai EI sendiri memiliki pengaruh yang signifikan pada beberapa parameter kualitas telur eksternal. Pada ayam $F_1K$ EI berpengaruh terhadap Sp telur ($p<0,001$) dan pada ayam $F_2K$ berpengaruh terhadap HV ($p<0,05$) dan GMD ($p<0,05$) (Tabel 15). Parameter ESA, HV, GMD dan Sp merupakan parameter yang digunakan dalam beberapa studi untuk memprediksi bobot tetas DOC, daya tetas telur (*hatchability*), kualitas cangkang telur, parameter kualitas interior telur, fungsi reproduksi, klasifikasi dan seleksi persilangan (Narushin, 2005; Havlíček *et al.,* 2008; Duman *et al.,* 2016; Ikegwu *et al.,* 2016). Dalam studi terpisah yang dilakukan oleh Havlíček *et al.* (2008) dan Waranusast *et al.* (2017) digunakan metode *image-based morphometrical measurement* telur ayam dengan tujuan meningkatkan efisiensi dalam menseleksi telur ayam berdasarkan kualitas eksternal, kategorisasi dan grading. Metode ini dianggap sukses menentukan dimensi telur dengan tingkat error sebesar 3,1% dan akurasi hingga 80,4% dalam mensortir 425 telur menggunakan data foto telur (Waranusast *et al.,* 2017).

Berdasarkan parameter Sp ($p<0,001$), HV (*round egg*; 57,13 ± 7,013, $p<0,05$) dan GMD (*round egg*; 4,56 ± 0,182, $p<0,05$) maka dapat disimpulkan bahwa bentuk telur $F_1K$ dan $F_2K$ tergolong *round egg* (EI>76), sehingga mengacu pada Ikegwu *et al.* (2016) maka kualitasnya tergolong dalam kategori AB. Berdasarkan parameter EW, ESA dan EI maka tidak terdapat pengaruh (ns; $p>0,05$) signifikan antara bentuk telur terhadap bobot telur dan luas permukaan cangkang telur. Dapat disimpulkan bahwa telur ayam $F_1K$ memiliki bobot dan luas permukaan seragam pada ketiga kategori bentuk telur, hal yang sama turut berlaku pada telur ayam $F_2K$.

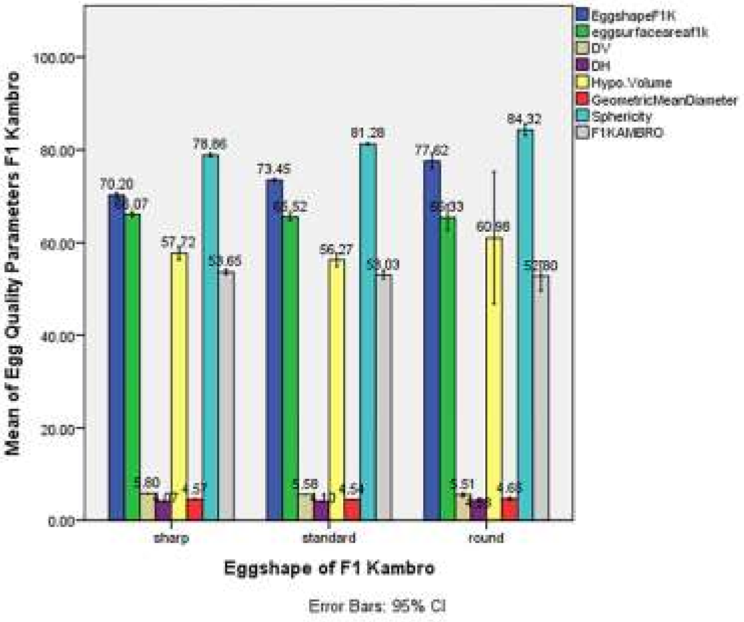

**(a)**

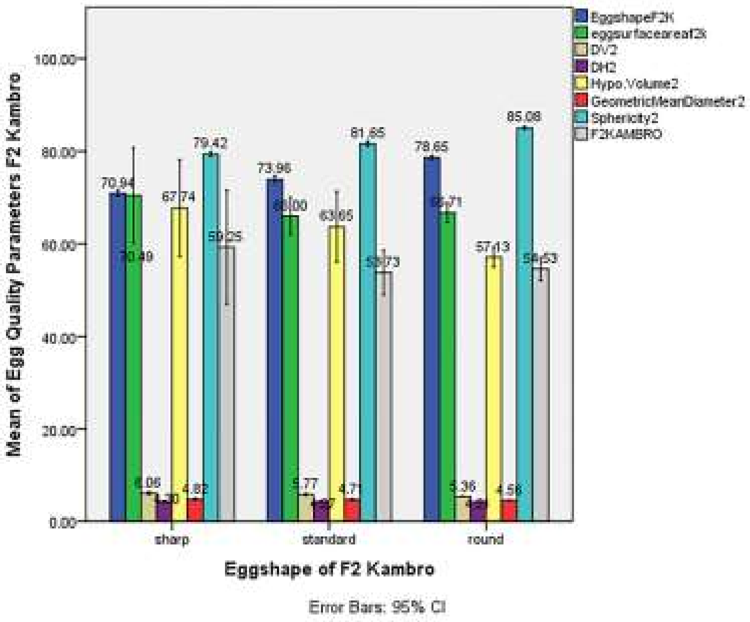

**(b)**

Gambar 15. Parameter kualitas telur eksternal (Olahan Pribadi, 2019).
(a) Parameter kualitas telur $F_1K$; (b) Parameter kualitas telur $F_2K$

Pada Gambar 15 ditunjukkan bahwa parameter kualitas telur eksterior pada masing-masing kategori telur. Berdasarkan Dv (lingkar vertikal) dan Dh (lingkar horizontal) ayam $F_1K$ dan $F_2K$ ditunjukkan bahwa telur kategori *sharp egg* memiliki rasio ukuran tertinggi terhadap *standard egg* dan *round egg*.

Berdasarkan parameter Sp maka telur $F_2K$ cederung lebih *spherical* menurut Setiawati dkk. (2016) dengan nilai HV pada $F_2$ lebih tinggi dibandingkan $F_1K$ pada kategori telur *sharp egg* dan *standard egg*. Dalam Duman *et al* (2016) ditemukan bahwa tidak terdapat pengaruh signifikan antara EI dan EW pada tiap kategori telur.

Berdasarkan rerata EW ayam $F_1K$ (53,34 ± 2,34 gram) dan $F_2K$ (54,92 ± 9,25 gram) maka tergolong ke dalam kategori *small egg* (Waranusast *et al.,* 2017). Dalam Belitz (2009) dikemukakan bahwa rerata berat telur ayam adalah 58 gram dengan tiga komponen yaitu kuning telur, putih telur dan cangkang. Parameter EW ayam Broiler Cobb 500 dapat mencapai 56,20 ± 1,62 gram dan Pelung dapat mencapai 20,20 ± 4,76 gram (Kabir *et al.,* 2012). Dari segi bobot tetas ayam $F_2K$ mencapai 36 ± 3,74 gram dan bobot tetas ayam $F_1K$ 35,18 ± 3,45 gram unggul terhadap ayam lokal asli Pelung yaitu 35,12 gram (Depison, 2009). Dalam Kabir *et al.* (2012) dijelaskan bahwa bobot telur berhubungan dengan parameter kualitas interior telur yaitu tinggi albumin dan kuning telur. Dalam Sekeroğlu *et al.* (2000) dijelaskan bahwa EI dapat digunakan sebagai penentu panjang albumin, lebar kuning telur, tinggi kuning telur dan warna kuning telur. Berdasarkan hal tersebut maka korelasi antara parameter kualitas eksterior telur dan EI dapat diselidiki (Tabel 16).

Tabel 16. Korelasi EI terhadap parameter kualitas telur eksternal ayam $F_1K$ dan $F_2K$

| Quality Characteristics | Sharp | Standard | Round | Eggshape Index |
|---|---|---|---|---|
| EW (gram) | 0,105 | -0,274 | 0,545 | -0,116 ns |
| ESA ($cm^2$) | 0,106 | -0,277 | 0,545 | -0,117 ns |
| HV ($cm^3$) | 0,433** | -0,167 | 0,633 | 0,164 ns |
| GMD (cm) | 0,477** | -0,158 | 0,661 | 0,180 ns |
| Sp ($cm^3$) | 1** | 1** | 1** | 1 ** |
| Quality Characteristics | Sharp | Standard | Round | Eggshape Index |
| EW (gram) | 0,012 | 0,413 | 0,025 | -0,054 ns |
| ESA ($cm^2$) | 0,004 | 0,418 | 0,022 | -0,049 ns |
| HV ($cm^3$) | -0,050 | -0,243 | 0,030 | -0,374 ** |
| GMD (cm) | -0,061 | -0,226 | 0,034 | -0,363 ** |
| Sp ($cm^3$) | 1** | 1** | 1** | 1** |

EW: *Egg Weight*; ESA: *Egg Surface Area*; HV: *Hypothetical Volume*; GMD: *Geometric Mean Diameter*; Sp: *Sphericity*. ns: *non-significant*; *: p<0,05; **: p<0,01; ***: p<0,001 (Olahan Pribadi, 2019)

Berdasarkan Tabel 16 ayam $F_1K$ memiliki korelasi signifikan (p<0,001) terhadap Sp dan ayam $F_2K$ memiliki korelasi negatif signifikan (p<0,001) terhadap HV, GMD dan dan korelasi positif signifikan ($p<0,001$) terhadap Sp. Pada grup ayam $F_1K$ terdapat korelasi positif signifikan antara *sharp egg* terhadap HV (0,433, $p<0,01$) dan GMD (0,477, $p<0,01$). Parameter EW dan ESA tidak berkorelasi terhadap EI (ns; $p>0,05$) pada ayam $F_1K$ dan $F_2K$. Berdasarkan data tersebut maka perlu dilakukan uji parameter kualitas interior dalam menentukan kualitas telur ayam sebab tidak terdapat korelasi signifikan antara EW terhadap EI. Dalam Duman *et al.* (2016) disimpulkan bahwa EI berkorelasi signifikan terhadap EW namun tidak berkorelasi terhadap indeks kuning telur.

Berdasarkan nilai korelasi (rGMD = -0,363, $p<0,001$) maka parameter GMD dapat digunakan sebagai variabel prediktor dalam menentukan EW pada ayam Kambro. Parameter Sp memiliki korelasi, r = 1, $p<0,001$ sehingga tidak dapat menggambarkan efek pergeseran kurva regresi. Parameter HV memiliki nilai korelasi lebih kecil terhadap parameter GMD (rHv = -0,374, $p<0,001$).

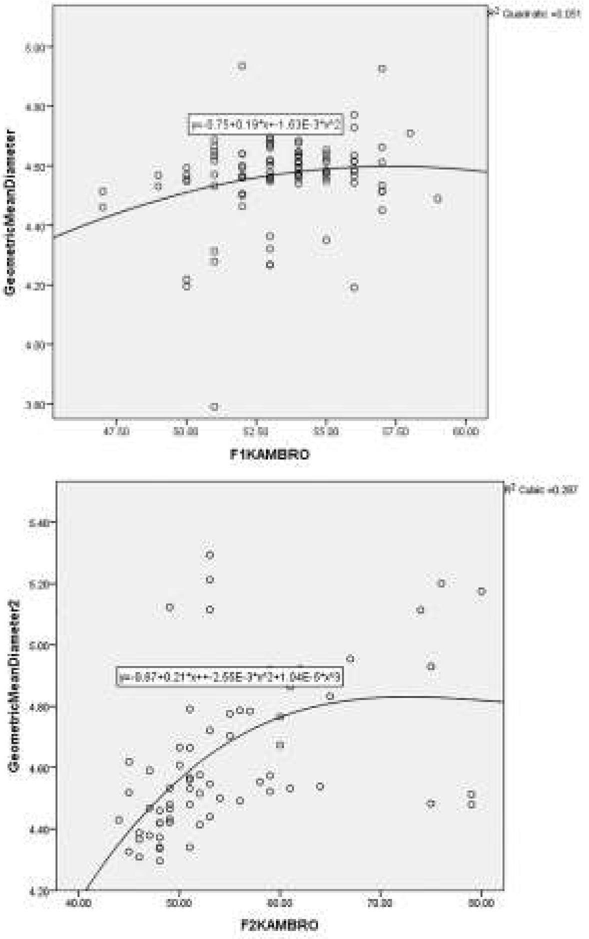

Gambar 16. Regresi Quadratic and Cubic Parameter GMD terhadap EW $F_1K$ dan $F_2K$ (Olahan Pribadi, 2019). GMD: *Geometric Mean Diameter*; $F_1K$: $F_1$ Kambro; $F_2K$: $F_2$ Kambro

Dalam Gambar 16 ditunjukkan hasil regresi linear parameter GMD terhadap EW $F_1$K dan $F_2$K. Berdasarkan hasil regresi linear GMD terhadap EW $F_1$K maka metode regresi dengan r tertinggi yaitu *quadratic* dengan capaian r = 0,051 dan persamaan **y = -0,75 + 0,19*x ± 1,63E-3*x^2**. Berdasarkan hasil regresi linear GMD terhadap $F_2$K maka metode regresi dengan r tertinggi yaitu *cubic* dengan capaian r = 0,287 dan persamaan **y = -0,67 +0,21*x+ ± 2,55E-3*x^2+1,04E-5*x^3**.

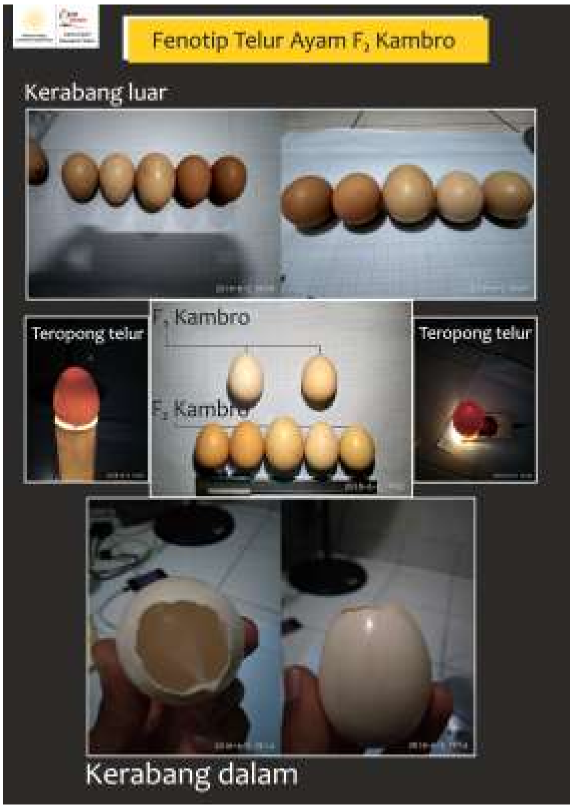

Gambar 17. Fenotip telur ayam $F_1$ Kambro dan $F_2$ Kambro (Dok. Pribadi, 2018)

Observasi skoring visual terhadap 96 butir telur ayam $F_1$ Kambro dan 66 butir telur ayam $F_2$ Kambro menunjukkan bahwa 100% telur ayam $F_1$ Kambro berwarna *white* (Gambar 17). Pada ayam $F_2$ Kambro ditemukan adanya empat tipe warna yaitu *light brown*, *cream, brown* dan *white*. Berdasarkan hasil observasi maka dapat disimpulkan bahwa terjadi segregasi dalam pewarisan alel sehingga bepengaruh terhadap warna cangkang telur. Dalam Lukanov *et al.* (2015) ditemukan bahwa yang mempengaruhi warna cangkang telur ayam domestik dan *wild-type* adalah protoporphyrin IX, biliverdin IX dan biliverdin zinc chelate. Dalam Setiawati dkk. (2016) dijelaskan bahwa ayam petelur medium memiliki kerabang atau cangkang berwarna cokelat dan tipe ringan berwarna putih. Dalam hal ini maka ayam Kambro tergolong ke dalam ayam tipe petelur ringan. Dalam kegiatan seleksi selanjutnya dapat dipilih generasi parental $F_2$ Kambro dengan tipe warna cokelat untuk dapat meningkatkan produksi filial $F_3$ Kambro.

#### C. Koefisien *Inbreeding* Ayam $F_2$ Kambro

*Inbreeding* dapat mengakibatkan penurunan kesehatan hibrida dan kemampuan reproduktif sehingga dapat digolongkan ke dalam efek negatif dalam persilangan. *Inbreeding* dapat menyebabkan penurunan persentase sperma normal dan peningkatan abnormalitas sperma (Oldenbroek and van der Waaij, 2014). Perkawinan antara parental sekerabat dapat meningkatkan tingkat homozigositas alel pada filial akibat *non random mating* (Eldik *et al.*, 2006). Peningkatan homozigositas alel dapat mengakibatkan penurunan variasi genetik populasi hibrida. Variasi genetik merupakan tingkat perbedaan genetik antara individu satu spesies, antar generasi atau di dalam suatu generasi tertentu (Oldenbroek and van der Waaij, 2014).

Tingkat *inbreeding* ditentukan dengan *inbreeding coefficient* populasi ayam $F_2$ Kambro. *Inbreeding coefficient* menentukan probabilitas hibrida ayam $F_2$ Kambro mewarisi alel generasi *grandparent stock* dan *parent stock* (Gambar 18). *Inbreeding coefficient* memiliki nilai antara 0 (*not inbred*) hingga 1 (*fully inbred*). Peningkatan nilai *inbreeding coefficient* (Fx) menentukan tingkat heterozigositas alel dalam populasi dan dapat mengakibatkan fenomena *inbreeding depression* (Konig *et al.,* 2010; Shad *et al.,* 2013; Wakchaure and Ganguly, 2015; Nietlisbach *et al.,* 2017).

Formula dalam menghitung *inbreeding coefficient* sebagai berikut:

$$\mathbf{F = \Sigma[(\tfrac{1}{2}^{n+1})\ (1 + F_{CA})]} \qquad (1)$$

F = nilai koefisien inbreeding
n = banyaknya garis dalam alur
FCA = koefisien moyang bersama

Telalbasic *et al.* (2007)

Formula dalam menghitung laju *inbreeding* sebagai berikut:

$$\mathbf{Laju\ inbreeding = 1/8\ Nm + 1/8\ Nf} \qquad (2)$$

Nm = jumlah pejantan dan calon pejantan
Nf = jumlah betina yang dapat dikawinkan

Sawitri dan Takandjandji (2012)

Dalam persilangan ayam $F_2$ Kambro terdapat distribusi alel dari *grandparent stock* yaitu jantan Pelung Blirik Hitam dan Broiler Cobb 500. Generasi *parent stock* yaitu ayam jantan dan betina $F_1$ Kambro yang disilangkan untuk menghasilkan ayam $F_2$ Kambro sesuai diagram silsilah pada Gambar 18.

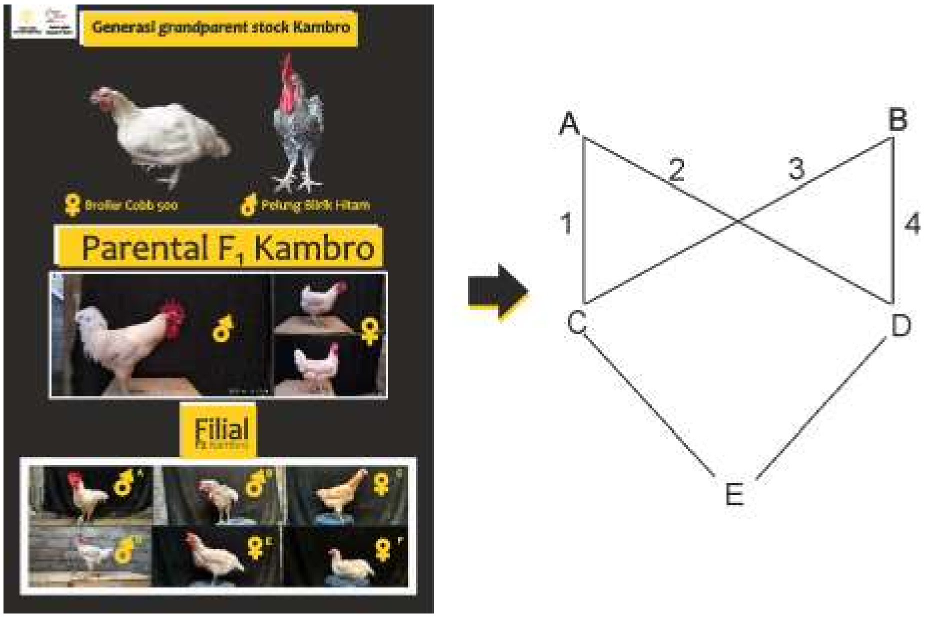

Gambar 18. Rekonstruksi diagram silsilah ayam $F_2$ Kambro
(Olahan Pribadi, 2019).
A: Broiler Cobb 500; B: Pelung Blirik Hitam; C: Jantan $F_1$ Kambro;
D: Betina $F_1$ Kambro; E: generasi $F_2$ Kambro

Berdasarkan diagram *inbreeding* ayam $F_2$ Kambro (Gambar 18) maka dapat dilakukan perhitungan *inbreeding* dan laju *inbreeding*. Berdasarkan perhitungan maka *inbreeding coefficient* (Fx) $F_2$ Kambro yaitu 25 % dan laju *inbreeding* (LI) yaitu 4,925%. Dalam populasi ayam $F_2$ Kambro nilai Fx menunjukkan bahwa lokus heterozigot pada generasi *grandparent stock* (GS) dan *parent stock* (PS) menjadi semakin homozigot. Dalam Habibah (2018) nilai ambang toleransi Fx sebesar 37,5%, dalam hal ini Fx $F_2$ Kambro (Fx<37,5%) masih dapat ditoleransi.

Nilai Fx dapat mengalami peningkatan apabila individu dalam generasi $F_2$ Kambro disilangkan dengan kerabatnya berdasarkan nilai LI sebesar 4,925%. Dalam Sawitri dan Takandjandji (2012) ambang toleransi LI sebesar 2% sehingga resiko terjadinya *inbreeding depression* pada perkawinan $F_2$ Kambro sekerabat dapat menghasilkan $F_3$ Kambro depresiasi karakter fenotip. Dalam Nietlisbach *et* *al.* (2017) *inbreeding depression* merupakan depresiasi nilai vitalitas (*fitness*) yang disebabkan oleh peningkatan probabilitas *identical-by-descent* (IBD). IBD merupakan probabilitas dua lokus alel homolog diwariskan dari generasi indukan kepada individu generasi selanjutnya (*inbred*) (Nietlisbach *et al.,* 2017). Peningkatan homozigositas berasosiasi dengan penurunan nilai vitalitas (*fitness*) disebabkan peningkatan ekspresi sebagian atau seluruh alel resesif atau alel homozigot yang diwariskan mengekspresikan karakter fenotip inferior dibandingkan alel heterozigot (Halverson *et al.,* 2006; Nietlisbach *et al.,* 2017). Dalam Wakchaure and Ganguly (2015) diketahui bahwa *inbreeding* berpengaruh terhadap beberapa performa fenotip diantaranya penurunan produktivitas reproduktif, peningkatan mortalitas, penurunan laju pertumbuhan dan penurunan imunitas hibrida.

Dalam populasi ayam $F_2$ Kambro terdapat tingkat mortalitas yang tinggi sebesar 50% dari total populasi (n = 22) lebih tinggi dibandingkan populasi ayam $F_1$ Kambro sebesar 5,5% dari total populasi (n = 20). Hal ini mengindikasikan adanya efek negatif persilangan sekerabat ayam $F_1$ Kambro disamping adanya beberapa faktor lain seperti infeksi coryza atau snot dan manajemen pemeliharaan.

Tabel 17. Rerata bobot tubuh (BW), parameter kualitas telur dan produktivitas telur ayam $F_2$ Kambro dan $F_1$ Kambro

| | Chicken Group | | | | | |
|---|---|---|---|---|---|---|
| | $F_2$K (n= 11) | BC5 (n = 22) | $F_1$K (n =17) | $F_1$P (n= 7) | *F* | η2 |
| BW (gram) | 753,36a (± 155,31) | 1.706,82b (± 262,54) | 1.244,14ab (± 453,82) | 602,88a (± 79,93) | $68{,}896^{***}$ | 0,796 |

| Descriptive statistics for eggshape index (EI) of $F_1$K | | | | | | |
|---|---|---|---|---|---|---|
| Shape Index | N (96) | Min (cm) | Max (cm) | Mean (cm) | SEM | Sd |
| Sharp egg | 51 | 60,45 | 71,99 | 70,19 | 0,249 | ±1,784 |
| Standard egg | 40 | 72,09 | 75,91 | 73,45 | 0,159 | ±1,007 |
| Round egg | 5 | 76,07 | 78,89 | 77,62 | 0,595 | ±1,329 |
| Descriptive statistics for eggshape index (EI) of $F_2$K | | | | | | |
| Shape Index | N (66) | Min | Max | Mean | SEM | Sd |
| Sharp egg | 8 | 69,60 | 71,72 | 70,941 | 0,278 | ±0,785 |
| Standard egg | 15 | 72,06 | 75,74 | 73,96 | 0,338 | ±1,309 |
| Round egg | 43 | 76,20 | 81,87 | 78,654 | 0,227 | ±1,487 |

| Quality Characteristics | Eggshape Index (EI) | | | | | |
|---|---|---|---|---|---|---|
| | Sharp | Standard | Round | SEM | Sd | *p* |
| EW $F_1$K (gram) | 53,65a (± 1,958) | 53,06b (± 2,731) | 52,8a (± 2,588) | 0,239 | 2,34 | ns |
| EW $F_2$K (gram) | 59,25a (± 14,77) | 53,73a (± 8,713) | 54,535a (± 8,163) | 1,12 | 9,254 | ns |

EW: *Egg Weight*; $F_1$K: $F_1$ Kambro; $F_2$K: $F_2$ Kambro (Olahan Pribadi, 2019)

Penurunan signifikan ($p<0{,}001$) capaian bobot tubuh (BW) ayam $F_2$ Kambro (753,36 ± 155,31) terhadap ayam $F_1$ Kambro (1.244,14 ± 453,82) (Tabel 14). Periode pengukuran produktivitas telur populasi ayam $F_1$ Kambro dan $F_2$ Kambro berlangsung selama ± 270 hari (April hingga Desember 2018). Produktivitas telur pada ayam $F_1$ Kambro sebanyak 96 butir mengungguli $F_2$ Kambro sebanyak 66 butir. Betina $F_1$ Kambro (n = 4) memasuki masa siap kelamin pada umur 6 bulan sementara betina $F_2$ Kambro (n = 4) pada umur 4 bulan. Dari segi produktivitas telur maka dapat disimpulkan bahwa ayam $F_2$ Kambro mengalami *inbreeding depression.* Dari segi bobot dan ukuran telur, telur ayam $F_2$ Kambro baik kategori *sharp, standard* dan *round* mengungguli telur ayam $F_1$ Kambro. Rerata ukuran (EI, *Eggshape Index*) pada populasi ayam $F_2$ Kambro yaitu *sharp* (70,941 ± 0,278 cm), *standard* (73,96 ± 0,338 cm) dan *round* (78,654 ± 0,227 cm). Rerata EI populasi ayam $F_1$ Kambro yaitu *sharp* (70,19 ± 0,249 cm), *standard* (73,45 ± 0,159 cm) dan *round* (77,62 ± 0,595 cm). Dari segi rerata bobot telur (EW, *Egg Weight*) ayam $F_1$ Kambro pada kategori *sharp* (53,65 ± 1,958 gram), *standard* (53,06 ± 2,731 gram) dan *round* (52,8 ± 2,588 gram), sedangkan ayam $F_2$ Kambro pada kategori *sharp* (59,25 ± 14,77 gram), *standard* (53,73 ± 8,713 gram) dan *round* (54,535 ± 8,163 gram). Berdasarkan ukuran telur dan bobot telur antara $F_1$K dan $F_2$K maka dapat disimpukan bahwa *inbreeding depression* bersifat searah dan random. Hal ini dapat disebabkan oleh pewarisan beberapa alel homozigot inferior terhadap pertumbuhan bobot tubuh dan produktivitas telur namun bersifat superior dalam penentuan kualitas telur ayam $F_2$ Kambro.

Beberapa penelitian menunjukkan terdapat korelasi signifikan antara peningkatan Fx terhadap penuruan capaian bobot tubuh dan produktivitas telur ayam (Sewalem *et al.,* 1999; Szwaczkowski *et al.,* 2003; Nwagu *et al.,* 2007; Yerturk *et al.,* 2008). Dalam Shad *et al.* (2013) dilaporkan bahwa tidak terdapat korelasi antara peningkatan Fx (0.002%) terhadap penurunan bobot tubuh dan produktivitas telur dengan faktor utama yaitu rendahnya individu *inbred* dalam populasi ayam lokal asli Iran. Dalam Konig *et al.* (2010) dilaporkan bahwa laju *inbreeding* dalam populasi *White Leghorn* sebesar 0,95% yang mengancam struktur produksi ayam tersebut sehingga membutuhkan program seleksi yang spesifik.

Persilangan *inbreeding* memiliki dampak negatif namun vital dalam pengembangan suatu galur atau strain khususnya pada ayam. Dalam pengembangan suatu galur ayam *inbreeding* berperan dalam mengeliminasi abnormalitas, alel letal dan beberapa karakter yang secara komersil inferior (Wakchaure and Ganguly, 2015). Perubahan frekuensi genetik dalam populasi hibrida ayam dapat difokuskan terhadap beberapa alel superior dengan meningkatkan relasi genetik hibrida.

Beberapa langkah yang dapat dilakukan dalam mengatasi efeek negatif persilangan *inbreeding* diantaranya adalah sebagai berikut:

1. Persilangan antar individu segalur non sekerabat
2. Peningkatan jumlah pejantan dan penggunaan jantan baru
3. Ukuran populasi efektif untuk menekan laju *inbreeding*
4. Konservasi plasma nutfah hewan dan *crossbreeding*
5. Struktur *pedigree* persilangan yang detail dan akurat

Wakchaure and Ganguly (2015)

Penggunaan *genomic selection* menggunakan marker gen dan mikrosatelit dapat menekan peningkatan Fx dan LI (Nietlisbach *et al*., 2017). Dalam Wolc *et al.* (2015) dilaporkan bahwa nilai Fx dan LI pada metode *genomic selection* 16 galur ayam Layer lebih rendah dibandingkan metode seleksi konvensional.

#### D. Karakter Genotip Ayam $F_2$ Kambro dan Validasi T-ARMS PCR

Berdasarkan data parameter bobot tubuh, parameter kualitas telur eksterior, produktivitas telur, fenotip telur dan koefisien *inbreeding* maka diperlukan investigasi molekuler. Investigasi molekuler gen atau alel terkait dengan performa reproduksi dan pertambahan bobot ayam khususnya tipe pedaging seperti ayam Kambro dapat dilakukan dengan *genotyping* gen pengkode leptin. Leptin dikode oleh gen *LEP* dan lokalisasi genomiknya pada *Gallus gallus gallus* masih menjadi perdebatan selama 20 tahun. Reseptor leptin disebut *leptin receptor* dikode oleh gen *LEPR* dan telah terpetakan pada ayam. Dalam penelitian ini akan digunakan metode *genotyping* T-ARMS PCR dalam mendeteksi *single nucloetide polymorphisms* (SNPs) pada ekson 9 *LEPR* ayam.

Leptin diekspresikan oleh gen *ob* dan mutasinya mengakibatkan abnormalitas fenotip seperti obesitas dan infertilitas (Zhang *et al*., 1994; Klok *et al*., 2007; Ohkubo and Adachi, 2008; Ohkubo, 2014). Leptin berasal dari kata yunani, leptos yang berarti tipis atau kurus (Hossner, 1998; Mrazova *et al*., 2012). Pada ayam gen leptin diidentifikasi sebagai *LEP* atau *leptin gene* (Ninov *et al*., 2008; Seroussi *et al*., 2016; Seroussi *et al*., 2017). Riset *LEP* sifatnya vital bagi peternakan ayam dalam kaitannya dengan peningkatan produktivitas (Ohkubo and Adachi, 2008; Ohkubo, 2014). Perdebatan mengenai *LEP* autentik ayam (*Gallus gallus gallus*) telah terjadi selama 20 tahun (Rodriguez, 2014; Seroussi *et al*., 2016). Ekspresi cDNA *LEP* unggas identik dengan cDNA *LEP* pada tikus (Taouis *et al*., 1998; Ashwell, 1999, Dai *et al*., 2007; Ohkubo, 2014). Beberapa tim riset meragukan lokalisasi *LEP* pada genom ayam (Friedman-Einat *et al*., 1999; Pitel *et al*., 2000; Dunn *et al*., 2001; Amills *et al*., 2003). Pitel *et al*. (2000) menyimpulkan bahwa *LEP* pada ayam belum terpetakan dengan rinci. Ekspresi leptin ayam pada sel Purkinje cerebellum relevansinya tinggi dengan administrasi eksternal leptin pada sel Purkinje cerebellum tikus dengan pengaruhnya terhadap propagasi, aktivasi dan viabilitas sel (Seroussi *et al*., 2016). Seroussi *et al*. (2016) mendeteksi pola ekspresi *LEP* dan *LEPR* pada beberapa jaringan ayam diantaranya embrio, ovarium, hipotalamus, cerebrum dan kelenjar adrenal, namun tidak terdeteksi pada jaringan hati, otot dada dan paru-paru. Lebih jauh pada Seroussi *et al*. (2017) berdasarkan perbandingan pemetaan genomik dan karakteristik sekuen, leptin ayam tidak terlokalisasi pada mikrokromosom (Seroussi *et al*., 2016) dengan konten G-C yang tinggi dan region repetitif, melainkan pada ujung distal 1p kromosom ayam (*Gallus gallus gallus*).

Gen *LEP* meregulasi asupan kalori, alokasi energi dan fungsi responsif endokrin hipotalamus dalam transformasi nutrisi (Abbasi *et al*., 2011). Peningkatan efisiensi pakan termasuk faktor kunci dalam mengurangi biaya produksi peternakan ayam dan meminimalisir dampak lingkungan peternakan ayam (de Verdal *et al*., 2011). Pakan mempengaruhi 70% biaya produksi peternakan ayam pedaging (Zhang and Aggrey, 2003). Beberapa faktor yang dapat mengurangi asupan pakan yaitu perbaikan genetik dengan persilangan untuk menekan laju kebutuhan *feed conversion ratio* (FCR) (Ferket and Gernat, 2006). Riset mendetail mengenai ekspresi, lokalisasi dan asosiasi fisiologis leptin pada ayam dapat meningkatkan produktivitas peternakan ayam.

Dalam penelitian ini mutasi bersifat transversi basa C ke A pada urutan basa nukleotida ke-127 (C127A) dengan produk amplikon sepanjang 174 bp. Metode T-ARMS PCR menggunakan 4 primer terdiri atas sepasang *Forward Outer Primer* (FOP) dan *Reverse Outer Primer* (ROP) dengan produk amplikon sepanjang 228 bp, *Forward Inner A Alelle* (FIA) dengan produk amplikon 114 bp dan *Reverse Inner C Allele* (RIC) dengan produk 169 bp. Konfigurasi optimasi temperatur, rasio primer dan konsentrasi primer dapat dilihat pada Tabel 18.

Tabel 18. Optimasi prosedur T-ARMS PCR deteksi SNPs ekson 9 *LEPR*

| Exp. | DNA (ng/ µL) | PC (pmol/µM) | PR (µL) | | | | PCR Steps | | | | | | Ag (%) | Electrophoresis | |
|---|---|---|---|---|---|---|---|---|---|---|---|---|---|---|---|
| | | IP/OP | FIA | RIC | FOP | ROP | ID (°C/mins) | Cyl (X) | DS (°C/s) | A (°C/s) | E (°C/s) | FE (°C/mins) | | V (volt) | D (mins) |
| 1 | 50 | 10/10 | 0,5 | 0,5 | 0,5 | 0,5 | 94/5 | 40 | 95/30 | Gradient/30 | 68/60 | 68/5 | 2 | 50 | 55 |
| 2 | 50 | 10/10 | 0,5 | 0,5 | 0,5 | 0,5 | 94/5 | 40 | 95/30 | 56,2/30 | 68/60 | 68/5 | 2 | 50 | 55 |
| 3a | 50 | 10/10 | 0,75 | 0,75 | 0,25 | 0,25 | 95/5 | 36 | 95/60 | 56,2/60 | 72/60 | 72/10 | 2 | 50 | 55 |
| 3b | 50 | 10/10 | 0,75 | 0,75 | 0,25 | 0,25 | 94/5 | 40 | 95/30 | 56,2/30 | 68/60 | 68/5 | 2 | 50 | 55 |
| 4 | 50 | 10/10 | 0,9 | 0,9 | 0,1 | 0,1 | 95/2 | 30 | 95/60 | 56,2/60 | 72/60 | 72/10 | 2 | 50 | 55 |
| 5 | 50 | 10/10 | 1,5 | 1,5 | 1 | 1 | 95/2 | 30 | 95/60 | 55,7/60 | 72/60 | 72/10 | 2 | 50 | 55 |
| 6a | 50 | 10/10 | 1,25 | 1,25 | 0,25 | 0,25 | 95/2 | 30 | 95/60 | 55,7/60 | 72/60 | 72/10 | 2 | 50 | 55 |
| 6b | 50 | 10/10 | 0,75 | 0,75 | 0,25 | 0,25 | 94/5 | 40 | 95/30 | 56,2/30 | 68/60 | 68/5 | 2 | 50 | 55 |
| 7 | 50 | 10/10 | 0,75 | 0,75 | 0,25 | 0,25 | 94/5 | 40 | 95/30 | 56,2/30 | 68/60 | 68/5 | 2 | 50 | 55 |
| 8 | 50 | 10/10 | 0,75 | 0,75 | 0,25 | 0,25 | 95/5 | 36 | 95/60 | 56,2/60 | 72/60 | 72/10 | 2 | 50 | 55 |
| 9 | 100 | 10/1 | 0,5 | 0,5 | 0,5 | 0,5 | 94/ 5 | 30 | 94/30 | 55,7/30 | 72/40 | 72/10 | 2.5 | 50 | 40 |

**FIA**: *Forward Inner A Allele*; **RIC**: *Reverse Inner C Allele*; **FOP**: *Forward Outer Primer*; **ROP**: *Reverse Outer Primer*; **PC**: *Primers Concentration*; **PR**: *Primers Ratio*; **IP/OP**: *Inner Primers/Outer Primers*; **ID**: *Initial Denaturation*; **Cyl**: *Cycle*; **DS**: *Denaturation Steps*; **A**: *Annealing*; **E**: *Extension*; **FE**: *Final Extension*; **Ag**: *Agarose*; **V**: *Voltage*; **D**: *Duration*; **Gradient**: 53˚C; 53,3˚C; 53,8˚C; 54,6˚C; 55,5˚C; 56,2˚C; 56,7˚C dan 57˚C (Dok. Pribadi, 2019)

*Tetra Primer Amplification Refractory Mutation System PCR* (Tetra ARMS PCR /T-ARMS PCR) tergolong dalam metode *genotyping*. Metode ini dikembangkan berdasarkan kekurangan yang terdapat pada amplifikasi PCR konvensional yang tidak efisien atau sepenuhnya refraktori apabila terdapat *mismatch* antara terminal 3' primer nukleotida dan sekuen DNA template (Newton *et al.,* 1989; Ye *et al.,* 1992; Landsverk and Wong, 2013; Alyethodi *et al.,* 2016; Peng *et al.,* 2017). Metode ini menggunakan 4 set primer, dua *outer* primer (OF, OR) dan dua inner primer pendeteksi alel spesifik (IF, IR). Primer OF dan OR menentukan spesifitas dan efisiensi pengikatan DNA template, sedangkan primer IF dan IR menentukan spesifitas pengikatan alel spesifik yang divisualisasikan dengan prosedur elektroforesis konvensional (Alyethodi *et al.,* 2016). Metode ini dipilih sebab efektif, simpel dan tergolong ekonomis dibandingkan metode *genotyping* lain. Beberapa hambatan dalam penggunaan metode T-ARMS PCR adalah prosedur optimisasi yang sulit dan dalam beberapa penelitian tidak dapat mendeteksi alel target dalam SNP *genotyping* (Ye *et al.,* 2001; Medrano and de Oliveira, 2014; Tanha *et al.,* 2015).

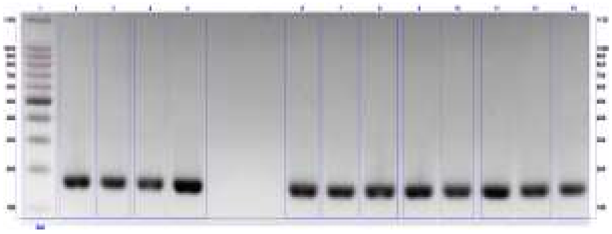

**Eksperimen 1**

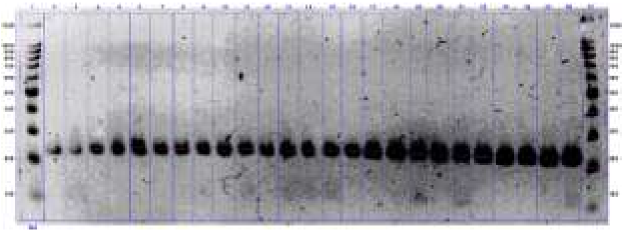

**Eksperimen 2A**

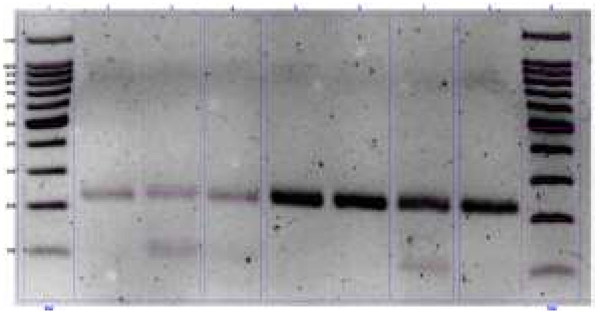

**Eksperimen 3**

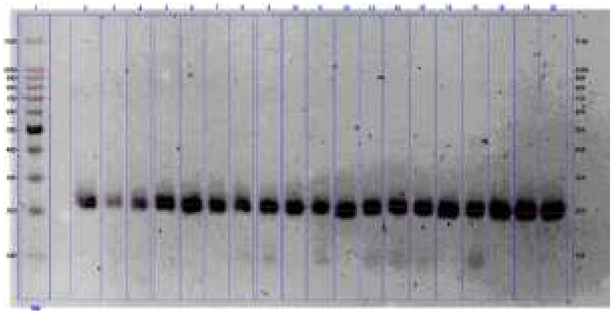

**Eksperimen 2B**

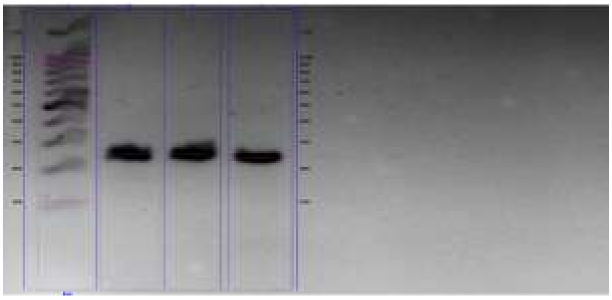

**Eksperimen 5**

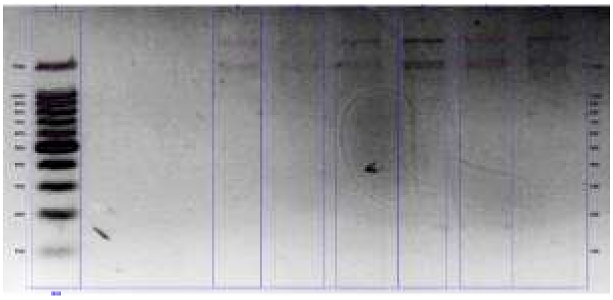

**Eksperimen 4**

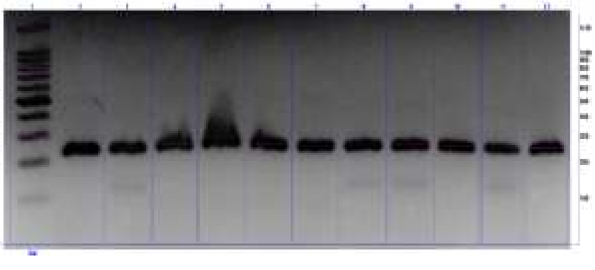

**Eksperimen 7A**

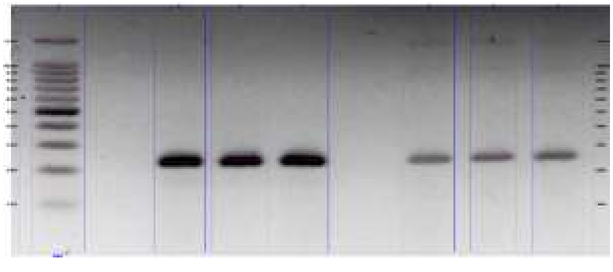

**Eksperimen 6A**

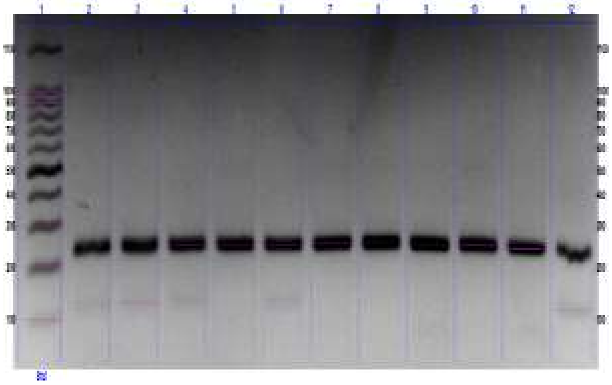

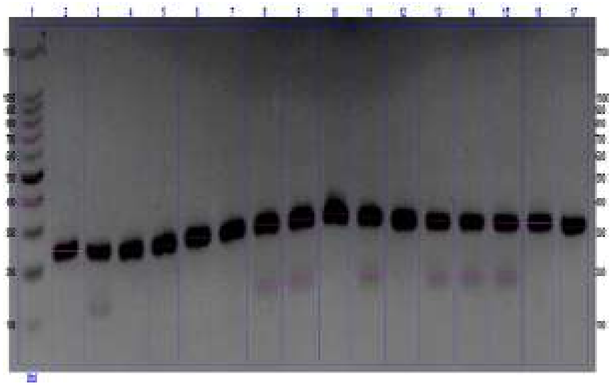

**Eksperimen 7B** **Eksperimen 6B**

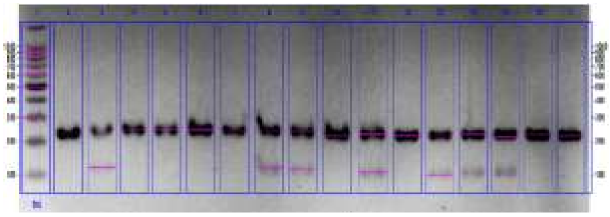

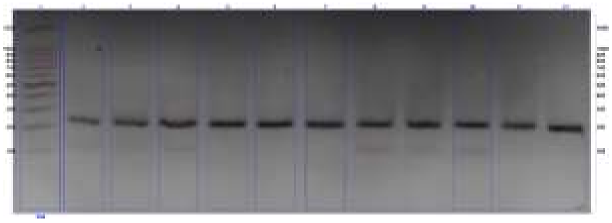

**Eksperimen 9A** **Eksperimen 8**

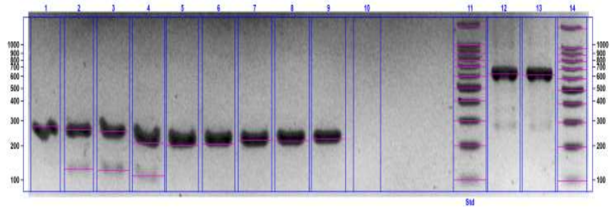

**Eksperimen 9B**

Gambar 19. Hasil eksperimen optimasi T-ARMS PCR SNPs ekson 9 *LEPR* (Dok. Pribadi, 2019).

Exp. 1: Sample tag 4 in all wells; Exp. 2.a: Sample tags 1 until 15, Bj, Ly, 3p, 7p, 8p, 10p, 82b, 90b, 92b and 97b; Exp. 2.b: Sample tags 1 until 15, 82b, 90b, 92b and 97b; Exp. 3: Sample tags Bj, Ly, 10p and 90b; Exp. 4: Sample tags Bj, 10p and 90b; Exp. 5: Sample tags Bj, Ly and 10p; Exp. 6.a: Sample tags Bj, Ly, 10p; Exp. 6.b: Sample tags 1 until 15 and Bj; Exp. 7A dan 7B: Sample tags 1-11, Bj, Ly, 10p; Exp 8: Sample tags 12, 13, 14, 1, 2, 3, 8p, 7p, 3p, 10p, 90b; Exp 9A: Sample tags 1-15, Bj; Exp 9B: Sample tags Ly, 82b, 90b, 92b, 97b, 3p, 7p, 8p, 10p. Abbreviation (Experiments: Exp.).

Dalam percobaan ini dilakukan 9 kali optimisasi hingga didapatkan konfigurasi prosedur yang sesuai (Gambar 19). Berdasarkan beberapa kriteria seperti kualitas amplikon DNA hasil PCR dan akurasi ukuran amplikon hasil optimasi maka dapat disimpulkan bahwa eksprimen 9 merupakan prosedur T-ARMS PCR yang sesuai dalam *genotyping* SNPs ekson 9 *LEPR Gallus gallus gallus* (Gambar 18). Protokol spesifik deteksi SNPs ekson 9 gen *LEPR* dengan metode T-ARMS PCR dapat mendeteksi mutasi C127A *LEPR* dengan rasio IP:OP 10:1 pmol/µM, konsentrasi template DNA ayam 100 ng/µL dengan temperatur *annealing* 55,7˚C selama 30 detik.

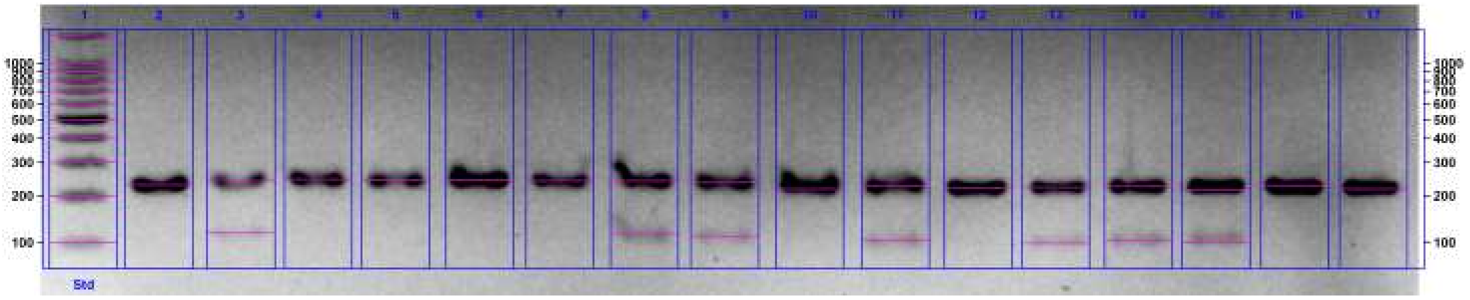

**(a)**

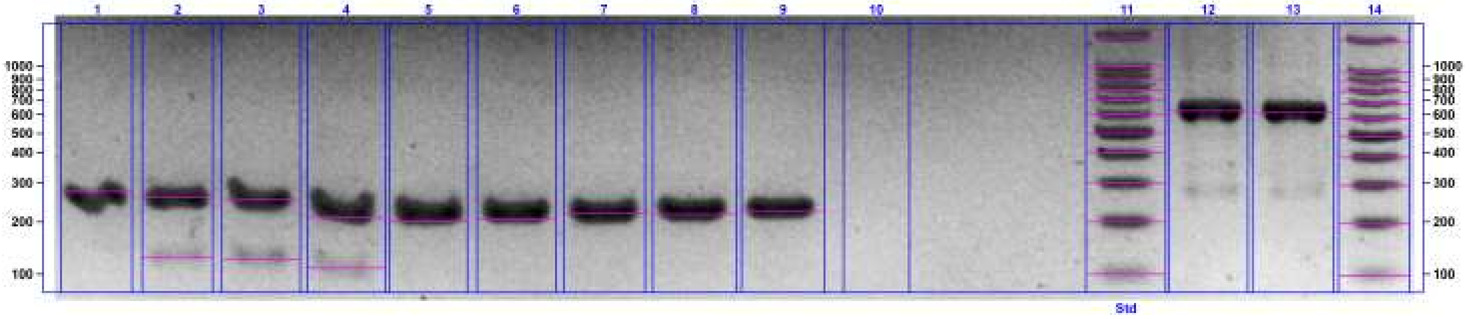

**(b)**

Gambar 20. Hasil amplifikasi SNPs ekson 9 *LEPR* ayam $F_2$ Kambro
(Dok. Pribadi, 2019)

Line 1: marker 100 bp
Line 2: OF/OR (Alel C/normal)
(a) Line 3, 8, 9, 11, 13, 14, 15: Alel A
(a) Line 2, 4, 5, 6, 7, 10, 12, 16, 17: OF/OR (Alel C/normal)
(b) Line 2, 3, 4: Alel A
(b) Line 1, 5, 6, 7, 8, 9: OF/OR (Alel C/normal)

Berdasarkan hasil *genotyping* Gambar 20 maka keberadaan mutasi transversi C127A SNPs ekson 9 *LEPR* dapat dideteksi dengan metode T-ARMS PCR. Transversi alel A *LEPR* terdeteksi pada individu ayam (a) Line 3, 8, 9, 11, 13, 14, 15 dan (b) Line 2, 3 dan 4. Penanda individu ayam tersebut secara berurutan yaitu tag 2, tag 7, tag 8, tag 10, tag 12, tag 13, tag 14, tag 82b, tag 90b dan tag 92b. Tag 2 (ChipChip3), tag 8 (ChipChip5), tag 12 (ChipChip2) dan tag 14 (ChipChip4) merupakan ayam $F_1$ Kambro. Tag 7 (Igor), tag 10 (Odin) dan tag 13 (Satrio) merupakan ayam $F_2$ Kambro. Tag 82b, tag 90b dan tag 92b merupakan ayam Broiler Cobb 500.

Individu ayam dengan alel *wild-type*/C terdeteksi pada individu ayam (a) Line 2, 4, 5, 6, 7, 10, 12, 16, 17 dan (b) Line 1, 5, 6, 7, 8 dan 9. Penanda individu ayam tersebut secara berurutan yaitu tag 1, tag 3, tag 4, tag 5, tag 6, tag 9, tag 11, tag 15, tag Bj, tag Ly, tag 97b, tag 3p, tag 7p, tag 8p dan tag 10p. Tag 1

(ChipChip) dan tag Bj (Bjorn) merupakan ayam $F_1$ Kambro. Tag 3 (Ragnar), tag 4 (Rollo), tag 5 (CaoCao), tag 6 (Clyde), tag 9 (Joy), tag 15 (Hilda) merupakan ayam $F_2$ Kambro. Tag 11 (Pelung Blirik Hitam), tag 3p, tag 7p, tag 8p dan tag 10p merupakan ayam Pelung. Tag Ly merupakan ayam Layer. Tag 97b merupakan ayam Broiler Cobb 500.

Berdasarkan hasil ini maka dapat ditemukan korelasi molekuler yang jelas terhadap fenomena yang diamati dalam parameter bobot tubuh, parameter kualitas telur, fenotip telur dan produktivitas telur. Dalam Guo *et al.* (2017) ditemukan bahwa mutasi transversi (Tv) dapat bersifat disruptif terhadap *transcription factor binding* (TF *binding*) dibandingkan mutasi transisi. Pengaruh disruptif ini berupa tidak mampunya terjadinya transkripsi beberapa asam amino yang asosiatif terhadap fungsi fisiologis. Berdasarkan hasil *genotyping* yang dilakukan maka individu *grandparent stock* (GS) Broiler Cobb 500 memiliki transversi alel A *LEPR*. Pewarisan transversi ini terwariskan kepada *parent stock* (PS) $F_1$ Kambro betina yang pewarisannya dapat dideteksi pada generasi $F_2$ Kambro.

Dari 5 sampel DNA ayam betina PS $F_1$ Kambro 80% mengalami transversi alel A pada *LEPR*. Pada ayam $F_2$ Kambro dari 15 sampel DNA terdeteksi 20% mengalami transversi alel A *LEPR* dengan jenis kelamin jantan. Pada generasi GS ayam betina Broiler Cobb 500 dari 4 sampel DNA terdeteksi 75% mengalami transversi alel A *LEPR*. Pada kelompok Pelung (n = 5 sampel DNA), Layer (n = 1 sampel DNA) dan generasi ayam $F_2$ Kambro (n = 12 sampel DNA) tidak terdeteksi adanya transversi alel A. Mutasi transversi pada *LEPR* bersifat vital sebab leptin berasosiasi dengan pertambahan bobot ayam dan produktivitas telur. Abbasi *et al.* (2011) mengungkapkan bahwa polimorfisme *LEPR* pada ayam Mazandaran dengan metode *Restriction Fragment Length Polymorphism* (RFLP) mengindikasikan dominansi frekuensi alel A. Dapat disimpulkan bahwa pewarisan mutasi transversi alel A *LEPR* terjadi pada individu betina GS Broiler Cobb 500 dan individu betina PS $F_1$ Kambro. Metode seleksi molekuler dengan T-ARMS PCR dapat digunakan dalam menseleksi parental generasi $F_3$ Kambro. Pengaruh transversi dan *inbreeding depression* mempengaruhi capaian parameter fenotip kualitatif, bobot tubuh, parameter kualitas telur eksterior, produktivitas telur dan fenotip telur pada ayam $F_2$ Kambro.

### BAB V
### SIMPULAN DAN SARAN

#### A. Simpulan

Berdasarkan penelitian yang telah dilakukan maka dapat disimpulkan bahwa:

1. Parameter fenotip kualitatif menunjukkan enam kelompok variasi fenotip tersegregasi dibandingkan ayam $F_1$ Kambro. Pertumbuhan bobot ayam $F_2$ Kambro mencapai 753,36 ± 155,31 gram dalam 7 minggu tidak signifikan terhadap ayam $F_1$ Kambro disebabkan adanya *inbreeding depression* (Fx = 25%, LI = 4,925%) dan mutasi transversi alel A *LEPR.*
2. Protokol spesifik deteksi SNPs ekson 9 gen *LEPR* dengan metode T-ARMS PCR dapat mendeteksi mutasi C127A *LEPR* dengan rasio IP:OP 10:1 pmol/µM, konsentrasi template DNA ayam 100 ng/µL dengan temperatur *annealing* 55,7˚C/30s.
3. Mutasi transversi alel A SNPs ekson 9 *LEPR* terdeteksi pada sampel DNA ayam betina $F_1$ Kambro (80%), ayam jantan $F_2$ Kambro (20%), ayam betina Broiler Cobb 500 (75%). Mutasi tersebut tidak terdeteksi pada sampel DNA ayam Layer, ayam Pelung Blirik Hitam dan ayam $F_1$ Pelung.

#### B. Saran

Saran untuk penelitian selanjutnya sebagai berikut:

1. Seleksi indukan dengan protokol spesifik deteksi *LEPR Gallus gallus* metode T-ARMS PCR untuk meningkatkan efektivitas persilangan terkait beberapa karakter pendukung produktivitas telur dan pertambahan bobot tubuh ayam.
2. Pemilihan indukan ayam $F_2$ Kambro hanya dari kelompok betina dengan alel *wild-type LEPR* dan jantan indukan merupakan galur ayam Pelung *outbreeding/crossbreeding* untuk menurunkan laju *inbreeding.*
3. Parameter vitalitas, rasio femur-tibia, analisis parameter kualitas interior telur ayam dapat diaplikasikan guna meningkatkan akurasi seleksi indukan.

### DAFTAR PUSTAKA

Abanikannda, OTF and Leigh, AO. 2012. Chicken age and egg morphometric measures on eggshell thickness. *Archiva Zootechnica*. 15(1): 61-68

Abbasi HA, Gharavysi S and Abdullahpour R. 2011. Genetic polymorphism exon 9-11 at the Leptin gene receptor in breeder hens of Mazandaran native fowls. *J. Anim. Vet. Adv*. 10(21): 2802-2805

Adachi H, Murase D and Ohkubo T. 2013. Inhibitory mechanism of signal transduction through chicken leptin receptor by suppressor of cytokine signaling 3 (SOCS3). *Jpn. Poult. Sci*. 50:262–269

Adachi H, Murase D, Atomura S, and Ohkubo T. 2012. Detection of Leptin Activity in Living Cells Expressing Chicken Leptin Receptor and STAT3. *J. Poult. Sci*. 49: 46-50

Ali M, Hossain MS, Akter S, Khan MAHNA and Hossain MM. 2013. Pathogenesis of infectious coryza in chickens (*Gallus gallus*) by Avibacterium paragallinarumn isolate of Bangladesh. *The Agriculturist*. 11(1): 39-46

Alyethodi RR, Singh U, Kumar S, Deb R, Alex R, Sharma S, Sengar GS and Prakash B. 2016. Development of a fast and economical genotyping protocol for bovine leukocyte adhesion deficiency (BLAD) in cattle. *SpringerPlus*. 5: 1442

Amills M, Jimenez N, Villalba D, Tor M, Molina E, Cubilo D, Marcos C, Francesch A, Sanchez A and Estany J. 2003. Identification of three single nucleotide polymorphisms in the chicken insulin-like growth factor 1 and 2 genes and their associations with growth and feeding traits. *Poult Sci*. 82:1485–1493

Anang A, Indrijani H dan Sundara TA. 2007. Model Matematika Kurva Produksi Telur Ayam Broiler Breeder Parent Stock *(The Mathematical Models For Egg Production Curve In Broiler Breeder Parent Stock*). *Jurnal Ilmu Ternak.* 7(1): 6-11

Anggitasari S, Sjofjan O and Djunaidi IH. 2016. Pengaruh beberapa jenis pakan komersial terhadap kinerja produksi kuantitatif dan kualitatif ayam pedaging. *Buletin Peternakan*. 40(3): 187-196

Ashwell CM, Czerwinski SM, Brocht DM and McMurtry JP. 1999. Hormonal regulation of leptin expression in *Broiler* chickens. *Am. J. Physiol*. 276:R226–R232

Assan N. 2015. Methodology and factors influencing the association of body weight, performance parameters with linear body measurements assessment in poultry. *Sci. J. Pure Appl. Sci*. 4(10): 200- 210

Bai H, Zhu J, Sun Y, Liu R, Liu N, Li D, Wen J and Chen J. 2014. Identification of genes related to beak deformity of chicken using digital gene expression profiling. *PLoS ONE*. 9(9): e107050

Bamidele O, As P Van and Elferink MG. 2012. Molecular Characterization of the Leptin Receptor Gene as a Candidate Gene in the Pulmonary Hypertension Syndrome in *Broiler* Chickens. *Pak. J. Biol. Sci*. 15:24:1187-1190

Belitz HD, Grosch W and Schieberle P. 2009. Eggs. *Food Chemistry*. p: 546-561

Cheng HW. 2010. Breeding of tomorrow's chickens to improve well-being. *Poult. Sci*. 89: 805-813

Dai HC, Long LQ, Zhang XW, Zhang WM and Wu XX. 2007. Cloning and Expression of the Duck Leptin Gene and the Effect of Leptin on Food Intake and Fatty Deposition in Mice. *Asian- Australasian Journal of Animal Sciences*. 20:6:850–55

Darwati S, Hasyim AR, Rukmiasih and Prabowo S. 2016. *Growth performance of pelung sentul kampung meat type chicken crossing on age 0-10 weeks*. Marjuki, Ridlowi A, Jaya F, Susilowati T, Wittayakun S, Bottema CDK, Alimon AR, Hsia LC, Thiruvenkadan AK, editors. Proceeding of the 3rd Animal Production International Seminar (3rd APIS) & 3rd ASEAN Regional Conference on Animal Production (3rd ARCAP). Batu (Indones): Universitas Brawijaya. p. 484-487

Darwati, S. 2000. Produktivitas Ayam Kampung, Pelung dan Resiprokalnya. *Med. Pet*. 23(2): 32- 35

Daryono BS, Roosdianto I dan Saragih HTS. 2010. Pewarisan karakter fenotip ayam hasil persilangan ayam Pelung dengan ayam Cemani. *J. Vet*. 11(4): 257-263

Das SC, Chowdhury SD, Khatun MA, Nishibori M, Isobe N and Yoshimura Y. 2008. Poultry production profile and expected future projection in Bangladesh. *World's Poult. Sci. Assoc*. 64: 99-118

de Verdal H, Narcy A, Bastianelli D, Chapuis H, Même N, Urvoix S, Le Biham-Duval E and Mignon-Grasteau S. 2011. Improving the efficiency of feed utilization in poultry by selection. 1. Genetic parameters of anatomy of the gastro-intestinal tract and digestive efficiency. *BMC Genet*. 12: 59

Depison. 2009. Karakteristik Kuantitatif dan Kualitatif Hasil Persilangan Beberapa Ayam Lokal. *Jurnal Ilmiah Ilmu-Ilmu Peternakan*. 12(1): 7-13

Diwyanto K dan Prijono SN. 2007. *Keanekaragaman sumber daya hayati ayam lokal Indonesia: manfaat dan potensi*. Bogor (Indones): Lembaga Ilmu Pengetahuan Indonesia

Duguma R. 2006. Phenotypic characterization of some indigenous chicken ecotypes of Ethiopia. *Livestock Res. Rural Dev*. 18(9)

Duman M, Şekeroğlu A, Yıldırım A, Eleroğlu H and Camcı Ö. 2016. Relation between egg shape index and egg quality characteristics. *Europ. Poult.Sci*. 80: 1-9

Dunn IC, Boswell T, Friedman-Einat M, Eshdat Y, Burt DW and Paton IR. 2000. Mapping of the leptin receptor gene (*LEPR*) to chicken chromosome 8. *Anim. Genet*. 31:290

Dunn IC, Girishvarma G, Talbot RT, Waddington D, Boswell T and Sharp PJ. 2001. Evidence for low homology between the chicken and mammalian leptin genes. *In: Avian Endocrinology* (Dawson A and Chaturvedi CM eds.). p. 327-336. Narosa Publishing House. New Delhi

El Ghany FAA, El Dein A, Soliman MM, Reza AM and El Sodany SM. 2011. Relationships between some body measurements and fertility in males of two local strains of chicken. *Egypt Poult. Sci*. 31:331–349

Eldik P. van, van der Waaij EH, Ducro B, Kooper AW, Stout TAE and Colenbrander B. 2006. Possible Negative Effects of Inbreeding on Semen Quality in Shetland Pony Stallions. *Theriogenology*. 65: 1159-1170

El-Moujahid EM, Chen S, Jin S and Lu Y. 2014. Association of leptin receptor gene polymorphisms with growth and feed efficiency in meat-type chickens. *Poult. Sci*. 93:1910–1915

Ernanto AR, Afifah D, Lesmana I and Daryono, BS. 2018. Isolation of DNA from chicken (*Gallus gallus domesticus* Linnaeus, 1758) feather with lysis buffer-phenol chloroform isoamyl alcohol method (PCI) and chelex method. Proceeding of 5th International Conference on Biological Sciences. Yogyakarta (Indones): Universitas Gadjah Mada. p. 1-5

Ernanto AR. 2017. Asosiasi Polimorfisme Gen *PRL* dan *IGF-1* terhadap Produktivitas Telur Ayam (*Gallus gallus domesticus* Linnaeus, 1758) $F_1$ Hasil Persilangan Ayam Pelung dan Layer. Thesis. Universitas Gadjah Mada. Yogyakarta

Fahey AG, Marchant-Forde RM and Cheng HW. 2007. Relationship between body weight and beak characteristics in one-day-old White Leghorn chicks: Its Implications for Beak Trimming. *Poult. Sci*. 86: 1312-1315

Fayeye TR, Hagan JK and Obadare AR. 2013. Morphometric traits and correlation between body weight and body size traits in Isa Brown and Ilorin ecotype chickens. *Iranian J. Appl. Anim. Sci*. 4(3): 609-614

Ferket PR and Gernat AG. 2006. Factors that affect feed intake of meat birds: A review. *Int. J. Poult. Sci*. 5:905–911

Frame DD. 2009. Molting and determining production of laying hens. DVM, Extension Poultry Specialist.2009-01pr. https://ucanr.edu/sites/poultry/files/186896.pdf

Friedman-Einat M, Boswell T, Horev G, Girishvarma G, Dunn IC, Talbot RT and Sharp PJ. 1999. The chicken leptin gene: has it been cloned?. *Gen. Comp. Endocrinol*. 115:354–363

Gebriel GM, Kalamah MA, El-Fiky AA, Ali AFA. 2009. Some factors affecting semen quality traits in norfa cocks. *Egypt Poult. Sci*. 29: 677–693

Gheyas AA, Boschiero C, Eory L, Ralph H, Kuo R, Woolliams JA and Burt DW. 2015. Functional classification of 15 million SNPs detected from diverse chicken populations. *DNA Res*. 22(3): 205–217

Gu Z, Zhao J, Li H, Meng H, Wang Q, and Zhu D. Single nucleotide polymorphism analysis in chicken leptin receptor exon 9. Animal Science Northeast Agricultural University. *unpublished*.

Guo, C, McDowell IC, Nodzenski M, Scholtens, DM, Allen AS, Lowe WL and Reddy TE. 2017. Transversions have larger regulatory effects than transitions. *BMC Genomics*. 18: 394

Habibah I. 2018. Karakter Fenotip, Koefisien *Inbreeding* dan Polimorfisme Gen c*TYR* Intron 4 Pada Ayam (*Gallus gallus gallus* Linnaeus, 1758) Hibrida Golden Kamper. Skripsi. Fakultas Biologi Universitas Gadjah Mada

Halverson MA, Skelly DK and Caccone A. 2006. Inbreeding Linked to Amphibian Survival in the Wild but Not in the Laboratory. *Journal of Heredity*. 97(5): 499-507

Hameed T, Mustafa MZ, Taj MK, Asadullah, Bajwa MA, Bukhar FA, Kiani MMT and Ahmed A. 2016. Hatchability and fertility in broiler breeder stock. *J. Chem. Biol. Phy. Sci*. 6(2): 266-274

Han JC, Qu HX, Wang JG, Chen GH, Yan YP, Zhang JL, Hu FM, You LY and Cheng YH. 2015. Comparison of the growth and mineralization of the femur, tibia, and metatarsus of broiler chicks. *Brazilian J. Poult. Sci.* 17(3): 333-340

Han J. 2014. Origin and evolution of molecular diversity of indigenous animal genetic resources. An invited plenary paper presented in the 16th AAAP Congress. Yogyakarta (Indonesia): Universitas Gadjah Mada

Hassan KMd, Kabir HMd, Sultana S, Hossen Amd and Haq MM. 2016. Management and production performance of Cobb-500 broiler parent stock under open housing system. *Asian Australas J. Biosci. Biotechnol.* 1(1): 66-72

Hasyim AR. 2015. *Performa hasil persilangan ayam kampung ras pedaging dengan pelung sentul pada umur 0-11 minggu*. (Thesis). [Bogor: (Indonesia)]: Institut Pertanian Bogor

Havenstein G, Ferket P, and Qureshi M. 2003. Growth, livability, and feed conversion of 1991 vs. 1957 *Broiler*s when fed typical 1957 and 2001 *Broiler* diets. *Poultry Science*. 82(March): 1500–1508.

Havlíček M, Nedomová Š, Simeonovová J, Severa L and Křivánek I. 2008. On The Evaluation Of Chicken Egg Shape Variability. *Acta univ. agric. et silvic. Mendel. Brun.* 56(5): 69–74

Henuk YL, Bale-Therik JF, Dewi GAK and Bailey CA. 2015. Native chickens and their production systems in Indonesia. *Khon Kaen Agr. J.* 43(2): 20-24

Henuk YL and Bailey CA. 2014. Husbandry systems for native chickens in Indonesia. Pp. 759 – 762. In: Proceedings of the 16th AAAP Animal Science Congress. Yogyakarta (Indones): Universitas Gadjah Mada

Hill WG. 2010. Understanding and using quantitative genetic variation. *Phil. Trans. R. Soc. B*. 365: 73-85

Horev G, Einat P, Aharoni T, Eshdat Y and Friedman-Einat M. 2000. Molecular cloning and properties of the chicken leptin-receptor (*CLEPR*) gene. *Mol. Cell Endocrinol*. 162:95–106

Hossner KL. 1998. Cellular, molecular and physiological aspects of leptin: Potential application in animal production. *Can. J. Anim. Sci*. 78:463–472

Ikegwu TM, Balogu VT, Balogu DO, Kolo SI and Babatunde J. 2016. Physical Properties of Hen's Egg. *Journal of Foods, Natural and Life Sciences*. 1: 16 – 23

International Chicken Genome Sequencing Consortium. 2004. Sequence and comparative analysis of the chicken genome provide unique perspectives on vertebrate evolution. *Nature* 432: 695–716

Iskandar S dan Susanti T. 2007. *Karakter dan Manfaat Ayam Pelung di Indonesia*. Balai Penelitian Ternak. Bogor. Hal: 128-136

Joller S, Bertschinger F, Kump E, Spiri A, von Rotz A, Schweizer-Gorgas D, Drogemuller C and Flury C. 2018. Crossed beaks in a local swiss chicken breed. *BMC Vet. Res*. 14: 68

Kabir, MdA, Islam MS and Dutta RK. 2012. Egg morphometric analyses in chickens and some selected birds. *Univ. j. zool. Rajshahi Univ*. (31): 85-87

Kapa Biosystems. 2016. KAPA Taq ReadyMix PCR Kit. p: 1-2

Kartasudjana R and Suprijatna E. 2010. *Manajemen ternak unggas*. Cetakan Kedua. Jakarta (Indones): Penerbit Penebar Swadaya

Kartika AA, Widayati KA, Burhanuddin, Ulfah A and Farajallah A. 2016. Eksplorasi preferensi masyarakat terhadap pemanfaatan ayam lokal di Kabupaten Bogor Jawa Barat. *Jurnal Ilmu Pertanian Indonesia (JIPI)*. 21(3): 180-185

Kerje S, Sharma P, Gunnarsson U, Kim H, Bagchi S, Fredriksson R, Schutz K, Jensen P, von Heijne G, Okimoto R and Andersson L. 2004. The dominant white, dun and smoky color variants in chicken are associated with insertion/deletion polymorphisms in the PMEL17 Gene. *Genetics*. 168: 1507-1518

Klok MD, Jakobsdottir S and Drent ML. 2007. The Role of Leptin and Ghrelin in the Regulation of Food Intake and Body Weight in Humans: A Review. *Obesity Reviews*. 8:1:21–34.

Konig S, Tsehay F, Sitzenstock F, von Borstel UU, Schmutz M, Preisinger R and Simianer H. 2010. Evaluation of inbreeding in laying hens by applying optimum genetic contribution and gene flow theory. *Poultry Science*. 89: 658-667

Landsverk ML and Wong LJC. 2013. *Next Generation Sequencing: Translation to Clinical Diagnostics*. New York. p: 19-36

Li H, Deeb N, Zhou H, Mitchell AD, Ashwell CM and Lamont SJ. 2003. Chicken Quantitative Trait Loci for Growth and Body Composition Associated with Transforming Growth Factor-β Genes. *Poultry Science*. 82: 347 - 356

Liu X, Dunn IC, Sharp PJ and Boswell T. 2007. Molecular cloning and tissue distribution of a short form chicken leptin receptor mRNA. *Domest. Anim. Endocrinol*. 32:155–166

Liyanage RP, Dematawewa CMB and Silva GLLP. 2015. Comparative Study on Morphological and Morphometric Features of Village Chicken in Sri Lanka. *Tropical Agricultural Research*. 26(2): 261 – 273

Lohmann Tierzucht. 2019. *Management Guide Layers*. Germany. p: 1-40

Lukanov H, Genchev A and Pavlov A. 2015. Colour Traits Of Chicken Eggs With Different Eggshell Pigmentation. *Trakia Journal of Sciences*. 2: 149-158

Mabelebele M, Norris D, Siwendu NA, Ng'ambi JW, Alabi OJ and Mbajiorgu CA. 2017. Bone morphometric parameters of the tibia and femur of indigenous and broiler chickens reared intensively. *Appl. Ecol. Environ. Res*. 15(4): 1387-1398

Mahardhika, IWS and Daryono BS. 2019. Kambro Chicken Phenotype Hybrid of ♀ Broiler Cobb 500 Cross ♂ Pelung Blirik Hitam. *Indonesian Journal of Animal and Veterinary. unpublished*

Mariandayani HN, Darwati S, Sutanto E and Sinaga E. 2017. Peningkatan produktivitas ayam lokal melalui persilangan tiga rumpun ayam lokal pada generasi kedua. Prosiding Seminar Nasional Biologi 2017:

Pendidikan Biologi untuk Masa Depan Bumi. Aceh (Indones): Jurusan Pendidikan Biologi, Universitas Syiah Kuala. p. 139-146

Mariandayani HN, Solihin DD, Sulandari S and Sumantri C. 2013. Keragaman fenotipik dan pendugaan jarak genetik pada ayam lokal dan ayam broiler menggunakan analisis morfologi. *J Vet*. 14(4): 475-484.

Martin A, Dunnington EA, Gross WB, Briles WE, Briles RW and Siegel PB. 1990. Production traits and allo antigen systems in lines of chickens selected for high or low antibody responses to sheep erythrocytes. *Poult. Sci*. 69:871–878

Medrano RFV and de Oliveira. 2014. Guidelines for the Tetra-Primer ARMS–PCR Technique Development. *Mol. Biotechnol*. 56:599–608

Mrázová Ľ, Angelovi M, and Král M. 2012. Monitoring the Expression of Leptin Receptor in Avian Model. *Anim. Sci. Biotech.* 45(1): 322–327

Naka T, Narazaki M, Hirata M, Matsumoto T, Minamoto S, Aono A, Nishimoto N, Kajita T, Taga T, Yoshizaki K, Akira S and Kishimoto T. 1997. Structure and function of a new STAT-induced STAT inhibitor. *Nature*. 387: 924-929

Narushin VG. 2005. Egg Geometry Calculation Using the Measurements of Length and Breadth. *Poultry Science*. 84:482–484

Nataamijaya AG. 2005. Karakteristik Penampilan Pola Warna Bulu, Kulit, Sisik Kaki, dan Paruh Ayam Pelung di Garut dan Ayam Sentul di Ciamis. *Buletin Plasma Nutfah* (11) 1: 1-5

Nataamijaya AG. 2010. Pengembangan Potensi Ayam Lokal Untuk Menunjang Peningkatan Kesejahteraan Petani. *Jurnal Litbang Pertanian*, 29(4): 131- 133

Navara KJ, Anderson EM and Edwards ML. 2012. Comb size and color relate to sperm quality: a test of the phenotype-linked fertility hypothesis. *Behav. Ecol*. 12: 1036-1041

Newton CR, Graham A, Heptinstall LE, Powell SJ, Summers C, Kalsheker N, Smith JC and Markham AF. 1989. Analysis of any point mutation in DNA. The ampliflcation refractory mutation system (ARMS). *Nucleic Acid Research*. 17(7): 2503-2516

Nie Q, Mingming L, Jianhua Q, Hua Z, Guanfu Y and Xiquan Z. 2005. Original article identification and characterization of single nucleotide polymorphisms in 12 chicken growth-correlated genes by denaturing high performance liquid chromatography. *Genet. Sel. Evol.* 37:339-360

Nietlisbach P, Keller LF, Camenish G, Guillaume F, Arcesee P, Reid JM and Postma E. 2017. Pedigree-based inbreeding coefficient explains more variation in fitness than heterozygosity at 160 microsatellites in a wild bird population. *Proc. R.Soc. B*. 284: 2016 - 2763

Ningsih R and Prabowo DW. 2017. Tingkat integrasi pasar ayam broiler di sentra produksi utama: studi kasus Jawa Timur dan Jawa Barat. *Buletin Ilmiah Litbang Perdagangan*. 11(2): 247- 270

Ninov K, Ledur MC, Alves HJ, Rosário MFdo, Nones K, and Coutinho LL. 2008. Investigation of Leptin gene in *Broiler* and *Layer* chicken lines. *Scientia Agricola*, 65(2), 214–219

Niv-Spector L, Raver N, Friedman-Einat M, Grosclaude J, Gussakovsky EE, Livnah O and Gertler A. 2005. Mapping Leptin-Interacting Sites in Recombinant Leptin-Binding Domain (LBD) Subcloned from Chicken Leptin Receptor. *Biochem. J.* 390:475–484

Nurfadillah S, Rachmina D and Kusnadi N. 2018. Impact of trade liberization on Indonesian broiler competitiveness. *J. Indonesian Trop. Anim. Agri.* 43(4): 429-437

Nurhuda SA. 2017. Pertumbuhan generasi ketiga hasil persilangan ayam lokal dengan ayam ras pedaging sampai umur 12 minggu (Thesis). [Bogor: (Indones)]: Institut Pertanian Bogor

Nuroso. 2010. *Ayam kampung pedaging hari per hari*. Jakarta (Indones): Penebar Swadaya

Nwagu BI, Olorunju SAS, Oni OO, Eduvie LO, Adeyinka IA, Sekoni AA and Abeke FO. 2007. In-Breeding Effect on Performance of Rhode Island Chickens Selected for Part-Period Egg Production. *International Journal of Poultry Science*. 6(1): 13-17

Ohkubo T and Adachi H. 2008. Leptin signaling and action in birds. *Journal of Poultry Science*. 45:269-273

Ohkubo T, Nishio M, Tsurudome M, Ito M and Ito Y. 2007. Existence of leptin receptor protein in chicken tissues: isolation of a monoclonal antibody against chicken leptin receptor. General and Comparative *Endocrinology*. 151:269-273

Ohkubo T, Tanaka M and Nakashima K. 2000. Structure and tissue distribution of chicken leptin receptor (*cOb-R*) mRNA. *Biochimica et Biophysica Acta*. 1491: 303-308

Ohkubo T. 2014. Recent Progress in Avian Leptin Research Recent Progress in Avian Leptin Research. *J. Poult. Sci.* 51:343-351

Oldenbroek K and van der Waaij L. 2014. *Textbook animal breeding: animal breeding and genetics for BSc students*. Centre for Genetic Resources (Netherlands): The Netherlands and Animal Breeding and Genomics Centre

Paczoska-Eliasiewicz HE, Gertler A, Proszkowiec M, Proudman J, Hrabia A, Sechman A and Rzasa J. 2003. Attenuation by leptin of the effects of fasting on ovarian function in hens (*Gallus domesticus*). *Reproduction.* 126:6: 739–751

Pauwels J, Coopman F, Cools A, Michiels J, Fremaut D, de Smet S and Janssens GPJ. 2015. Selection for growth performance in broiler chickens associates with less diet flexibility. *PLoS ONE*. 10(6): e0127819

Paxton H, Tickle PG, Rankin JW, Codd JR and Hutchinson JR. 2014. Anatomical and biomechanical traits of broiler chickens across ontogeny. Part II. Body segment inertial properties and muscle architecture of the pelvic limb. *Peer J.* 2: e473

Peng B, Wang Q, Luo Y, He J, Tan T, and Zhu H. 2017. A novel and quick PCR- based method to genotype mice with a leptin receptor mutation (db / db mice). *Acta Pharmacologica Sinica*. 2017: 1-7

Pinard-van der Laan MH, Siegel PB, and Lamont SJ. 1998. Lessons from selection experiments on immune response in the chicken. *Poult. Avian Biol. Rev*. 9: 125–141

Pitel F, Bergé R, Coquerelle G, Crooijmans RPMA, Groenen MAM, Vignal A, and Tixier-Boichard M. 2000. Mapping the Naked Neck (NA) and Polydactyly (PO) mutants of the chicken with microsatellite molecular markers. *Genet. Sel Evol*. 32: 73–86

Pratama A, Suradi K, Balia RL, Chairunnisa H, Sutardjo DS, Suryaningsih L dan Putranto S. 2015. Evaluasi Karakteristik Sifat Fisik Karkas Ayam Broiler Berdasarkan Bobot Badan Hidup (*Evaluation of physical characteristics of broiler carcasses based on live*). *Jurnal Ilmu Ternak*. 15(2): 61–64.

Rahman MR, Chowdhury SD, Hossain ME and Ahammed M. 2015. Growth and early laying performance of a broiler parent stock in an open-sided house under restricted feeding. *Bangladesh J. Anim. Sci*. 44(1): 40-45

Richards MP and Poch SM. 2003. Molecular cloning and expression of the turkey leptin receptor gene. Comparative Biochemistry and Physiology. *Biochemistry and Molecular Biology*. 136: 833-847

Sartika T, Sopiyana S and Iskandar S. 2010. Performa ayam sentul koleksi ex-situ di balai peneltian ternak. Pengembangan peternakan berkelanjutan: sistem produksi berbasis ekosistem lokal. Hernaman I, Tanuwira UH, Lengkey HAW, Yumiati H, Sulistyati M, Hidayati YA, Herlina L, Indrijani H, Sujana E, Putranto WS, Islami RZ, Widiawati Y, Sofyan O, Syamsu JA, penyunting. Prosiding Seminar Nasional Peternakan Berkelanjutan ke-2. Jatinangor (Indones): Fakultas Peternakan Universitas Padjajaran. p. 39-51

Sartika T, Sulandari S, Zein MSA and Paryati S. 2016. Mengangkat potensi genetik dan produktivitas ayam gaok. Kementerian Pertanian RI, penyunting. Prosiding Lokakarya Nasional Pengelolaan dan Perlindungan Sumber Daya Genetik di Indonesia: Manfaat Ekonomi untuk Mewujudkan Ketahanan Nasional. Bogor (Indones): Badan Litbang Pertanian. p. 251-256

Sawitri R dan Takandjandji M. 2012. Inbreeding pada Populasi Banteng (*Bos javanicus*, d'Alton 1832) di Kebun Binatang Surabaya. *Buletin Plasma Nutfah*. 18(2): 84-94

Schmid M, Nanda I, Guttenbach M, Steinlein C, Hoehn M, Schartl M and Mizuno S. 2000. First report on chicken genes and chromosomes 2000. *Cytogenetic and Genome Research*. 90(3–4): 169–218

Schmid M, Nanda I, Hoehn H, Schartl M, Haaf T, Buerstedd JM and Mizuno S. 2015. Third Report on Chicken Genes and Chromosomes. *BioR*. 14: 1–19

Schmid M, Nanda I, Hoehn H, Schartl M, Haaf T, Buerstedde JM and Mizuno S. 2005. Second report on chicken genes and chromosomes 2005. *Cytogenetic and Genome Research*. 109(4): 415–479

Schwartz MW, Seeley RJ, woods SC, Weigle DS, Campfield LA, Burn P and Baskin DG. 1997. Leptin increase hypothalamic pro-opiomelanocortin mRNA expression in the rostral arcuate nucleus. *Diabetes*. 46: 2119 - 2123

Sekeroğlu A, Kayaalp GT and Sarica M. 2000: The Regression and correlation analysis on egg parameters in Denizli poultry. *Journal of Agricultural Faculty, Cukurova University*. 15: 69-74

Semakula J, Lusembo P, Kugonza DR, Mutetikka D, Ssennyonjo J and Mwesigwa M. 2011. Estimation of live body weight using zoometrical measurements for improved marketing of indigenous chicken in the Lake Victoria basin of Uganda. *Livestock Res. Rural Dev.* 23(8)

Seroussi E, Cinnamon Y, Yosefi S, Genin O, Smith JG, Rafati N and Friedman-Einat M. 2016. Identification of the long-sought leptin in chicken and duck: Expression pattern of the highly GC-rich avian leptin fits an autocrine/paracrine rather than endocrine function. *Endocrinology*. 157(2): 737–751

Seroussi E, Pitel F, Leroux S, Morisson M, Bornelöv S, Miyara S and Friedman-Einat M. 2017. Correction: Mapping of leptin and its syntenic genes to chicken chromosome 1p. *BMC Genetics*. 18(1): 1–8

Setiawati T, Afnan R dan Ulupi N. 2016. Performa Produksi dan Kualitas Telur Ayam Petelur pada Sistem Litter dan Cage dengan Suhu Kandang Berbeda. *Jurnal Ilmu Produksi dan Teknologi Hasil Peternakan*. 4(1): 197-203

Sewalem A, Johansson K, Wilhelmson M and Lippers K. 1999. Inbreeding and inbreeding depression on reproduction and production traits of White leghorn lines selected for egg production traits. *J. Dairy Sci*. 40: 203 - 208

Shad AGK, Zalani AM and Nasr J. 2013. Estimation of Genetic Parameters, Inbreeding Trend and its Effects on Production and Reproduction Traits of Native Fowls in Fars Province. *Pakistan Journal of Biological Sciences*. 16(12): 598-600

Shim MY, Karnuah AB, Mitchell AD, Anthony NB, Pesti GM and Anggrey SE. 2012. The effects of growth rate on leg morphology and tibia breaking strength, mineral density, mineral content, and bone ash in broilers. *Poult. Sci*. 91: 1790-1795

Sogindor BA. 2017. Performa pertumbuhan hasil persilangan ayam lokal dengan ayam ras pedaging umur 1 sampai 12 minggu (Thesis). [Bogor: (Indones)]: Institut Pertanian Bogor

Solikin T, Tanwiriah W dan Asmara IY. 2016. Bobot akhir, bobot karkas, dan income over feed and chick cost ayam sentul Barokah Abadi Farm Ciamis. *Student e-Journal Fakultas Peternakan Universitas Padjadjaran*. 5(4): 1-9.

Starr R, Willson TA, Viney EM, Murray LJ, Rayner JR, Jenkins BJ, Gonda TJ, Alexander WS, Metcalf D, Nicola NA and Hilton DJ. 1997. A family of cytokine-inducible inhibitors of signalling. *Nature*. 387: 917-921.

Statistik Peternakan dan Kesehatan Hewan. 2017. Jakarta (Indones): Direktorat Jenderal Peternakan dan Kesehatan Hewan Kementerian Pertanian RI. p. 91-92

Sudrajat dan Isyanto AY. 2018. Keragaan peternakan ayam sentul di Kabupaten Ciamis. *Jurnal Pemikiran Masyarakat Ilmiah Berwawasan Agribisnis*. 4(2): 237-253

Sun Y, Zhao G, Liu R, Zheng M, Hu Y, Wu D, Zhang L, Li P and Wen J. 2013. The identification of 14 new genes for meat quality traits in chicken using a genome-wide association study. *BMC Genomics*. 14: 458

Suprijatna E. 2010. Strategi pengembangan ayam lokal berbasis sumber daya lokal dan berwawasan lingkungan. Seminar Nasional Unggas Lokal ke IV. Sunarti D, Suprijatna E, Mahfudz LD, Sarengat W, Karno, Nuswantara LK, Surono, Sarjana TA, penyunting. Bogor (Indones): Fakultas Peternakan Universitas Diponegoro. p. 55-88

Suwandi. 2015. Outlook komoditas pertanian subsektor peternakan daging ayam. Jakarta (Indones): Pusat Data dan Sistem Informasi Pertanian Sekretariat Jenderal Kementerian Pertanian. p. 8-27

Szwaczkowski T, Katarzyna Cywa-Benko and Stanislaw W. 2003. A note on inbreeding effects on productive and reproductive traits in laying hens. *Animal Science Papers and Reports*. 21: 121-129

Tamzil MH, Lestari L and Indarsih B. 2018. Measurement of several qualitative traits and body size of Lombok Muscovy ducks (Cairina moshcata) in semi-intensive rearing. *J. Indonesian Trop. Anim. Sci*. 43(4): 333-342

Tanha HM, Mojtabavi NM, Rahgozar S, Rasa SM and Vallian S. 2015. Modified tetra-primer ARMS PCR as a single-nucleotide polymorphism genotyping tool. *Genet. Test Mol. Biomarkers.* 19(3): 156 - 161

Taouis M, Chen JW, Daviaud C, Dupont J, Derouet M and Simon J. 1998. Cloning the chicken leptin gene. *Gene*. 208: 239-242

Telalbasic R, Baban M and Rahmanovic A. 2007. Inbreeding. *Biotechnology in Animal Husbandry*. 23(5-6): 113 - 130

Udeh I, Ugwu SOC and Ogagifo NL. 2011. Predicting semen traits of local and exotic cocks using linear body measurements. *Asian J. Anim. Sci*. 5: 268–276

Ukwu HO, Okoro VMO and Nosike RJ. 2014. Statistical modelling of body weight and linear body measurements in Nigerian indigenous chicken. *IOSR J. Agri. Vet. Sci*. 7(1): 27-30

van der Pol CW, Molenaar R, Buitink CJ, van Roovert-Reijrink IAM, Maatjens CM, van den Brand H and Kemp B. 2015. Lighting schedule and dimming period in early life: consequences for broiler chicken leg bone development. *Poult. Sci*. 94: 2980-2988

Wakchaure R and Ganguly S. 2015. Inbreeding, its Effects and Applications in Animal Genetics and Breeding: A Review. *International Journal of Emerging Technology and Advanced Engineering*. 5(9): 73-76

Walpole RE. 1995. *Pengantar statistika*. Jakarta (Indones): Gramedia Pustaka Umum

Wang F, Lu L, Yuan H, Tian Y, Li J, Shen J, Tao Z and Fu Y. 2011. Molecular cloning, expression, and regulation of goose leptin receptor gene in adipocytes. *Molecular and Cellular Biochemistry*. 353: 267-274

Wang Y, Li H, Zhang Y, Gu Z, Li Z and Wang Q. 2006. Analysis on Association of a SNP in the Chicken *OBR* Gene with Growth and Body Composition Traits. *Asian-Aust J. Anim. Sci*. 19(12): 1706–1710.

Waranusast R, Intayod P and Makhod D. 2017. Egg Size Classification on Android Mobile Devices Using Image Processing and Machine Learning. Department of Electrical and Computer Engineering Faculty of Engineering, Naresuan University, Phitsanulok, Thailand.

Wilson, D. 2010. *Poultry a guide to anatomy and selected species*. Illinois (USA): ITCS Instructional Materials University of Illinois. p. 3-4

Wolc A, Zhao HH, Arango J, Settar P, Fulton JE, O'Sullivan NP, Preisinger R, Stricker C, Habier D, Fernando RL, Garrick DJ, Lamont SJ and Dekker JCM. 2015. Response and inbreeding from a genomic selection experiment in layer chickens. *Genomic Selection Evolution*. 47: 59

Xu J, Lin S, Gao X, Nie Q, Luo Q and Zhang X. 2017. Mapping of id locus for dermal shank melanin in a chinese indigenous chicken breed. *J. Gen*. 96(6): 977-983

Yakubu A, Kuje D and Okpeku M. 2009. Principal components as measures of size and shape in Nigerian indigenous chickens. *Thai. J. Agri. Sci*. 42(3): 167-176

Ye S, Humphries S and Green F. 1992. Allele specific amplification by tetra-primer PCR. *Nucleic Acid Ressearch.* 20(5): 1152

Ye S, Dhillon S, Ke X, Collins AR, and Day INM. 2001. An efficient procedure for genotyping single nucleotide polymorphisms. *Nucleic Acid Res.,* 29(17)

Yerturk M, Avci M and Bozkaya F. 2008. Effects of closed breeding on some reproductive performance of a small Japanese quail flock in Sanliurfa. *J.Anim. Vet. Adv.* 7: 963 - 967

Zhang W and Aggrey SE. 2003. Genetic variation in feed utilization efficiency of meat-type chickens. *World's Poult. Sci. J.* 59: 328–339

Zhang Y, Proenca R, Maffei M, Barone M, Leopold L and Friedman JM. 1994. Positional cloning of the mouse obese gene and its human homologue. *Nature*. 372: 425–431

Zhou P, Zheng W, Zhao C, Shen C and Sun G, 2009, in IFIP International Federation for Information Processing, Volume 295, Computer and Computing Technologies in Agriculture II, Volume 3, eds. D. Li, Z. Chunjiang, (Boston: Springer). pp: 1647–1653
